# A combinatorial domain screening platform reveals epigenetic effector interactions for transcriptional perturbation

**DOI:** 10.1101/2024.10.28.620683

**Authors:** Hyungseok C. Moon, Michael H. Herschl, April Pawluk, Silvana Konermann, Patrick D. Hsu

## Abstract

Epigenetic regulation involves the coordinated interplay of diverse proteins. To systematically explore these combinations, we present COMBINE (combinatorial interaction exploration), a high-throughput platform that tests over 50,000 pairs of epigenetic effector domains up to 2,094 amino acids in length for their ability to modulate endogenous human gene transcription. COMBINE revealed diverse synergistic and antagonistic interactions between epigenetic protein domains, including a potent KRAB-L3MBTL3 fusion that enhanced gene silencing up to 34-fold in dose-limited conditions and enabled robust bidirectional CRISPR perturbation. Inducible screening showed DNA methylation modifiers are essential for epigenetic memory, with distinct combinations driving long-term silencing, repression, or activation. Notably, we identified TET1-based combinations that induce hit-and-run upregulation for over 50 days, demonstrating long-term transcriptional activation. This systematic analysis of pairwise domain interactions provides a rich resource for understanding epigenetic crosstalk and developing next-generation epigenome editing tools. More broadly, COMBINE offers a generalizable platform to functionally characterize combinatorial biological processes at scale.

## Introduction

Epigenetic regulation in mammalian cells is a highly coordinated process, essential for organizing genetic information and orchestrating pivotal cellular functions^1^ like gene regulation, differentiation, and DNA replication^2^ or repair^3^. Central to this regulation are large, multi-protein complexes that control epigenetic states through intricate combinations of post-translational modifications of DNA-scaffolding histone proteins and epigenetic modifications of the genome itself^4^. These complexes often contain numerous subunits with distinct functionalities^5,6^, such as the Polycomb repressive complexes^7^, that coordinate a complex code of different histone modifications to enable context-dependent control of gene expression and chromatin state.

Inspired by these natural mechanisms, the epigenome editing toolbox leverages similar combinatorial concepts that bring together programmable DNA-binding domains with epigenetic effector proteins^8,9^. For maximal activity, transcriptional activators such as the <u>s</u>ynergistic <u>a</u>ctivation <u>m</u>ediator (SAM) complex fuse a viral activation domain to dCas9 in addition to aptamer-based recruitment of two other activators^10^. Epigenetic silencers such as CRISPRoff or EvoETR have also benefited from combining DNA methyltransferase domains with Krüppel associated box (KRAB) repressors, enabling long-term gene silencing with hit-and-run delivery of the editor system^11–13^.

To date, the development of epigenetic editors has largely relied on rational design or low-throughput, guess-and-test approaches^10–23^. Recent advances in pooled gene synthesis have enabled the functional screening of short peptide libraries of ∼80 amino acids tiling larger effector protein candidates^24–27^, further allowing the annotation of novel bivalent interactions that are synergistic or antagonistic^28^. However, the synthesis length restriction of current-generation oligonucleotide pools may not accommodate the size of functionally active elements, and the use of engineered reporter loci may not extrapolate to endogenous genetic contexts. One recent study used barcoded multicistronic adaptors to clone and screen pairs of transcription factors up to 5.8 kilobases in length using a T cell knock-in system, but the generalizability and scale of this approach is limited by the requirement of a cell-type specific functionally monoallelic target locus and a sharp length-dependent integration bias^29^.

To address these challenges, we sought to develop a generalizable high-throughput screening platform to functionally characterize combinations of long and heterogenous epigenetic effectors, with the goal of 1) identifying new effector combinations that increase the potency or durability of transcriptional perturbations, and 2) uncovering previously unknown interactions between individual effector domains.

Here, we report COMBINE (<u>comb</u>inatorial <u>in</u>teraction <u>e</u>xploration), a high-throughput and inducible screening platform that can accommodate heterogeneously-sized fusion elements, including large catalytic domains up to 6.3 kilobases in length. We applied COMBINE to measure the transcriptional perturbation landscape of an endogenous human gene with over 50,000 bivalent epigenetic effectors, composed of pairs of protein domains from diverse classes including readers, recruiters, structural factors, and catalytic writers or erasers.

More than 6,500 bivalent epigenetic effectors were identified as significant hits that perturb the transcription of an endogenous target gene in human cells. Systematic analysis and validation of combinatorial interactions enabled the discovery of previously unknown synergistic or antagonistic interdomain interactions that control epigenetic regulation of transcription in human cells. Our screening data allowed us to nominate new epigenetic effector domain combinations for synthetic biology and genetic medicine, including a potent KRAB-L3MBTL3 fusion that enhanced gene silencing up to 34-fold in dose-limited conditions and enabled a robust bidirectional CRISPR perturbation system to simultaneously activate one target gene and silence another. Examining durable long-term perturbations in our screen, we further identified novel combinations containing CpG DNA methylation-related domains that drive heritable silencing, repression, or activation of an endogenous target gene. COMBINE should be generalizable to a wide variety of lentivirus-transducible cell types and protein domain classes, as well as DNA and RNA elements, providing a broad platform to interrogate biological interactions at scale.

## Results

### Combinatorial interaction exploration (COMBINE) enables high-throughput bivalent domain screening

Considering the combinatorial nature of epigenetic regulation (**Fig. 1a**), we sought to assess the functional effects of epigenetic domain pairs on gene expression to uncover new biology and potential domain combinations for epigenome editing tools. The limitations of currently available bivalent domain screening technologies necessitated the development of a new pairwise domain screening approach that could accommodate larger domain sizes (many over 1,000 amino acids in length) to assess combinations of epigenetic reader, recruiter, writer, and eraser domains for their effects on gene expression in a high-throughput pooled screen.

**Figure 1.**
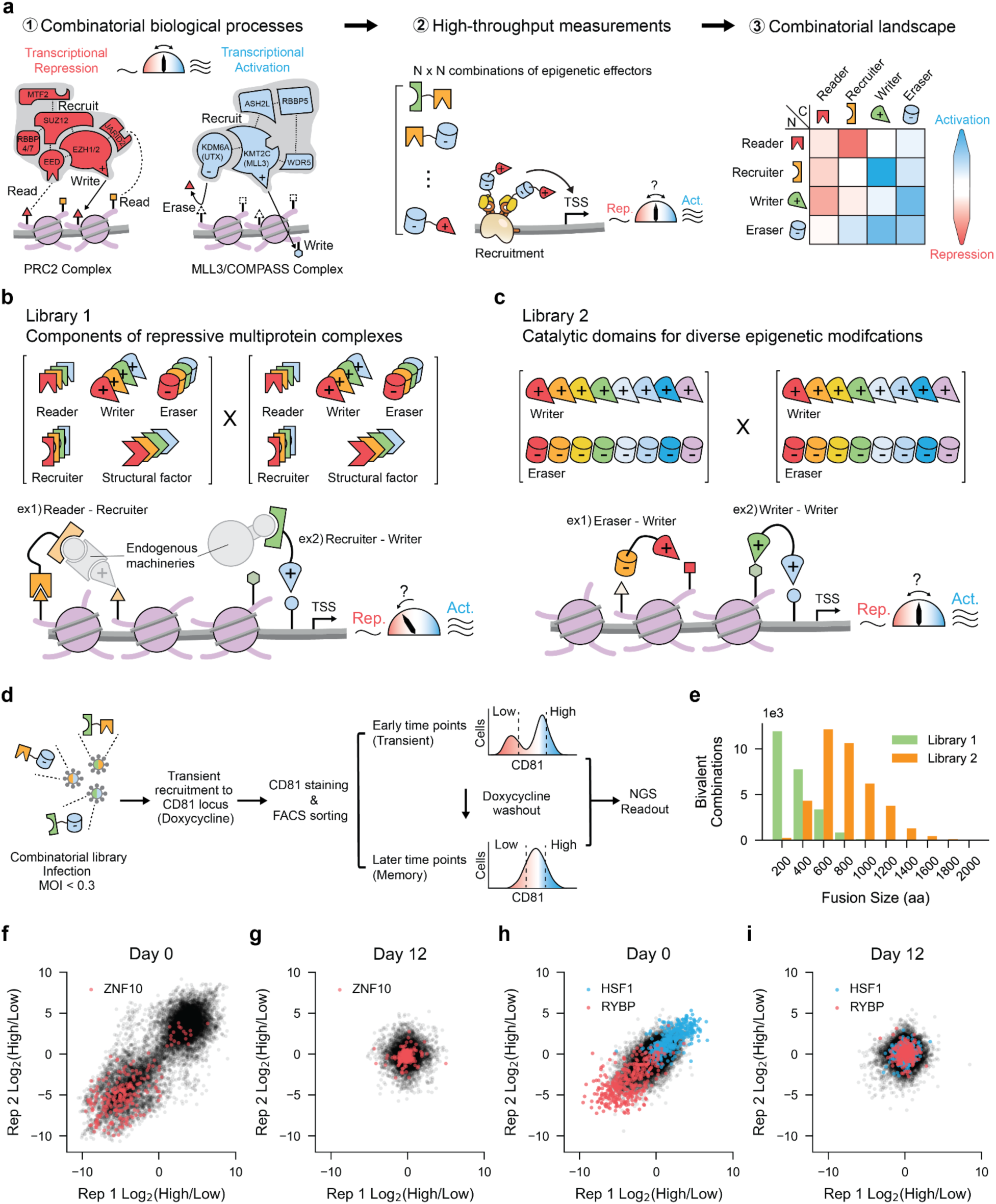
COMBINE discovers novel epigenetic effector pairs that regulate endogenous human transcription. (**a**) Conceptual framework for interrogating combinatorial biological processes through HTS of an N x N combinatorial domain library. Schematics of representative multi-protein complexes, including the Polycomb repressive complex 2 (PRC2) and the activating MLL3/COMPASS complex, are shown. N x N combinatorial domains are recruited to the endogenous target gene loci by a dCas9/MS2 system. Subpopulations of cells are enriched based on the target gene expression level in a high-throughput manner. Following the NGS readout, the combinatorial landscape of transcriptional perturbation is analyzed. (**b**) Schematic of Library 1 composed of epigenetic readers, recruiters, structural factors, and writers/erasers curated from diverse multiprotein complexes with an established ability to mediate transcriptional repression. Two examples of possible domain pairs from Library 1 are shown. A full list of members is available in **Supplementary Table 1**. (**c**) Schematic of the composition of Library 2 including direct writers and erasers of diverse epigenetic modifications, curated agnostic of their known effect on transcription. Two examples of bivalent domain candidates from Library 2 are shown. A full list of members is available in **Supplementary Table 2**. (**d**) Experimental procedure for the combinatorial HTS of epigenetic effector pairs. Combinatorial domain candidates are recruited to the endogenous CD81 locus for 5-6 days with the addition of doxycycline. After staining with anti-CD81 antibody, cells are sorted into two bins (High and Low) based on the CD81 expression level. NGS is performed to measure the enrichment of bivalent domain candidates within sorted populations. Multiple timepoints are taken following the doxycycline removal – Day 0, Day 6, and Day 12 for Library 1 and Day 0 and Day 12 for Library 2. For more detailed descriptions, refer to methods and **Supplementary Figs. 5 and 6**. (**e**) Theoretical size distribution of bivalent domain candidates from Library 1 and 2 assuming uniform coverage. (**f**) Scatter plot of enrichment scores from Library 1 at Day 0. Effectors containing the repressive ZNF10 KRAB domain are highlighted in red. (**g**) Scatter plot of enrichment scores from Library 1 at Day 12. Effectors containing the repressive ZNF10 KRAB domain are highlighted in red. (**h**) Scatter plot of enrichment scores from Library 2 at Day 0. Effectors containing the repressive RYBP domain are highlighted in red. Effectors containing the HSF1 activator are highlighted in blue. (**i**) Scatter plot of enrichment scores from Library 2 at Day 12. Effectors containing the repressive RYBP domain are highlighted in red. Effectors containing the HSF1 activator are highlighted in blue.

To achieve this, we first curated two complementary libraries of epigenetic effectors, each with distinct design principles. Library 1 was designed to identify combinatorial epigenetic repressors and is composed of readers, recruiters, structural factors, writers, and erasers from known repressive epigenetic multiprotein complexes (**Fig. 1b** and **Supplementary Table 1**). It contains a total of 155 domains ranging from 80 aa to 545 aa (**Supplementary Fig. 1a,b**). Library 2 was designed to investigate how diverse types of direct epigenetic modifications interact to modulate transcription in both directions (repression or activation). This library is exclusively composed of catalytic writers and erasers of histone and DNA modifications, irrespective of their known effects on transcription (**Fig. 1c** and **Supplementary Table 2**). It contains 198 domains, with a wider size distribution from 100 aa to 1,036 aa (**Supplementary Fig. 1c,d**). To mitigate length bias in lentivirus generation, transduction^30^, and NGS readout^31^, we pooled library members of each library in a weighted fashion and utilized sequential golden gate assembly with CcdB dropout selection to increase cloning efficiency (Methods, **Supplementary Figs. 2 and 3**).

To recruit these epigenetic effectors to an endogenous gene promoter, we used dCas9 targeting in combination with the MS2-MCP aptamer system derived from RNA bacteriophage MS2^32^, analogous to our prior designs that utilize sgRNAs with embedded MS2 stem-loops^10,33^. For temporal control over the editing complex, doxycycline-inducible promoters drove the expression of both dCas9 and the effector combination fused to monomeric MS2-coat protein (MCP) or synonymous mutated tandem MCP (stdMCP)^34^ (**Supplementary Fig. 4a,e**). To assess transcriptional outcomes on an endogenous target locus, we chose the gene encoding membrane protein CD81, whose expression can be measured using flow cytometry. We designed a three sgRNA expression cassette targeting upstream of the CD81 transcription start site (TSS) and confirmed the ability to transiently activate and repress CD81 expression with this experimental setup. Using well-characterized effectors including KRAB and the transcriptional activation domain of HSF1, CD81 expression levels showed a good dynamic range of both repression and activation during doxycycline treatment, and returned to baseline at 9 days after the removal of doxycycline (Methods, **Supplementary Fig. 4b**-d,**f-h**).

For our screen, we transduced the lentiviral libraries encoding Library 1 or Library 2 into K562 cells containing Tet-On dCas9 at a low multiplicity of infection (MOI<0.3) with a minimum effector coverage of 100X (Methods, **Fig. 1d** and **Supplementary Figs. 5a and 6a**). The combinatorial effector was recruited to CD81 loci for 5-6 days by adding doxycycline, followed by cell sorting based on CD81 expression (Day 0 timepoint to evaluate immediate effects on gene expression) (Methods, **Supplementary Figs. 5b**-d and 6b**-d**). This was followed by a 12-day effector washout period without doxycycline and a final round of cell sorting (Day 12 timepoint to evaluate durable effects on gene expression in the absence of effectors).

The size differences between Library 1 and Library 2 effector domain combinations (**Fig. 1e**) necessitated different NGS readout strategies. The majority (∼82%) of bivalent members in Library 1 fell below the 1500 nucleotide upper limit for short-read NGS platforms^35^, enabling us to directly sequence the N- and C-termini of effector pairs using short-read sequencing (**Supplementary Fig. 5a**). In contrast, Library 2 often exceeded this limit. To overcome this, we included a short 20N barcode in the cloning process, which was initially mapped to the effector pair by nanopore sequencing and read out by short-read sequencing at the end of the screen (Methods, **Supplementary Figs. 3g**-k and 6a,**e**). Targeted nanopore sequencing of integrated Library 2 members indicated less than 10% barcode swapping (**Supplementary Fig. 7**). Our barcoded approach with exponential effector pooling for Library 2 resulted in a reduction of the Day 0 dropout rate from 42.9% in Library 1 to 4.2% and successfully offset the exponential decay of effector coverage with increasing length that was observed for Library 1 and previous combinatorial screening approaches (**Supplementary Fig. 8a,d**)^29,36^. Effector dropout on Day 12 remained consistent at 42.1% for Library 1 and increased to 19.3% for Library 2 (**Supplementary Fig. 8b,c,e**).

The transcriptional effects of each domain combination were determined by measuring the relative abundance of each unique effector pair in the sorted cells from the high and low CD81 expression bins, calculated as log_2_ fold enrichment. Both libraries effectively captured the temporal dynamics of CD81 transcriptional perturbation induced by well-established control effectors. In Library 1, target gene repression by combinations containing the KRAB domain from ZNF10 was strong at Day 0 and disappeared by Day 12 (log_2_ fold score of -4.7 and -0.24, **Fig. 1f, g** and **Supplementary Table 3**). Similarly, repression by full-length RYBP, a known epigenetic silencer that recruits Polycomb repressive complex 1 (PRC1)^37^, and activation by the activation domain of HSF1 were captured at Day 0 and dissipated by Day 12 after dox removal for Library 2 (average log_2_ fold scores of -3.6 and 0.083 for RYBP and 1.9 and 0.047 for HSF1, **Fig. 1h, i** and **Supplementary Table 4**). Based on both CD81 expression levels measured by flow cytometry and HTS enrichment scores (**Supplementary Fig. 5c and Supplementary Tables 3** and **4**), the majority of transcriptional effects disappeared by 12 days after the removal of doxycycline (Day 12), highlighting the transient nature of perturbations introduced by the majority of epigenetic and transcriptional modifiers.

### COMBINE generates rich combinatorial landscapes of pairwise epigenetic interactions

To visualize transient perturbation outcomes across the large space of combinations tested, we generated enrichment score heatmaps (**Fig. 2a, b** and **Supplementary Fig. 9**). We observed clear diagonal symmetry between the X-Y and Y-X orientations of the same two effectors, with a high degree of correlation across pairs, indicating that most effector pairs behaved similarly regardless of orientation (Library: 1 r = 0.75; Library 2: r = 0.69, **Supplementary Fig. 10**).

**Figure 2.**
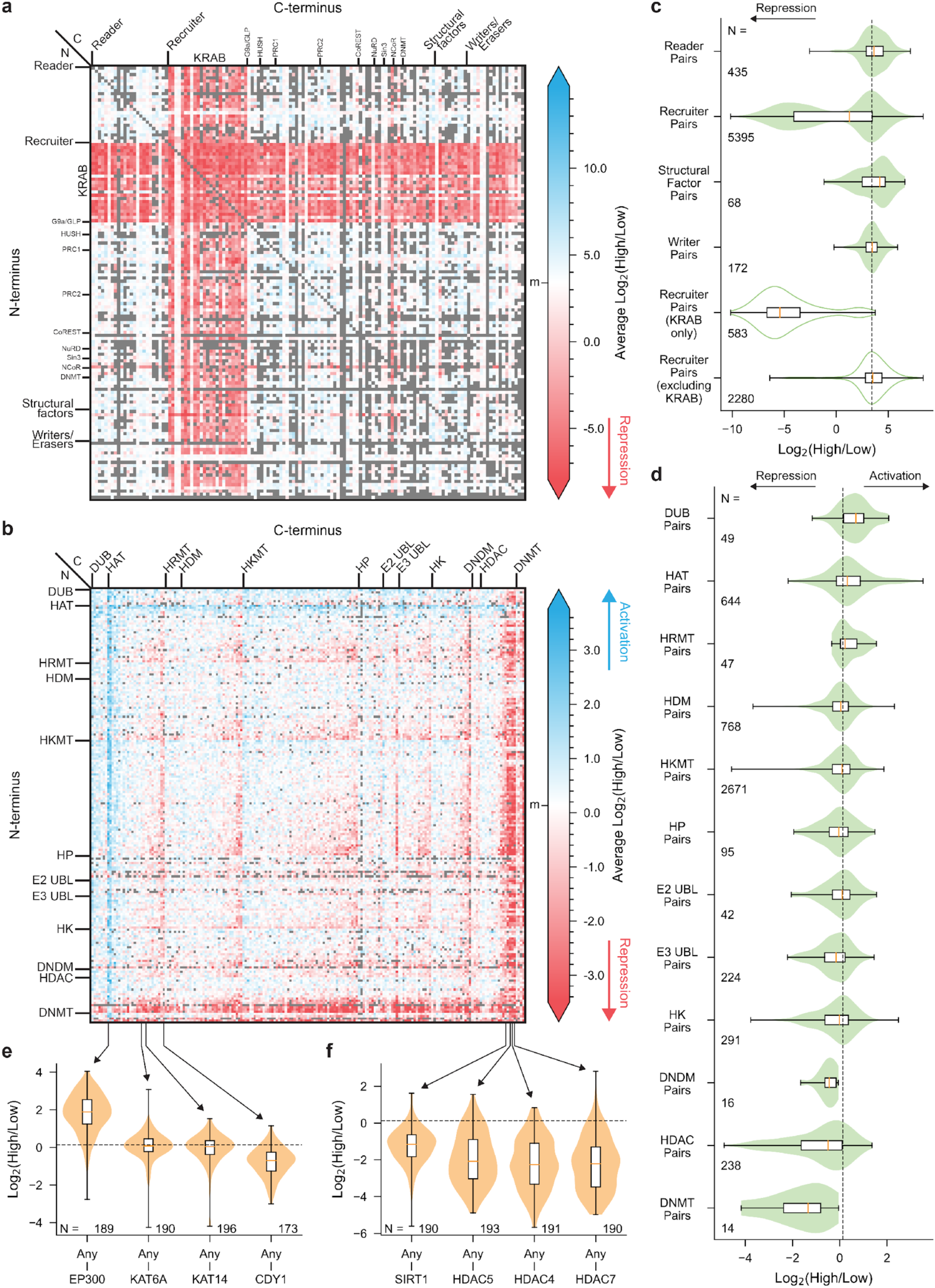
Epigenetic pairs generate a rich combinatorial landscape of transient gene expression outcomes. **(a)** Heatmap of enrichment scores generated from Library 1 at Day 0. Average Log_2_(High/Low) enrichment scores across both replicates are plotted. Color scale is centered at the mode of the enrichment score distribution, indicated by the “m” tick. For this repressor-focused library, lower log_2_ scores indicate stronger repression. Dropouts are indicated in gray. (**b**) Heatmap of enrichment scores generated from Library 2 at Day 0. Average Log_2_(High/Low) enrichment scores across both replicates are plotted. Color scale is centered at the mode of the enrichment score distribution, indicated by the “m” tick. For this library composed of putative repressors and activators, lower scores indicate more repression and higher scores indicate more activation. Histone deubiquitinase (DUB), histone acetyltransferase (HAT), histone arginine methyltransferase (HRMT), histone demethylase (HDM), histone lysine methyltransferase (HKMT), histone phosphatase (HP), E2 ubiquitin ligases (E2 UBL), E3 ubiquitin ligases (E3 UBL), histone kinase (HK), DNA demethylation machinery (DNDM), histone deacetylase (HDAC), and DNA methyltransferase (DNMT). (**c**) Violin plots of all possible bivalent combinations within the same classes from Library 1. A breakdown of the recruiter class into KRAB and non-KRAB pairs is also shown (not shaded). The orange bar represents the median. Box spans quartile 1 to quartile 3. Whiskers extend to minimum and maximum. Number of combinations per class is indicated. The dashed line indicates the distribution mode of all pairs. (**d**) Violin plots of all possible bivalent combinations within the same classes from Library 2. The orange bar represents the median. Box spans quartile 1 to quartile 3. Whiskers extend to minimum and maximum. Number of combinations per class is indicated. The dashed line indicates the distribution mode of all pairs. (**e**) Violin plots of all possible bivalent combinations with the specified HAT effectors on the C-terminus. The orange bar represents the median. Box spans quartile 1 to quartile 3. Whiskers extend to minimum and maximum. Number of combinations per effector is indicated. The dashed line indicates the distribution mode of all pairs. (**f**) Violin plots of all possible bivalent combinations with the specified HDAC effectors on the C-terminus. The orange bar represents the median. Box spans quartile 1 to quartile 3. Whiskers extend to minimum and maximum. Number of combinations per effector is indicated. The dashed line indicates the distribution mode of all pairs.

To identify general trends for the transcriptional impact of effector classes, we grouped pairs based on the families of individual effectors (**Fig. 2c,d**). While Library 1 revealed the strongest repressive phenotypes among KRAB family pairs, Library 2 revealed strong repression across pairs of histone deacetylases (HDACs) and DNA methyltransferases (DNMTs). We also observed a surprising trend toward mild target gene repression with pairs of DNA demethylation machinery (DNDM) domains. Pairs of histone acetyltransferases (HATs) tended to activate gene expression, as did histone deubiquitinases (DUBs) despite their ability to remove both activating and repressive histone ubiquitination marks. Finally, pairs of histone lysine methyltransferases (HKMTs) and histone kinases (HKs) ranged broadly across both activation and repression outcomes.

Next, we examined the directionality of individual effectors in the context of these class-level trends. In the HAT class, pairs containing C-terminal EP300 (a broad HAT capable of acetylating all four core histones^38^) tended to activate transcription, whereas combinations containing C-terminal CDY1 (which displays a strong acetylation preference for histone H4^39^) tended to repress transcription (**Fig. 2e**). In addition, some HAT effectors, such as KAT6A or KAT14 (a broad histone and weak H4 acetyltransferase, respectively^40,41^), exhibited more partner-dependent effects.

Compared to the HAT class, the HDAC domain class had more members that consistently repressed transcription irrespective of their N-terminal partners such as SIRT1 (a broad histone deacetylase and decrotonylase^42^) and class IIa HDACs (HDAC5, HDAC4, and HDAC7, **Fig. 2f**). However, the degree of repression ranged widely depending on the exact effector pairing. For example, the combination of EP300 and HDAC5 activated transcription with an average log_2_ fold enrichment of 1.1, while the pairing of UBE2E1 and HDAC5 repressed transcription with a score of -4.5. These global observations motivated us to quantitatively analyze the contribution of individual domain effects and domain-domain interactions to gene expression outcomes.

### Marginal effector analysis reveals how individual domains affect transcription irrespective of partners

To quantitatively assess how each of the individual effector domains in our two libraries contribute to epigenetic control of gene expression, we calculated a ‘marginal effector score’ for each domain (**Fig. 3a, b**). This score reflects the average enrichment value across all bivalent combinations containing the domain of interest relative to all combinations lacking it, yielding a measurement of each domain’s relative contribution to target gene expression (positive or negative) across pairings.

**Figure 3.**
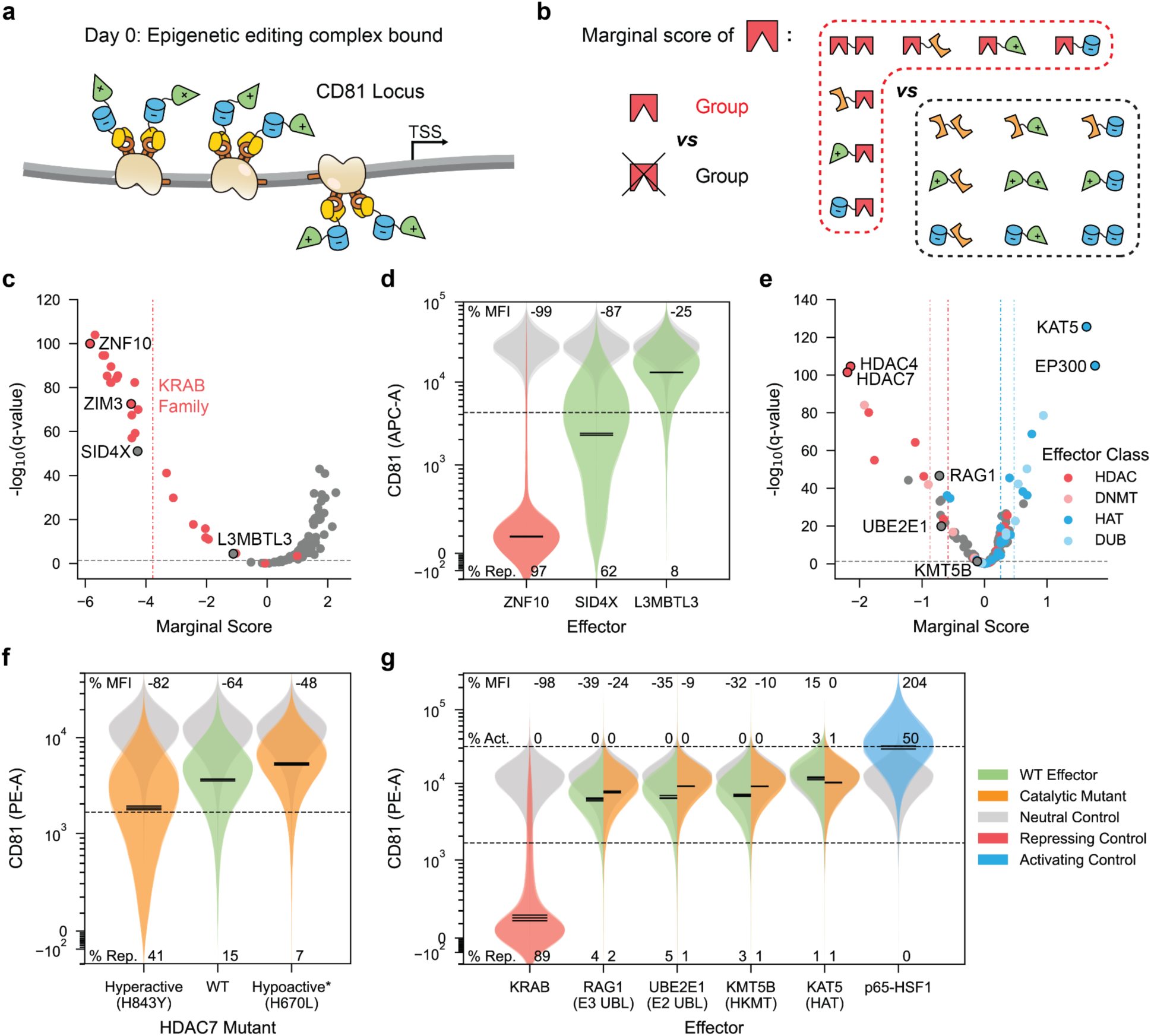
Marginal effector analysis reveals how individual domains perturb transcription across partners. (**a**) Schematic of the dCas9/MS2 epigenetic editing complex bound at CD81 locus. Complex expression is induced by the addition of doxycycline and is recruited to the CD81 loci for 5-6 days prior to screen readout. (**b**) Concept of marginal effector analysis. For a given effector, the marginal score is calculated as the difference in mean log_2_(High/Low) between all pairs containing the effector versus not containing the effector. Q-values were calculated as FDR-corrected p-values from Welsh’s t-test. (**c**) Volcano plot of marginal scores from Library 1 at Day 0. KRAB family members are highlighted in red. Vertical dashed line indicates the average marginal score across the KRAB family. Horizontal dashed line indicates q-value of 0.05. (**d**) Violin plots of CD81 expression after expression of individual effectors from Library 1 in individual validation (3 days post nucleofection of effector encoding plasmids). Mock transfection of pUC19 colored in gray. 2 independent replicates illustrated as translucent overlays. The geometric means of each replicate are shown as solid black lines. Dashed line indicates repression gate at the 1st percentile of the pUC19 condition. The average percentages of repressed cells in each condition are indicated. The average percent changes in CD81 mean fluorescence intensity (MFI) versus pUC19 for each condition are indicated. (**e**) Volcano plot of marginal scores from Library 2 at Day 0. Effectors from HDAC, DNMT, HAT, and DUB classes are highlighted in red, pink, blue, and sky blue respectively. Vertical dashed lines indicate average marginal score for each respective colored class. Horizontal dashed line indicates q-value of 0.05. (**f**) Violin plots of CD81 expression after expression of the HDAC7 deacetylase domain and its mutants in individual validation (5 days post nucleofection and doxycycline induction). WT HDAC7 is colored green. Mutants are colored orange. MCP-DMD neutral condition is colored in gray. 3 independent replicates are illustrated as translucent overlays. The geometric means of each replicate are shown as solid black lines. Dashed line indicates repression gate at the 1st percentile of the neutral condition. The average percentages of repressed cells in each condition are indicated. The average percent changes in CD81 MFI versus DMD for each condition are indicated. *proposed mutant inferred from HDAC4. (**g**) Violin plots of CD81 expression after expression of individual effectors from Library 2 in individual validation (5 days post nucleofection and doxycycline induction). WT effectors are colored green. Catalytic mutants are colored orange. Repressing and activating controls are colored red and blue respectively. MCP-DMD neutral condition is colored gray. 3 independent replicates are illustrated as translucent overlays. The geometric means of each replicate are shown as solid black lines. Dashed lines indicate repression and activation gate at the 1st and 99th percentile of neutral condition. The average percentages of repressed and activated cells in each condition are indicated. The average percent changes in CD81 MFI versus the neutral DMD domain for each condition are indicated.

Marginal effector analysis of Library 1 clearly reflected the well-known repressive effect of KRAB domains on gene expression^16,24^. The average marginal score across the KRAB domain class was -3.8, indicating an average 14-fold decrease in enrichment scores when KRAB is present in an effector pair (2^3.8^ ≈ 14, **Fig. 3c** and **Supplementary Table 3**). The ZNF10 (KOX1) KRAB domain – which is commonly used in CRISPR interference (CRISPRi) applications^43^ – exhibited the lowest marginal score at -5.8. The more recently identified ZIM3 KRAB domain also exhibited strong repression with a score of -4.5^20^. Notably, only two out of 25 tested KRAB domains did not consistently repress gene expression (ZNF791 and ZNF823; marginal scores of -0.1 and 1.0), suggesting that this family contains a wide spectrum of additional potential tool domains that have yet to be explored.

Outside the KRAB domain family, we identified two other individual domains with significantly repressive marginal scores in Library 1: SID4X (marginal score -4.3) and the sterile alpha motif (SAM) domain of L3MBTL3 (marginal score -1.1). SID4X is an engineered ternary version of the Sin3 interacting domain (SID) from the MAX dimerization protein 1 (MXD1) that efficiently recruits the Sin3-HDAC complex^44^, and the SAM domain of L3MBTL3 is known to multimerize and recruit lysine-specific histone demethylase 1A (LSD1/KDM1A)^45,46^, providing clear mechanistic hypotheses for the repressive effects of each of these domains.

To independently validate each domain’s individual contribution to transcriptional repression outside of the bivalent effector context, we fused ZNF10 KRAB, SID4X, or L3MBTL3 SAM to stdMCP and expressed them in the Tet-On dCas9 K562 cell line alongside a CD81-targeting guide array (**Fig. 3d**). Repression of CD81 target gene expression, as measured by the change in total mean fluorescence intensity (MFI), correlated with each domain’s relative marginal effect score: ZNF10 KRAB decreased CD81 MFI by 99%, SID4X by 90%, and L3MBTL3 by 46% relative to the negative control. Notably, unlike the all-or-nothing silencing pattern of KRAB^16^, neither SID4X nor L3MBTL3 fully silenced CD81, but rather resulted in more graded repression.

The marginal scores calculated from Library 2 were lower in magnitude, likely reflecting the more varied composition of this library in comparison to the repression-focused Library 1 (**Fig. 3e** and **Supplementary Table 4**). At the class level, the DNMT class exhibited the highest average repression, with a moderate effect size but high consistency across members (average marginal score -0.87). The DUB class resulted in the highest average gene activation with moderate but consistent effects (average marginal score 0.47). At the individual effector level, five of the seven top repressive domains in Library 2 belonged to the HDAC class (including HDAC7 and HDAC4 with scores of -2.2 and -2.1, respectively), while the HAT class contained the two strongest individual activators (EP300 and KAT5 with scores of 1.8 and 1.6, respectively).

Next, we confirmed the effects of 5 individual domains by fusing them to MCP and transiently expressing them in the same cell line used above. The strongest repressor, HDAC7 (marginal score -2.2), reduced CD81 MFI the most, by 64%, and was modulated by the introduction of point mutations (**Fig. 3f**). The hyperactive H843Y mutant^47^ increased repression to 82%, and the hypoactive H670L mutant^48^ diminished repression to 48% (Methods, **Supplementary Table 5**). Three additional repressors from diverse classes also repressed CD81 in concordance with their marginal scores. RAG1 (marginal score -0.72), an E3 ubiquitin ligase (E3 UBL) associated with histone H3 monoubiquitination during V(D)J recombination^49,50^, repressed CD81 by 39%; UBE2E1 (marginal score -0.69), an E2 ubiquitin ligase (E2 UBL) associated with PRC1^51^, repressed CD81 by 35%; and KMT5B (marginal score -0.12), a histone lysine methyltransferase (HKMT) that writes H3K20 methylation^52^ – a modification for which no targeted epigenetic editing tools currently exist, repressed CD81 by 32% (**Fig. 3g**). Catalytic mutations of these three enzymes^53–55^ reduced CD81 repression to 24%, 9%, and 10% respectively.

For activation, KAT5 (marginal score 1.6) was the second-ranking activator of gene expression in the screen, and validation experiments confirmed that it resulted in a 15% increase in CD81 MFI (**Fig. 3g**) which was completely abolished upon introduction of a catalytic point mutation. Altogether, our marginal effector analysis and validations of domains from Library 2 indicated a wide range of repressive and activating effects.

### COMBINE reveals synergy and antagonism in epigenetic domain interactions

Beyond the effects of individual domains on target gene expression, a key advantage of COMBINE is the ability to quantify synergistic or antagonistic interactions among domain pairs. To systematically assess domain interactions, we calculated a ‘synergy score’ for each effector pair that uses individual marginal scores to calculate the difference between expected and observed effector strength for each domain combination (Methods, **Fig. 4a**). Positive scores indicate synergy and negative scores indicate antagonism in the direction of the stronger marginal effector. Across both libraries, we identified many strong synergistic or antagonistic interactions between domains that provide a rich resource for further exploration (**Supplementary Figs. 11 and 12 and Supplementary Tables 6** and **7**). Compellingly, our analysis highlighted synergistic interactions between several PRC1 recruiters and a reader of H2AK119 monoubiquitination, recapitulating the known reader-writer synergy of variant PRC1 (**Supplementary Fig. 13 and Supplementary Note 1**)^37^.

**Figure 4.**
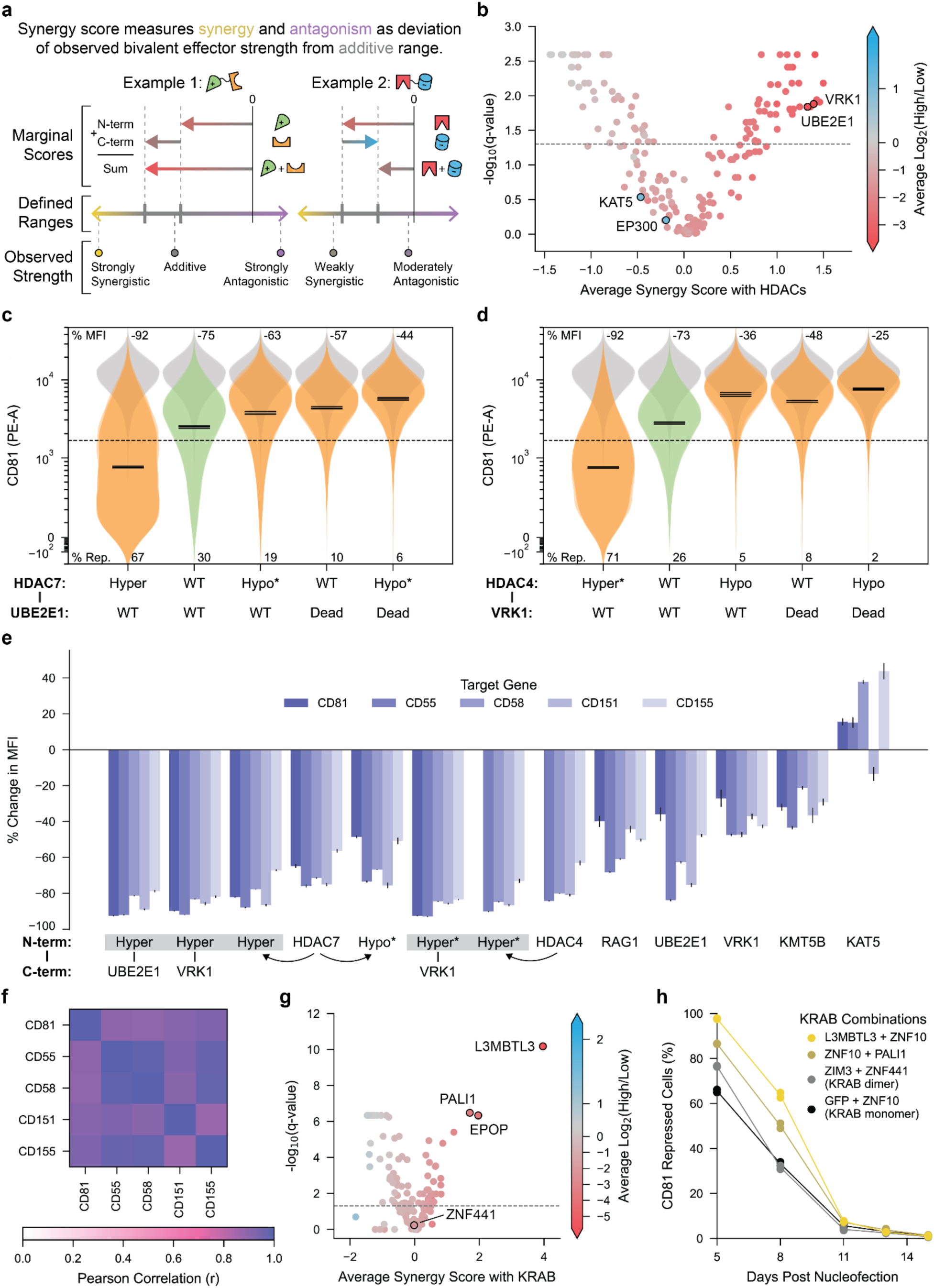
COMBINE uncovers diverse modes of epigenetic interactions. (**a**) Concept of synergy score calculation to quantify synergy and antagonism. Marginal scores of each individual effector are used to calculate an additive range for each bivalent combination. The lower bound of the additive range is defined as the minimum of the N-terminal effector’s marginal score, the C-terminal effector’s marginal score, and the sum of the marginal scores. The upper bound is defined as either the sum of the marginal scores or the minimum of the N-terminal and C-terminal marginal scores (whichever is higher). The magnitude of the synergy score represents how far outside the additive range the mean-centered observed bivalent enrichment score falls. Synergy is indicated by positive synergy scores, while antagonism is indicated by negative synergy scores. Two example cases illustrate the calculation of synergy scores. Library 1 synergy scores are all calculated in the repression direction. Library 2 synergy scores are calculated both in the activation and repression direction depending on whether the sum of marginal scores is positive or negative, respectively. (**b**) Volcano plot of average synergy scores with HDAC family from Library 2 on Day 0. Effectors are colored based on the average log_2_(High/Low) enrichment scores across all combinations of the given effector with significant HDAC repressors. Q-values were calculated as FDR-corrected p-values from a one-sample Wilcoxon test. Horizontal dashed line indicates q-value of 0.05. (**c**) Violin plots of CD81 expression at 5 days after nucleofection and induction of the HDAC7 + UBE2E1 combination and its mutants. WT domains are colored green. Mutant combinations are colored orange. MCP-DMD neutral condition is colored in gray. 3 independent replicates are illustrated as translucent overlays. The geometric means of each replicate are shown as solid black lines. Dashed line indicates repression gate at the 1st percentile of neutral condition. The average percentages of repressed cells in each condition are indicated. The average percent changes in CD81 MFI versus DMD for each condition are indicated. *proposed mutant inferred from homology to HDAC4. (**d**) Violin plots of CD81 expression at 5 days after nucleofection and induction of HDAC4 + VRK1 and enzymatic mutants. WT domains are colored in green. Mutant combinations are colored orange. MCP-DMD neutral condition colored gray. 3 independent replicates are illustrated as translucent overlays. The geometric means of each replicate are shown as solid black lines. Dashed line indicates repression gate at the 1st percentile of neutral condition. The average percentages of repressed cells in each condition are indicated. The average percent changes in CD81 MFI versus DMD for each condition are indicated. *proposed mutant inferred from HDAC7. (**e**) Percent change in MFI relative to the neutral DMD control following dual nucleofection of plasmids encoding key transient effectors from Library 2 and 3X guide arrays targeting CD55, CD58, CD151, and CD155. CD81 percent MFIs are from single plasmid nucleofection into Tet-On dCas9 cell line with preinstalled 3X CD81 guide array. The dCas9/MS2 complex was induced by doxycycline for 5 days. Error bars represent the standard deviation between 3 independent replicates. *, proposed mutant inferred from homology. Lack of C-terminal effector indicates that the effector was tested monovalently. Arrows point from WT effector to mutant form of the same effector. Gray boxes indicate the same effector. Additional detailed data are shown in **Supplementary Fig. 16**. (**f**) Heatmap of Pearson correlations of percent change in MFI versus the neutral DMD control between each gene tested. Datasets consist of the 13 effectors shown in (**e**). Additional detailed data are shown in **Supplementary Fig. 17**. (**g**) Volcano plot of average synergy scores with KRAB family from Library 1 on Day 6. Effectors are colored based on the average log_2_(High/Low) enrichment scores across all combinations of the given effector with significant KRAB repressors. Q-values were calculated as FDR-corrected p-values from a one-sample Wilcoxon test. Horizontal dashed line indicates q-value of 0.05. (**h**) Timecourse of CD81 repression following nucleofection of plasmids encoding KRAB combinations (n=2 independent replicates).

We were intrigued by the broad interaction patterns observed across the robust repressive domain classes, KRAB and HDAC. When analyzing the synergy scores for all KRAB and HDAC members with clear individual repressive effects (marginal scores below -0.5, **Supplementary Fig. 14**), we were able to classify additive, synergistic, and antagonistic effector pairings with other library members. For example, we observed that the strongly activating HAT members KAT5 and EP300 fully canceled out the repressive effect of HDACs in a way that was explained by a direct additive relationship (**Fig. 4b**). In contrast, 46 effectors resulted in synergistic enhancement of repression by HDAC domains (**Supplementary Table 7**). We selected two pairs with particularly high synergy scores for experimental validation: HDAC7 + UBE2E1 (synergy score 2.4) and HDAC4 + VRK1 (synergy score 2.0).

In validation experiments, we found that the HDAC7 + UBE2E1 and HDAC4 + VRK1 combinations reduced CD81 MFI by 75% and 73%, respectively, relative to the neutral control. This enhanced repressive effect was at least partially dependent on the catalytic activities of both effectors, as hypoactive mutations in the HDAC partners^48^ reduced each combination’s repression level to 63% and 36%, respectively, while loss-of-function mutations in the UBE2E1 and VRK1 partners^54,56^ attenuated repression to 57% and 48%, respectively (**Fig. 4c, d**).

Installing hyperactive mutations into the HDAC partner^47^ boosted CD81 repression to 92% for both combinations, further emphasizing the role of HDAC catalytic activity in this outcome.

As an E2 UBL associated with PRC1, UBE2E1 participates in the ubiquitination of H2AK119 to promote gene repression^51^. We hypothesize that gene repression by HDACs is enhanced by these epigenetic effectors since the removal of histone acetylation marks near the CD81 locus (**Supplementary Fig. 15a**)^57^ could increase the efficiency of writing repressive marks through established PRC crosstalk mechanisms^4^. While future mechanistic studies are needed, these results demonstrate the ability of our combinatorial screening platform to generate novel mechanistic hypotheses about fundamental biological processes.

To address the generalizability of our potent HDAC combinations as well as individual domains identified from Library 2, we tested 13 key effectors on a panel of 4 additional basally expressed genes including CD55, CD58, CD151, and CD155. Five days post-nucleofection of both effector and 3X guide array plasmids, we observed that the percent changes in MFI for each of the effectors were similar across all tested genes (**Fig. 4e** and **Supplementary Fig. 16**). The percent changes were highly correlated between genes, spanning a range of r = 0.87-0.98 and demonstrating that our findings are broadly applicable to other target genes (**Fig. 4f** and **Supplementary Fig. 17**).

Next, we examined synergy scores among the KRAB family of repressive domains to discern partner domains that could enhance or antagonize KRAB activity. Our synergy score analyses identified the SAM domain from L3MBTL3 and two PRC2 recruiters (PALI1 and EPOP) as the most potent synergistic partners of KRAB domains, all exhibiting both high average synergy scores and low enrichment scores on Day 6 (average synergy scores 4.0, 2.0, and 1.7 and log_2_ fold enrichments -5.7, -3.7, and -3.4, respectively) (**Fig. 4g**, **Supplementary Fig. 11b and Supplementary Table 6**). In individual validation of these domain combinations, all three KRAB combinations – L3MBTL3, PALI1, and EPOP – further enhanced repression of CD81 compared to the ZNF10 KRAB domain alone (L3MBTL3 + KRAB, KRAB + PALI1, GFP + KRAB were 97.7%, 86.6%, and 65.6% of CD81-repressed cells, **Fig. 4h** and **Supplementary Fig. 18a,b**).

PALI1 and EPOP are vertebrate-specific and mutually exclusive PRC2 subunits^58^ that can recruit the PRC2.1 complex, thereby promoting the deposition of H3K27 trimethylation^59^. In our screen, the PRC2 interacting domains (PID) of PALI1 [1058 - 1250 aa] and EPOP [300 - 379 aa] exhibited synergy with KRAB. KRAB + PALI1 PID and KRAB + EPOP PID combinations are expected to install both H3K9 methylation marks via KRAB and H3K27 trimethylation marks through recruitment of PRC2. Notably, in its endogenous context, the full-length PALI1 protein [1 – 1557 aa] is also capable of installing both H3K9 and H3K27 methylation marks, as it contains both the H3K9 methyltransferase G9A interacting domain [310 - 723 aa] and PRC2 interacting domain [1058 - 1250 aa]^60^. Considering that both G9A and KRAB zinc-finger proteins are vertebrate-specific proteins, synergistic interactions with vertebrate-specific components of PRC2 complexes raises interesting questions about the co-evolution of epigenetic components. Overall, we demonstrate the ability of COMBINE to highlight naturally occurring synergistic combinations of epigenetic domains.

### KRAB + L3MBTL3 combination enables potent transcriptional repression in challenging conditions

Individual KRAB domains, including those of ZNF10 and ZIM3, have been harnessed as epigenetic repressors in many contexts^20,43^ and yielded strong individual marginal scores in our screen (-5.8 and -4.5, respectively). However, in certain contexts – for example, under dose-limited conditions due to viral delivery, or when recruited to a locus in a more indirect manner – further enhancement of repressive activity would be beneficial. To provide new tool options for these challenging conditions, we examined our screen data for synergistic combinations that would enhance the potency of these KRAB domains.

In our screen data as well as in individual validation, the synergistic pairing of ZNF10 KRAB with the SAM domain from L3MBTL3 was the most potent combination identified. Notably, the SAM domain is known for its ability to multimerize^45^, potentially recruiting multiple KRAB domains to the target gene (**Fig. 5a**). We tested this combination under dose-limiting conditions using low MOI lentiviral transduction of K562 cells with either the stdMCP-KRAB or stdMCP-KRAB-L3MBTL3 effectors combined with a single sgRNA – instead of three – targeting CD81 (**Supplementary Table 8**). We observed significantly enhanced repression of the combined KRAB + L3MBTL3 effector under these challenging conditions, particularly at low levels of doxycycline induction where we achieved up to ∼34-fold increased repression (KRAB: 1.4% vs KRAB + L3MBTL3: 48.1% at 5 ng/μL dox) (**Fig. 5b**). Even at standard 1000 ng/μL doxycycline conditions, the KRAB + L3MBTL3 combination exhibited ∼6-fold increased repression compared to KRAB alone.

**Figure 5.**
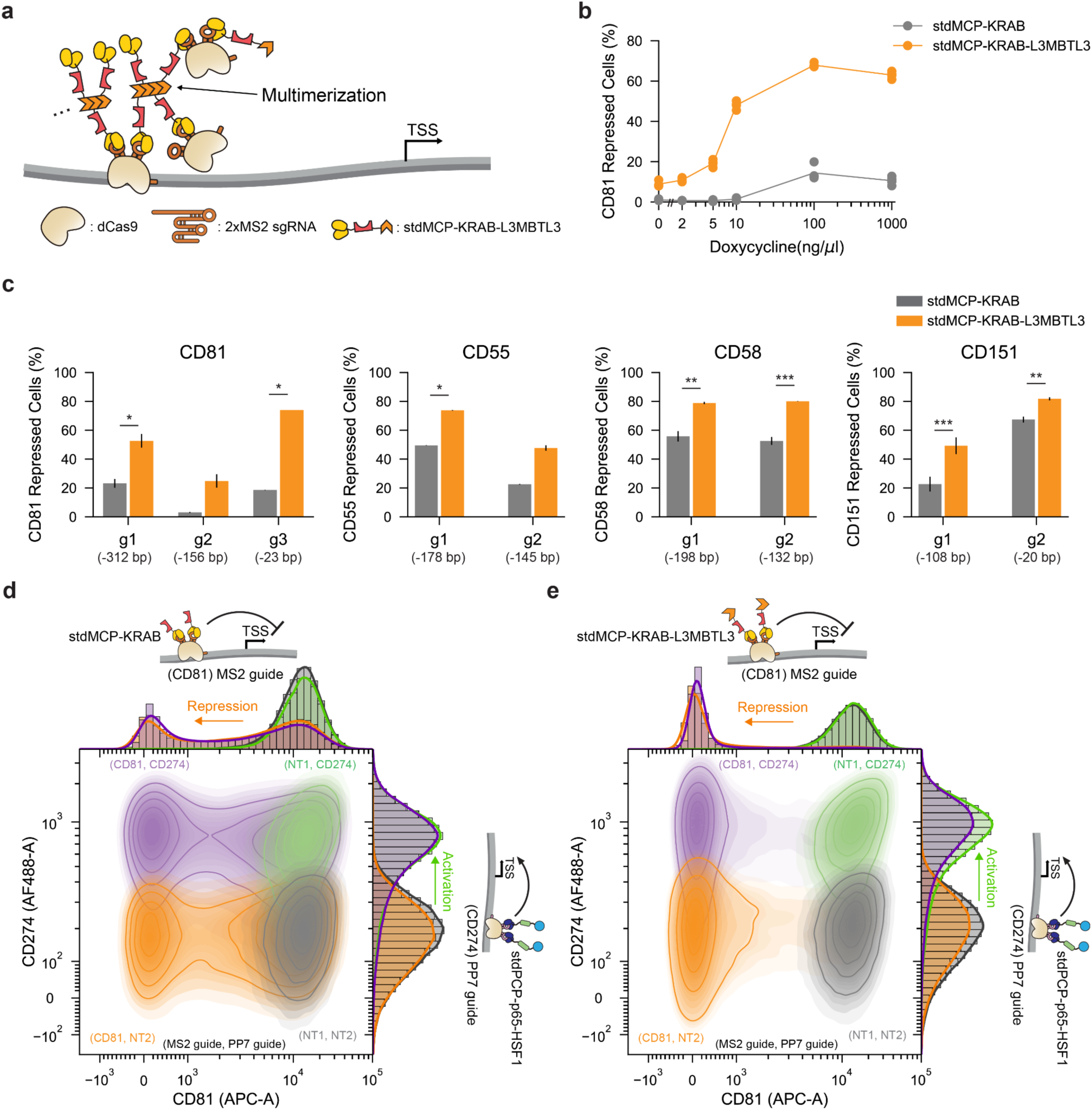
A combination of KRAB and L3MBTL3 enables robust and bidirectional transcriptional perturbation. (**a**) Illustration of hypothetical clustering of the dCas9/MS2 complex due to the multimerization property of the SAM domain of L3MBTL3. (**b**) Doxycycline response curve of CD81 repression by stdMCP-KRAB and stdMCP-KRAB-L3MBTL3 4 days after doxycycline induction (n = 4 biological replicates). KRAB domain is from ZNF10. (**c**) Target gene repression 5 days following lentiviral transduction with sgRNA and doxycycline induction of the effector components (n ≥ 2 biological replicates). Distance between the sgRNA target site and the TSS is indicated in parentheses. Error bars indicate the standard deviation of 4 biological replicates. * p-value < 0.05, ** p-value < 0.01, *** p-value < 0.001; two-tailed t-test. (**d**) 2D-contour plot and kernel density estimate (KDE) plots of populations transduced with dual MS2/PP7 sgRNAs. stdMCP-KRAB was recruited for CD81 gene repression and stdPCP-p65-HSF1 was recruited for the activation of CD274 gene expression. CD81/CD274 expression profiles were measured 5 days following lentiviral transduction with sgRNAs and doxycycline induction of the effector components. Genes within parentheses indicate the target genes for MS2 and PP7 sgRNAs, respectively. For example, (CD81, NT1) denotes that the CD81 gene is targeted by the MS2 sgRNA, while NT1 (non-target 1) is a control sgRNA used with PP7. (**e**) 2D-contour plot and kernel density estimate (KDE) plots of populations infected with dual MS2/PP7 sgRNAs. stdMCP-KRAB-L3MBTL3 was recruited for CD81 gene repression and stdPCP-p65-HSF1 was recruited for the activation of CD274 gene expression. CD81/CD274 expression profiles were measured 5 days following lentiviral transduction with sgRNAs and doxycycline induction of the effector components. Genes within parentheses indicate the target genes for MS2 and PP7 sgRNAs, respectively. For example, (CD81, NT1) denotes that the CD81 gene is targeted by the MS2 sgRNA, while NT1 (non-target 1) is a control sgRNA used with PP7.

To test the generalizability of this effect, we selected three additional gene targets encoding cell surface proteins with basal expression in K562 cells (CD55, CD58, and CD151). The KRAB + L3MBTL3 effector consistently demonstrated superior repression across three additional target genes compared to KRAB alone (**Fig. 5c**), ranging from 1.2-fold improvement with the highly effective CD151 targeting sgRNA to 8.0-fold for a weaker CD81 targeting sgRNA. Together, these results support the ability of this domain combination to overcome the limitations of weaker sgRNAs or lower effector expression to achieve robust repression across diverse target genes.

Beyond the general enhancement of CRISPRi activity in challenging settings, we reasoned that the increased efficacy afforded by the KRAB + L3MBTL3 combination could be particularly enabling for bidirectional CRISPR perturbations, in which two orthogonal epigenetic effectors can be targeted to two distinct loci to achieve upregulation of one gene and simultaneous repression of another^10,61,62^. To assess this possibility, we used the stdMCP fusion strategy to recruit KRAB alone or KRAB + L3MBTL3 to an MS2-containing CD81 sgRNA or non-targeting control guide, while using an analogous setup to recruit the p65-HSF1 CRISPRa effector^63^ using the orthogonal stdPCP-PP7 system^34,61^. We engineered cell lines to express dCas9, stdMCP-KRAB or stdMCP-KRAB-L3MBTL3, and stdPCP-p65-HSF1 in a doxycycline-inducible manner and introduced four dual MS2/PP7 sgRNA combinations – each containing either a CD81-targeting or a non-targeting MS2 sgRNA paired with a CD274-targeting or a non-targeting PP7 sgRNA – using lentivirus.

Importantly, we found that the KRAB + L3MBTL3 combination fully silenced CD81 expression to produce four distinct populations, unlike the KRAB-only condition (**Fig. 5d, e**). This improved separation resulted from augmented repression by the KRAB + L3MBTL3 combination (KRAB: 41.5% vs KRAB + L3MBTL3: 89.4% CD81 repressed cells) without interfering with CD274 activation by the PP7 system (KRAB: 66.5% vs KRAB + L3MBTL3: 74.5% CD274 activated cells) (**Supplementary Fig. 19**). The KRAB + L3MBTL3 combination therefore enabled us to overcome limitations in the efficacy of other MS2-based repressors and develop a robust MS2/PP7-based bidirectional perturbation system, which may be applied to future bidirectional CRISPR screening^64,65^ or therapeutic applications.

### Long-term epigenetic memory requires modifiers of DNA methylation

Thus far, we have demonstrated the power of COMBINE to identify new individual effectors and combinations of effectors that transiently perturb gene expression, accurately measuring effector strength and uncovering unexpected synergistic and antagonistic interactions. Next, we investigated durable effects on target gene expression produced by transient expression of the epigenetic editing complex (**Fig. 6a**). Following the initial 5-6 day effector recruitment phase, doxycycline was removed to halt any further production of dCas9 and the (std)MCP-fused effectors. Using our pilot studies as a reference, in which short-term repression by KRAB dissipated after twelve days (**Supplementary Fig. 4b**-d), we cultured cells for twelve days following doxycycline removal to our final screening timepoint. By Day 12, the majority of transcriptional effects had dissipated (**Fig. 1g, i** and **Supplementary Tables 3** and **4**).

**Figure 6.**
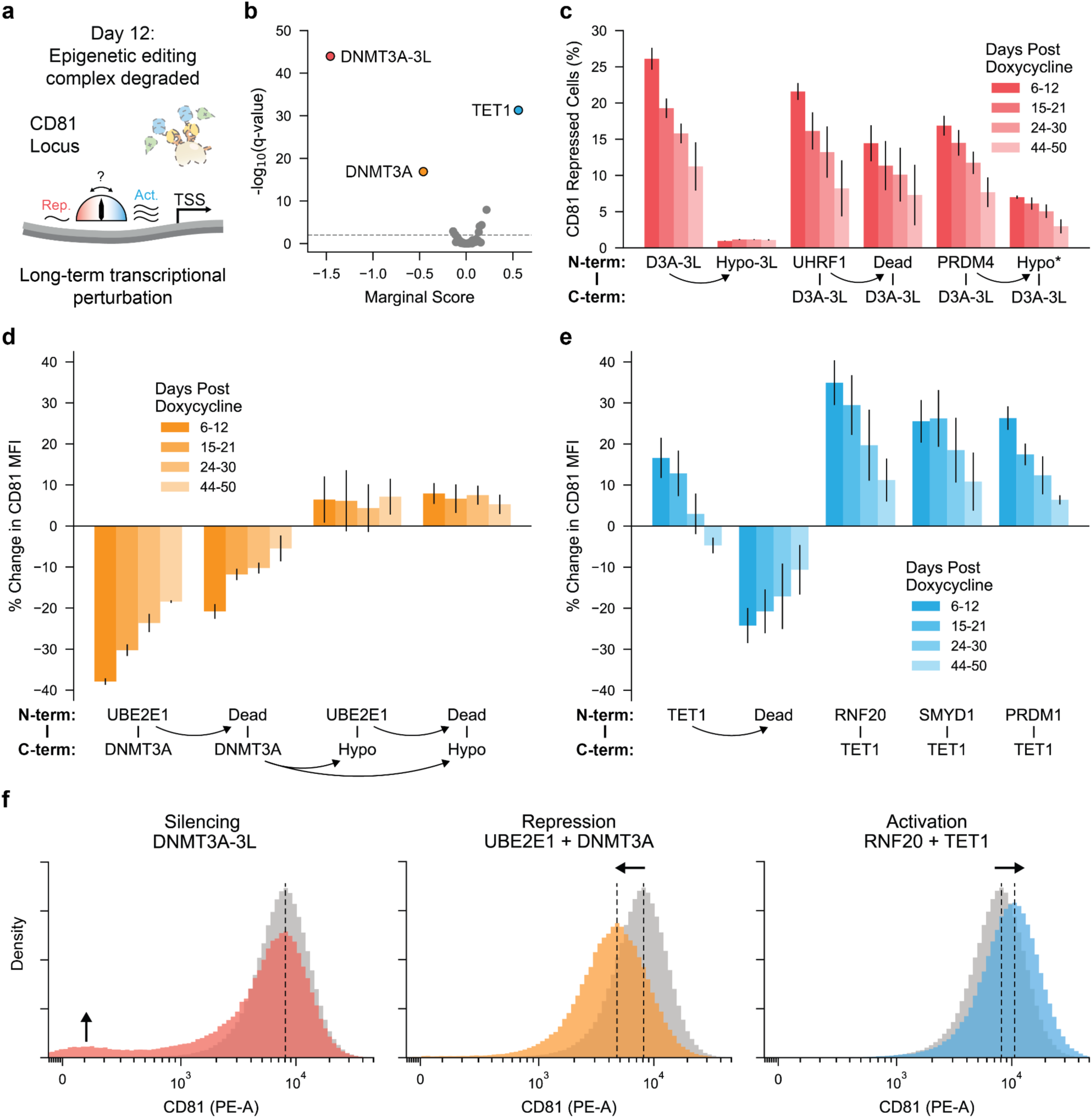
DNA methylation writers and erasers are essential for the memory of silencing, repression, and activation of gene expression. (**a**) Schematic showing loss of the dCas9/MS2 epigenetic editing complex 12 days post-withdrawal of doxycycline. CD81 expression perturbations rely on sustained epigenetic modifications. (**b**) Volcano plot of marginal scores from Library 2 on Day 12. Horizontal dashed line indicates q-value of 0.05. (**c**) Timecourse of CD81 repression following nucleofection of plasmids encoding DNMT3A-3L combinations and induction of the dCas9/MS2 complex by doxycycline treatment for 5 days. Day 0 indicates the day of doxycycline washout. Each bar represents 3 timepoints collected within the specified days. The bar heights represent the average perturbation across the timepoints, and the error bars represent the standard deviation between 3 independent replicates. *proposed mutant inferred from PRDM7. Lack of C-terminal effector indicates that the effector was tested monovalently. Arrows point from WT effector to mutant form of the same effector. Additional detailed data are shown in **Supplementary Fig. 21**. (**d**) Timecourse of CD81 MFI relative to the neutral DMD control following nucleofection of plasmids encoding UBE2E1 + DNMT3A combinations and induction of the dCas9/MS2 complex by doxycycline for 5 days. Day 0 indicates the day of doxycycline washout. Each bar represents 3 timepoints collected within the specified days. The bar heights represent the average perturbation across the timepoints, and the error bars represent the standard deviation between 3 independent replicates. Arrows point from WT effector to mutant form of the same effector. Additional detailed data are shown in **Supplementary Fig. 22**. (**e**) Timecourse of CD81 MFI relative to the neutral DMD control following nucleofection of plasmids encoding TET1 combinations and induction of the dCas9/MS2 complex by doxycycline for 5 days. Day 0 indicates the day of doxycycline washout. Each bar represents 3 timepoints collected within the specified days. The bar heights represent the average perturbation across the timepoints, and the error bars represent the standard deviation between 3 independent replicates. Lack of C-terminal effector indicates that the effector was tested monovalently. Arrows point from WT effector to mutant form of the same effector. Additional detailed data are shown in **Supplementary Figs. 23 and 24**. (**f**) Representative CD81 expression profiles illustrating three types of gene expression memory. CD81 expression levels were measured 12 days after doxycycline washout following 5 days of doxycycline induction.

Implementing the marginal score metric that we developed earlier allowed us to quantitatively determine the magnitude of long-term changes to target gene expression induced by each individual effector across different combinations. Across both libraries, only three individual effectors induced significant long-term transcriptional changes with absolute marginal scores greater than 0.5, and all of these effectors participate in DNA methylation (**Fig. 6b** and **Supplementary Fig. 20a**). Two of the identified effectors, DNMT3A-3L^17^ and DNMT3A, stably repressed target gene expression, while one effector, TET1, stably activated target gene expression. In contrast to DNMT3A-3L and DNMT3A, which are capable of *de novo* methylation of CpG sites^66–68^, TET1 oxidizes methylated cytosines to promote DNA demethylation^68,69^.

Notably, both DNMTs have been shown to induce long-term silencing^12,17,19^, while TET1 has been used to reactivate disease-relevant or synthetically silenced genes in long-term^12,70^ and upregulate target gene expression in a transient manner^68,71,72^. Implementing our synergy score metric on Day 12 screen data recovered the established synergistic interaction between the KRAB domain and DNMT3A – the basis for CRISPRoff technology^11,12^ – as the strongest hit in Library 1 (**Supplementary Fig. 20b,c**).

We followed up on each of these three domains - including different pairings suggested by our screen data - with individual validation experiments. DNMT3A-3L exhibited moderate but stable long-term repression both individually as well as in combinations identified to induce repressive memory in the screens (DNMT3A-3L, UHRF1 + DNMT3A-3L, PRDM4 + DNMT3A-3L, 23.4%, 19.7%, and 18.5% of cells repressed 12 days after doxycycline removal, respectively, **Fig. 6c** and **Supplementary Fig. 21a**-d). This repression was sustained for up to 50 days. We further tested a hypoactive mutant of UHRF1 and an inferred hypoactive mutant of PRDM4, mutating a conserved tyrosine that is catalytically active in closely related PRDM9^73,74^. Incorporating these mutations led to a 1.4-fold and 2.4-fold reduction in repression on average over the 50-day timecourse (**Supplementary Fig. 21e**). The RING finger domain of UHRF1 is an E3 UBL that catalyzes ubiquitination of H3K18 and H3K23, which in turn recruits and stimulates DNMT1 to faithfully maintain DNA methylation^75–78^. It follows that DNMT1 may aid DNMT3A-3L in establishing methylation patterns that are heritable, for example by converting *de novo* hemimethylated CpGs into fully methylated CpGs.

For DMNT3A, we selected the combination with UBE2E1 for individual validation. Interestingly, we observed that this combination led to minimal complete silencing (2.1% cells fully silenced at 12 days), but did result in a 40.5% reduction in mean CD81 expression across all cells (measured as MFI) at 12 days, which lasted up to 50 days (**Fig. 6d** and **Supplementary Fig. 22a**-c). Notably, this effect was reduced by 2.3-fold when UBE2E1 was mutated at its catalytic cysteine and completely abolished when DNMT3A was catalytically inactivated (**Supplementary Fig. 22d**)^54,79^. These results indicate that DNMT3A is the primary mediator of this repressive epigenetic memory, but that UBE2E1 augments DNMT3A’s activity, at least partially through its catalytic function as an E2 UBL. The ability to stably reduce gene expression without silencing will enable future functional genomics studies interrogating gene dosage as well as offer therapeutic downregulation of genes in situations where maintaining low levels of target gene expression is required.

In addition to long-term repression, we observed long-term epigenetic activation catalyzed by TET1 combinations. While TET1 has been previously used to reactivate the expression of target genes silenced by DNA methylation in synthetic or disease contexts^12,70^, our marginal effector score analysis suggests that TET1 can also increase the expression of a basally expressed target gene not only transiently^68,71,72^ but also in a long-term manner. For individual validation, we selected the three TET1 partners with the highest activation from our screen - RNF20’s RING finger domain, SMYD1’s SET domain, and PRDM1’s PR/SET domain. All three TET1 combinations led to an increase in CD81 expression that lasted up to 50 days and reached 40.3%, 28.7%, and 35.6% respectively at 6 days post-doxycycline washout (**Fig. 6e** and **Supplementary Fig. 23**). Activation by TET1 alone, on the other hand, only lasted up to 21 days and reached 19.5% at day 6. Mutation of the known catalytic residues of TET1 not only abrogated the activation phenotype^69^ but surprisingly led to stable gene repression, while catalytic mutations in the TET1 partners had a neutral or positive effect on long-term activation compared to their unmutated counterparts (**Supplementary Fig. 24**). These results indicate that other as-yet unknown factors, such as for example the recruitment of endogenous machinery, can augment long-term activation induced by TET1.

In summary, we delineated three distinct modes of epigenetic memory mediated by modulators of DNA methylation: silencing, reduction in gene expression, and activation (**Fig. 6f**). We further demonstrate the ability to tune the degree of long-term repression on endogenous genes and establish long-term activation of a basally expressed gene on a single cell level. Future work will delineate the mechanistic differences between the precise interactions of effector pairs and their resulting modes of epigenetic memory.

## Discussion

In this work, we devise a new platform technology called COMBINE for systematic, high-throughput functional screening of domain pairs. Importantly, COMBINE overcomes the length limitations and length-based biases of previous screening methodologies^28,29,36^ to enable the characterization of large protein domains and beyond, which we demonstrate by testing combinatorial domain candidates up to 2,094 amino acids in length.

Applying COMBINE to epigenetic regulation of gene expression, we test over 50,000 pairs of epigenetic reader, recruiter, writer, and eraser domains across two libraries to understand emergent properties of combinatorial domain interactions that illuminate new biology and provide a rich ground for future epigenome editing tool development. In particular, our systematic analysis allowed us to capture broad, class-level trends of epigenetic effector activity, discover new and unexpected synergistic or antagonistic domain interactions, and quantitatively understand the contribution of individual domains and domain pairs to the modulation of target gene expression. Importantly, we establish our screen results are quantitative and generalizable across 5 endogenous genes through extensive secondary validation. We highlight new domain combinations that may prove useful for next-generation epigenome editing tools, including the pairing of the ZNF10 KRAB domain with the SAM domain from L3MBTL3 which improved target gene silencing by up to 34-fold in challenging, dose-limiting conditions. This robust activity enabled the substantial improvement of an aptamer-based bidirectional CRISPR perturbation setup^61,62^, in which one target gene is silenced while another is activated using an orthogonal guide RNA-mediated effector recruitment strategy.

The inducible nature of the COMBINE platform allowed us to interrogate the durable effects of domain pairs on target gene transcription following inducer washout. From this long-term analysis, DNA methylation was nominated as the key modification driving heritable epigenetic regulation. Expanding beyond previously observed epigenetic silencing memory^11,12^, we also observed domain combinations that elicited long-term graded target gene repression of approximately 40%, or long-term gene activation of 30-40% – both of which were sustained for at least 50 days. Importantly, the latter example establishes the ability of TET1 pairs to initiate heritable gene activation from baseline expression levels and emphasizes the utility of our combinatorial screening approach to identify unexpected pairings that greatly enhance the potency and durability of this effect. We anticipate that these new epigenetic memory effectors will expand the toolbox of long-term expression modulation technologies, complementary to existing approaches such as CRISPRoff^12^.

Our rich dataset quantifying the transcriptional output induced by over 50,000 epigenetic effector pairs provides a broad resource for the epigenetics community to explore new mechanistic hypotheses and pursue new biotechnological tools. Further, the COMBINE platform is generalizable to many cell types due to its lentiviral delivery and is applicable beyond epigenetics to study many outstanding questions in combinatorial biology. The ability to systematically evaluate the functional interplay between entities–whether that includes DNA elements, RNA species, protein domains, or entire proteins–will unlock a sophisticated understanding of interactions in biology, where nothing acts in isolation.

## Methods

### Cell lines and cell culture

All experiments in this study were carried out in K562 cells (ATCC, CCL-243, female). Cells were cultured in a humidified incubator at 37°C and 5% CO_2_, in RPMI 1640 (Gibco, 61870036) media supplemented with 10% fetal bovine serum (FBS) (Gibco, 26140095) and 1% penicillin-streptomycin (Gibco, 15140122). For Library 1 lentivirus generation, 293FT (Thermo Fisher Scientific) or Lenti-X 293T (Takara) were cultured in a humidified incubator at 37°C and 5% CO_2_, in DMEM (Gibco, 10566016) media supplemented with 10% fetal bovine serum (FBS) (Gibco, 26140095) and 1% penicillin-streptomycin (Gibco, 15140122). For Library 2 lentivirus generation, viral production cells (Gibco, A35347) were cultured according to the manufacturer’s instructions.

### Guide design and cloning

CRISPRa guides were generated using CHOPCHOP for the CD81, CD55, CD58, CD151, and CD155 genes, and three were selected to tile each promoter region from -500-0 to the TSS (**Supplementary Table 8**, Guides)^80^. A tiered golden gate approach was developed to assemble a 3X guide array and to easily insert this array into a landing pad PiggyBac vector, which was cloned in part using the EMMA toolkit^81^, between divergent expression cassettes pTRE3G-dCas9-tagBFP and pEF1a-BlastR-P2A-rTA3 (**Supplementary Table 8**, Plasmids, pMH224). Briefly, the three guides were cloned separately via BsmBI golden gate into three distinct guide expression plasmids containing different golden gate overhangs around the guide cassette (**Supplementary Table 8**, Plasmids, pMH213-pMH215). The guide scaffold contained MS2 aptamers in stemloop two and the tetraloop and was sourced from recent optimization^33^. These individual guide plasmids were then assembled via BsaI golden gate into a RFP dropout landing pad containing pSV40-mCherry for mammalian expression (**Supplementary Table 8**, Plasmids, pMH222). Guides were arranged in the 3X array such that the closest guide to the TSS was placed first, followed by the middle guide and then the furthest guide. The CD81 3X guide array plasmid was used directly for guide nucleofection in screening of Library 1 (**Supplementary Table 8**, Plasmids, pMH228). This 3X array was further assembled into pMH224 via a final BsmBI golden gate for PiggyBac insertion and constitutive guide expression, which was used for screening Library 2 (**Supplementary Table 8**, Plasmids, pMH260). Additional sgRNAs targeting before the TSS of CD55, CD58, CD151, and CD274 were generated using CHOPCHOP (**Supplementary Table 8**, Guides)^80^. CD55, CD58, and CD151 guides were inserted into a lentiviral vector containing the recently optimized MS2 guide scaffold^33^ driven by human U6 promoter and SV40 promoter driven-mCherry marker. For bidirectional perturbation experiments, the CD274 guide (**Supplementary Table 8**, Guides) was cloned into the human U6-driven PP7 guide scaffold within a lentiviral vector, where both MS2 sgRNA and PP7 sgRNA are expressed from distinct mouse and human U6 promoters, respectively.

### PB Tet-On dCas9 cell line generation

To generate Tet-On dCas9 K562 cell lines expressing pTRE3G-dCas9-tagBFP, pEF1a-BlastR-P2A-rTA3 with or without the 3X CD81 guide array, K562 cells were nucleofected with 600 ng of pMH224 or pMH260 and 200 ng of a separate PiggyBac transposase plasmid using SF Cell Line 96-well Nucleofector kit and protocol (Lonza, V4SC-2096) (**Supplementary Table 8**, Plasmids). Two days following nucleofection, a 10-day selection began with 10 μg/mL blasticidin (Gibco, A1113903). After selection, cells were expanded and assessed for BFP induction in response to 1 μg/mL doxycycline. This Tet-On dCas9 K562 (with CD81 guide array) cell line was used for Library 2 HTS and Library 1 validation experiments, while this Tet-On dCas9 K562 (without CD81 guide array) was used for validation of Library 2 effectors on additional genes.

An additional Tet-On dCas9 K562 (without CD81 guide array) cell line was generated that expresses pTREtight-dCas9 and pEF1a-BlastR-P2A-rTA3. K562 cells were co-transfected with Piggybac plasmid encoding both pTREtight-dCas9 and pEF1a-BlastR-P2A-rTA3 (without transposases) and PiggyBac transposase plasmid using Lipofectamine 2000 (Invitrogen). Two days following transfection, 20 μg/mL blasticidin (Gibco, A1113903) selection was performed for at least 10 days. This Tet-On dCas9 K562 (without CD81 guide array) cell line was used for Library 1 HTS.

### Library 1 design and cloning

A literature review was conducted to cover critical components of the key repressive epigenetic multiprotein complexes and structural factors for chromatin organization. The majority of library members were selected based on previous arrayed studies employing truncation-based co-immunoprecipitation experiments or luciferase assays, except for the few members from high-throughput studies^24^. Peptides shorter than 80 amino acids were expanded equally on either side to reach 80 amino acids in total. Start codons were removed as effectors were to be fused to the C-terminus of aptamer binding protein. Amino acid sequences were back-translated with mammalian species codon optimization and forbidding 3′ end 6-mer of both amplification primers motifs (GGTGTG and ATGGCC) by Geneious Prime. Second codon optimization was performed for human codon usage, removing several type IIS restriction enzyme sites (BbsI, BsaI, BsmBI, BspQI, BtgZI, and SapI) and constraining GC content to between 35% and 65% in every 50 nucleotides window by DNA chisel^82^. Forward amplification primer binding handle (TCCaGACCGTTCaGGTGTG adapted from previous study^24^), BsaI binding site (GGTCTCT), and overhang (GTCA) facilitating ‘Glutamic-acid (E) – Serine (S)’ peptide overhang for C-terminus of XTEN16 linker were appended to 5′ of the gene fragment (**Supplementary Fig. 2a**). Overhang (AGTG) for ‘Serine (S) – Glycine (G)’ peptide overhang at N-terminus of XTEN16 linker, BsaI binding site (CGAGACC), and reverse amplification primer binding handle (gGCCaTGCGGaATGGGTTA) were appended to 3′ of the gene fragment. Optimized golden gate assembly overhangs were selected based on a previous study^83^. A few BsmBI sites, BsaI sites, and 3′ end 6-mer of both amplification primers motifs (GGTGTG and ATGGCC) that were unintentionally generated by former procedures were manually modified by silent mutations. Finally, library members that did not pass the Gblock or Eblock complexity test were human codon-optimized by IDT webtool, and again BsmBI, BsaI sites, and 3′ end 6-mer of both amplification primers motifs were manually modified by silent mutations.

Double-stranded gene fragments were synthesized (IDT and Twist Biosciences) and then pooled manually without PCR amplifications to make a 20 nM solution with the molarity of each fragment increasing proportional to its length with a slope of 1. For example, we added 1.5-fold of the 450 bp gene fragment compared to the 300 bp gene fragment. The pooled library was subcloned into KanR1_receiver (pCM001) and KanR2_reveiver (pCM002) plasmids by two separate golden gate reactions. For a total volume of 40 μL golden gate reaction, we used 185 ng of KanR1 (pCM001) or KanR2 (pCM002) receiver plasmid, 10 μL of 20 nM pooled library for ∼2:1 molar ratio between backbone:insert, 2 μL of BsaI-HFv2 (NEB, 20000 U/mL), 2 μL of T4 DNA ligase (NEB, 400000 U/mL), and 4 μL of 10X T4 DNA ligase buffer (NEB). The thermocycling protocol was 30 cycles of 5 min digestion at 37°C and 5 min ligation at 16°C, followed by a final 5 min digestion at 37°C, 5 min heat inactivation at 60°C and an extra 20 min heat inactivation at 70°C. The reactions were then purified with DNA Clean & Concentrator-5 (Zymo) and were eluted by 8 μL of nuclease-free water (Ambion). 1 μL of purified DNA was transformed into 25 μL of Endura electrocompetent cells (Lucigen) following the manufacturer’s instructions with 2 mL of prewarmed recovery media. 2 mL of cells were plated onto a 245 mm x 245 mm plate, and serially diluted cells were plated onto 100 mm diameter plates for the estimation of colony numbers. After overnight incubation at 30°C, colonies were collected, and plasmid libraries were extracted with Maxiprep (Machery-Nagel). Colony coverage was at least 34000X for 155 library members.

For the final bivalent library construction, the LV_AmpR_backbone (pCM003) was predigested with FastDigest Esp3I (Thermo Fisher Scientific) and gel extracted. For a total volume of 100 μL golden gate reaction, we used 800 ng of predigested and gel-extracted LV_AmpR_backbone (pCM003), 1000 ng of N-terminus effector (KanR1_recevier) library, 1000 ng of C-terminus effector (KanR2_receiver) library, 10 μL of FastDigest Esp3I (Thermo Fisher Scientific), 5 μL of T4 DNA ligase (NEB, 400000 U/mL), and 10 μL of 10X T4 DNA ligase buffer (NEB). The thermocycling protocol was initial 5 min digestion at 37°C, 30 cycles of 5 min digestion at 37°C and 10 min ligation at 16°C, final 20 min ligation at 16°C, final 5 min digestion at 37°C, and 20 min heat-inactivation at 75°C. The golden gate reaction was then isopropanol precipitated into 10 μL of nuclease-free water (Ambion)^84^. Again, 1 μL of purified DNA was transformed into 25 μL of Endura electrocompetent cells (Lucigen) following the manufacturer’s instructions with 2 mL of prewarmed recovery media. 2 mL of cells were plated onto a 245 mm x 245 mm plate and serially diluted cells were plated onto 100 mm diameter plates for the estimation of colony numbers. After overnight (15 hours) incubation at 30°C, colonies were collected, and plasmid pools were extracted with Maxiprep (Machery-Nagel). Colony coverage for 1 bioassay plate was at least 1100X for 155^2^ library members. Two separate maxi-prepped DNA solutions from two bioassay plates were mixed to achieve at least 2200X colony coverage of library members.

### High-throughput assay to measure transcriptional effects of Library 1

Low passage 293FT cells (Invitrogen) or Lenti-X (Takara) 293T cells were grown in DMEM (Invitrogen, 10566-016) supplemented with 10% FBS (Gibco) and 1%Penicillin-Streptomycin (Gibco). For one T225 flask, 39.9 μL of 1 mg/mL PEI MAX (Polysciences) and 1 mL of DMEM were mixed by inversion and incubated at room temperature for 10 min. During incubation, 5.6 μg of pMD2.G, 11.3 μg of psPAX2, and 22.7 μg of target plasmid or plasmid library were added to new 1 mL of DMEM and were thoroughly mixed by vortexing. After incubation, plasmids + DMEM mixture was added to PEI MAX + DMEM mixture followed by 30 sec vortexing. The final ∼2 mL mixture of plasmids + PEI MAX + DMEM was incubated at room temperature for 30 min. During incubation, 293FT or Lenti-X 293T cells were split by TrypLE (Gibco) and 33.4 million cells were added to a standing T225 flask in 5 mL of fresh growth media. After the 30 min incubation, plasmids + PEI MAX + DMEM mixture was added to the flask, and the flask was gently mixed by swirling. The flask was incubated for 5 min in the upright position. Following the 5 min incubation, fresh growth media was added up to 45 mL, and cells were incubated at 37°C, 5% CO_2_ overnight. The next day (Day 1), the media was changed to 45 mL of fresh growth media and incubated for an additional 2 days, after which the supernatant was collected and concentrated by Lenti-X concentrator (Takara) following the manufacturer’s protocol.

Upon infection of 40 million PB Tet-On dCas9 cells (without CD81 guide array) with 200 μL of lentivirus per replicate on Day -13, cells were cultured in growth media containing 1-2 μg/mL doxycycline (**Supplementary Fig. 5**). In parallel, 2 control cell lines (negative control: PB Tet-On dCas9 cells without any effector & positive control: PB Tet-On dCas9 cells with LV-pTRE3G-PuroR-T2A-stdMCP-eGFP-KRAB (ZNF10) preinstalled by lentiviral infection) started doxycycline incubation as well. Doxycycline was replaced or added daily, maintaining the concentration of 0.5-2 μg/mL to compensate for the short half of doxycycline in the media. 1 day post-infection (Day -12), puromycin selection started with a 1 μg/mL concentration. To simultaneously measure the lentiviral titer, subsets of cells were passaged to 96 well plates for the viability assay. 3 days after the start of puromycin selection (Day -9), lentiviral titer was measured by the CellTiter-Glo 2.0 Cell Viability Assay (Promega G9242) as previously described^84^. Measured viability was 0.25-0.29 suggesting successful low MOI infection. Puromycin selection was continued for an additional 3 days until Day -6. We observed an increase in the percentage of live cells by trypan blue staining (Thermo Fisher Scientific) during the additional 3 days of selection, suggesting the outgrowth of infected cells over debris from dead cells.

After a total of 6 days of puromycin selection and outgrowth (Day -6), 39.2 million surviving cells were mixed with 0.8 million positive control cells (LV-pTRE3G-PuroR-T2A-stdMCP-eGFP-KRAB (ZNF10)) per replicate. On Day -6, mixed cells were nucleofected with the CD81 targeting 3X guide array plasmid (pMH228). For each cuvette, 4 million cells were nucleofected with 4.5 μg of pMH228 following the manufacturer’s K562-specific protocol (4D X-unit, FF-120 program, Lonza). For each replicate, 40 million cells were nucleofected by 10 cuvettes. 4 million positive control cells were also nucleofected by 1 cuvette. 3 days post-nucleofection (Day -3), mCherry-positive cells were sorted. To reduce the variability generated by sgRNA expression level, a population expressing moderate levels of mCherry was sorted (**Supplementary Fig. 5b**). In total, ∼9.7 million cells (∼400X coverage) were sorted for each replicate. 3 days post-sorting (Day 0), at least 11.5 million cells per replicate were frozen for future FACS sorting. The remaining half of at least 11.5 million cells were centrifuged and passaged to media without doxycycline to stop the expression of dCas9 and the stdMCP-combinatorial effectors. Cells were continuously expanded until Day 6 in culture media without doxycycline. On Day 6, 50 million cells were frozen for each replicate. Cells were again continuously expanded until Day 12. On Day 12, 120 million cells were frozen for each replicate.

On the day of FACS, frozen cells from various timepoints (Day 0, 6, and 12) were thawed briefly and washed twice with eBioscience Flow Cytometry Staining Buffer (Invitrogen). APC Mouse Anti-Human CD81 antibody (Cat# 561958, BD Bioscience) was diluted 1:10 in 100 μL stain buffer per 0.1 million cells. Staining was performed on a 3D rotary mixer in a cold room (4°C) for 1 hour. Cells were washed twice with stain buffer and sorted using a FACSAria Fusion Special Order Research Product (BD Bioscience). mCherry-negative singlet populations were separated into two populations based on CD81 level (**Supplementary Fig. 5c**). Sorted cells were frozen at -80°C until genomic DNA extraction.

Genomic DNA (gDNA) was extracted using columns from the Quick-DNA Miniprep Plus kit (Zymo Research) and reagents from the Quick-DNA Midiprep Plus kit (Zymo Research) with the following modifications in the protocol. Cells were split into multiple columns if the number of cells was larger than 3 million. Per 3 million cells, excess genomic lysis buffer (Beta-Mercaptoethanol added) with a volume of 1.6 mL from the Quick-DNA Midiprep Plus kit (Zymo Research) was used instead of BioFluid & Cell Buffer from the Quick-DNA Miniprep Plus kit (Zymo Research). After the addition of 1.6 mL genomic lysis buffer to the cells, the mixture was vortexed thoroughly. Cells were lysed in genomic lysis buffer for 1 hour at room temperature with 3D rotation. Excess genomic lysis buffer ensured complete lysis of the cells, yielding a high amount of extracted gDNA. Except for the described cell lysis step, we followed the manufacturer’s protocol for the Quick-DNA Miniprep Plus kit (Zymo Research).

Enriched combinatorial library members were PCR amplified from the gDNA using the following conditions. For one 50 μL of the PCR reaction, 25 μL of 2X NEBNext HiFi (NEB), 1 μg of gDNA, and a final concentration of 0.5 μM for each forward and reverse NGS primers (**Supplementary Table 8**) were prepared and mixed on ice. One-step NGS primers were designed to have 0∼7 random spacer nucleotides to increase the base diversity during the initial sequencing cycles^85^. The thermocycling protocol was 1 cycle of 98°C for 3 min, 27 cycles of 98°C for 10 sec, 64°C for 30 sec, 72°C for 75 sec, and final 1 cycle of 75°C for 2 min. After thermocycling, PCR reactions were pooled and run through multiple lanes in 1.2% TBE gel. Rectangular regions of the gel spanning from 800 bp ∼ 2 kb were excised, and PCR products were extracted using Monarch DNA Gel Extraction Kit (NEB). The gel-extracted library was quantified by Nanodrop (Thermo Fisher Scientific), Qubit HS kit (Thermo Fisher Scientific), Bioanalyzer (Agilent Technologies), and KAPA Library Quantification Kit (Roche). We noticed approximate agreement in quantification between Qubit and Bioanalyzer while KAPA library quantification exhibited the most discrepancy, potentially due to differences in the size between our library and the standards provided by the kit. Based on the results of Qubit and Bioanalyzer, the library was pooled with ∼10% PhiX control (Illumina) and sequenced on a NextSeq2000 (Illumina) with 2 x 300 bp P2 kits (Illumina). In total, two 2 x 300 bp P2 kits were used. Libraries from Day 0 and 6 were pooled together and sequenced with one kit, and the Day 12 library was sequenced separately using the second kit.

### NGS analysis of bivalent domains from Library 1

To address the issue of lower R2 sequencing quality, which is occasionally observed during paired-end readout of long fragments^86^, we implemented an approach of spreading ten 15-mer short identifiers (IDs) across the entire 301 bp sequencing region. If any of the 10 IDs exactly matched the NGS read, the read was assigned to the corresponding domain. Unique 15 mer IDs for identifying N- and C-terminal domains were generated by the following steps.

Step 1) From fasta files composed of expected 301 bp NGS reads for every domain (R1_Forward_P5_301cycles.fasta & R2_Reverse_P7_301cycles.fasta), all possible unique 15 mers were generated using UniqueKMER with the following commands^87^.

$ uniquekmer -f R1_Forward_P5_301cycles.fasta -k 15 -o “kmercollection_R1“
$ uniquekmer -f R2_Reverse_P7_301cycles.fasta -k 15 -o “kmercollection_R2“

Step 2) If a 15 mer ID for one domain had a hamming distance of less than 2 with an ID from another domain, one of the IDs was deleted from the domain with a larger number of IDs by executing Library1_step2.py. Output files were numerically sorted using Seqkit with the following commands^88^.

$ seqkit sort -nN R1_15mer_HD2_UMIs.fasta -o R1_15mer_HD2_UMIs_sorted.fasta
$ seqkit sort -nN R2_15mer_HD2_UMIs.fasta -o R2_15mer_HD2_UMIs_sorted.fasta

Step 3) Using pairwise2 (Biopython) local alignment mode with the scoring scheme of (match: 1, mismatch: 0, opening gap: -1, extending gap: -1), IDs that had an alignment score higher than 12 with any other domains’ expected 301 bp NGS reads were removed. Also, 10 IDs were chosen to be spread across the 301 bp region as evenly as possible by executing Library1_step3.py.

Step 4) For domains that ended up with fewer than 10 IDs, previously removed IDs with alignment scores (pairwise2) of 13 were recalled and assigned to ensure that the total number of IDs reached 10 for every domain by executing Library1_step4.py. Finally, 10 IDs (R1_15mer_UMIs_10per_effector.fasta and R2_15mer_UMIs_10per_effector.fasta) were generated for each domain.

By matching R1 and R2 reads, a count table for bivalent candidates was generated. We noticed that there was a trend of count numbers for homo-bivalent candidates being higher than hetero-bivalent candidates which might be caused by recombination artifacts. Due to the difficulty in discerning whether the high count number arises from transcriptional perturbation of CD81 or from recombination artifacts, we opted not to analyze homo-bivalent candidates. If the sum of NGS reads from High and Low bins was less than 40 in at least one replicate, the bivalent candidate was considered a dropout. The ratio of a bivalent candidate in a specific bin was calculated by the following formula.

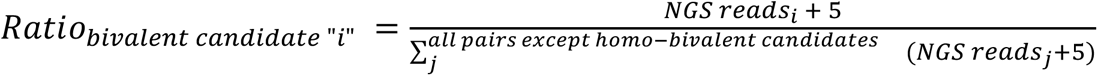

The enrichment score of bivalent candidate “i” was calculated by dividing Ratio_i,CD81 High bin_ by Ratio_i,CD81 Low bin_.

### Secondary validation of individual and bivalent effectors from Library 1

To harness previously ordered effector gene fragments, we again generated a series of landing pad plasmids for nucleofection-based delivery of effectors. For bivalent domain candidates, similar to the library cloning procedure, 3 sequential golden gate reactions were performed. The landing pad plasmid for the 1st golden gate reaction was the same KanR1_receiver (pCM001) as the library cloning, whereas KanR2_individual_validation_receiver (pCM023) was used as the landing pad plasmid for the 2nd golden gate reaction. For the final golden gate reaction, the landing pad was pCAG_AmpR_backbone (pCM024), which enables the constitutive expression of the effector by the CAG promoter. For monovalent domain plasmid cloning, 2 sequential golden gate reactions were performed. For the 1st golden gate reaction, the landing pad plasmid was KanR_monovalent_receiver (pCM025). For the 2nd golden gate reaction, the landing pad plasmid was the same as the bivalent domain candidate cloning, pCAG_AmpR_backbone (pCM024).

For cloning linear gene fragments into KanR1_receiver (pCM001), KanR2_individual_validation_receiver (pCM023), and KanR_monovalent_receiver (pCM025), 0.25 μL of 60 ng/μL landing pad plasmids, 2.5 μL of 20 nM gene fragment, 0.5 μL of 10X T4 ligase buffer (NEB), 0.25 μL BsaI-HFv2 (NEB, 20000 U/mL), 0.25 μL T4 Ligase (NEB, 400000 U/mL) and 1.25 μL nuclease-free water (Ambion) was mixed well resulting in a total of 5 μL reaction. The thermocycling condition was 30 cycles of 37°C for 1 min and 16°C for 1 min followed by 1 cycle of 37°C for 5 min and 75°C for 5 min. For the final golden gate reaction, 0.5 μL of 100 ng/μL pCAG_AmpR_backbone (pCM024), 1 μL of 50 ng/μL N-terminus effector plasmid, 1 μL of 50 ng/μL C-terminus effector plasmid, 1 μL of FastDigest Esp3I (Thermo Fisher Scientific), 0.5 μL of T4 DNA ligase (NEB, 400000 U/mL), 1 μL of 10X T4 buffer (NEB), and 5 μL of nuclease-free water (Ambion) were mixed well resulting in a total of 10 μL reaction. KanR_monovalent_receiver (pCM025) was used instead of KanR1_receiver (pCM001) or KanR2_individual_validation_receiver (pCM023) for cloning the monovalent effector plasmid. The thermocycling protocol was the same: 30 cycles of 37°C for 1 min and 16°C for 1 min followed by 1 cycle of 37°C for 5 min and 75°C for 5 min.

Nucleofection of effector plasmids was performed with SF Cell Line 4D-Nucleofector X Kit S following the manufacturer’s protocol (4D X-unit, FF-120 program, Lonza). PB Tet-On dCas9 K562 cell line (with CD81 guide array) was preinduced with 1-2 μg/mL doxycycline starting 1-3 days before the day of nucleofection. For complete details refer to **Supplementary Table 9**. On the day of measurement, ∼300 μL of cells were washed twice with eBioscience Flow Cytometry Staining Buffer, and 99 μL staining buffer with 1 μL of APC Mouse Anti-Human CD81 antibody (Cat# 561958, BD Bioscience) was used per well in 96 well plates. Following 1 hour of incubation at 4°C, cells were washed twice, and the CD81 level was measured by Attune NxT Flow Cytometer (Thermo Fisher Scientific). Empty vector pUC19 nucleofected cells were used as control cells for gating on BFP, GFP, mCherry, and CD81 (APC) levels.

### Library 2 design, cloning, and barcode mapping

A literature review was conducted to identify minimally active catalytic epigenetic effectors across 11 well-established classes: DNA demethylation machinery (DNDM), DNA methyltransferase (DNMT), E2 and E3 ubiquitin ligases (UBL), histone acetyltransferase (HAT), histone arginine methyltransferase (HRMT), histone deacetylase (HDAC), histone demethylase (HDM), histone deubiquitinase (DUB), histone kinase (HK), histone lysine methyltransferase (HKMT), and histone phosphatase (HP). Human proteins associated with each class were identified using Gene Ontology terms, except for HKMTs, which were identified as human proteins containing SET domains in addition to GO terms. Within each list, each protein was manually reviewed to assess for possible catalytic activity and to search for catalytically active truncations. In total, 195 catalytic domains were identified across all classes, with approximately 75% sourced directly from literature, while the remaining domain bounds were sourced from alignment with closely related proteins or simply estimated from Pfam annotations. Additionally, three non-catalytically active controls were included: RYBP for silencing^37^, a 400 aa fragment of DMD determined to be neutral from HT-recruit^24^, and HSF1 for activation^10^. All domains and references are included in **Supplementary Table 2**.

Amino acid sequences for each effector were codon optimized for human expression using DNA Chisel with constraints to avoid restriction enzyme sites (BsaI, BsmBI, BbsI, BtgZI), avoid patterns (5 x 3-mer, 2 x 15-mer, 2 x 12-mer, 8 x homopolymer), and enforce GC content between 35-65% in each 50 bp window^82^. Each effector was ordered from Twist Biosciences as a clonal gene in pTwist Kan High Copy with flanking sequences containing nested BsmBI and BsaI golden gate sites to facilitate cloning bivalent combinations (**Supplementary Fig. 3a and Supplementary Table 8**, Misc., 5′ and 3′ Effector Flank). All golden gate overhangs were sourced from a 15 member high fidelity set, and all landing pads contained CcdB dropout genes to improve cloning efficiency^83^. Individual effector plasmids were transformed into MACH1 cells (Thermo Fisher Scientific) and miniprepped using QIAprep Spin Miniprep Kit (Qiagen, 27106). Because lentiviral transduction efficiency decreases roughly 3.5X per additional kilobase of DNA^30^, effector plasmids were pooled exponentially by effector size by manually adding 0.2*3.5^(x/1000) pmol of plasmid DNA, where x is the length of the effector in basepairs (**Supplementary Fig. 3b,c**). The BsmBI sites in this pool (EF2 pool) were later used to clone into the N-terminal Effector 2 position, and the BsaI sites were used to subclone the pooled effectors into a new high copy Chloramphenicol resistant backbone via golden gate reaction (50 ng pMH249 (**Supplementary Table 8**, Plasmids), 2-fold molar excess EF2 pool, 1 μL BsaI-HFv2 (NEB, R3733L), 1 μL T4 ligase (NEB, R0202L), 2.5 μL T4 ligase buffer (NEB, B0202S), and H_2_O to 25 μL incubated at 37°C for 5 min, followed by 30 cycles of 37°C for 5 min and 16°C for 10 min, followed by final ligation at 16°C for 20 min, final digestion at 37°C for 30 min, and heat inactivation at 80°C for 20 min) (**Supplementary Fig. 3d**). The reaction was then cleaned using DNA Clean & Concentrator-5 (Zymo, D4013) and eluted in 6 μL H_2_O, before electroporating 2 μL into 25 μL of Endura electrocompetent cells following the manufacturer’s instructions (Biosearch Technologies, 60242-1). Recovered cells were plated onto a 245 mm x 245 mm plate and incubated overnight at 37°C. The next day, colonies were scraped from the plate and the plasmid pool was maxiprepped (Machery-Nagel, 740424.50). The resulting EF1 pool contained BsmBI golden gate sites at the same 4 bases as the BsaI insertion of the effector to allow for subsequent cloning into the C-terminal Effector 1 position (**Supplementary Fig. 3e**).

Before the final bivalent assembly of the EF1 and EF2 pool into our lentiviral landing pad, the XTEN16 linker and variable barcodes were prepared. The XTEN16 linker was ordered as two ssDNA oligos with the correct overhangs for assembly between the EF1 and EF2 position and annealed by equimolar mixing at room temperature (**Supplementary Fig. 3f and Supplementary Table 8**, Misc.). Variable barcodes were ordered from Integrated DNA Technologies (IDT) as a 20xN ssDNA oligo flanked by NGS primer binding sites and BsmBI golden gate sites for assembly after EF2 (**Supplementary Fig. 3g and Supplementary Table 8**, Misc.). The oligo was ordered preannealed with the reverse complement of the 3′ flanking sequence to facilitate filling in of the bottom strand of the barcode and 5′ flanking sequence by polymerase extension (2.5 μL of preannealed oligos at 10 uM, 12.5 μL KAPA HiFi HotStart ReadyMix (Roche KK2601), and H_2_O to 25 μL followed by incubation at 98°C for 3 min and 72°C for 30 min). The extended barcode was cleaned with DNA Clean & Concentrator-5 (Zymo, D4013) and eluted in 20 μL H_2_O, before predigestion (1 μg extended barcode, 2 μL Esp3I (Thermo Fisher Scientific, FD0454), 2 μL 10X FastDigest buffer (Thermo Fisher Scientific, B64), and H_2_O to 20 μL incubated at 37°C for 25 min). The digested barcode was cleaned with DNA Clean & Concentrator-5 (Zymo, D4013) and diluted to 2 ng/μL. Final bivalent assembly was performed via 5-piece golden gate reaction into our lentiviral landing pad, which was cloned in part using the EMMA toolkit^81^, containing pTRE3G-MCP-XTEN80 and pSV40-PuroR cassettes (250 ng pMH276 (**Supplementary Table 8**, Plasmids), 1X molar EF1 pool, 1X molar annealed XTEN16 linker, 1X molar EF2 pool, 1X molar digested barcode, 1 μL Esp3I (Thermo Scientific, FD0454), 1 μL T4 ligase (NEB, R0202L), 2.5 μL T4 ligase buffer (NEB, B0202S), and H_2_O to 25 μL incubated at 37°C for 5 min, followed by 30 cycles of 37°C for 5 min and 16°C for 10 min, followed by final ligation at 16°C for 20 min, final digestion at 37°C for 30 min, and heat inactivation at 80°C for 20 min) (**Supplementary Fig. 3h**). The reaction was then cleaned using DNA Clean & Concentrator-5 (Zymo, D4013) and eluted in 6 μL H_2_O, before electroporating 2x 2 μL into 25 μL of Endura electrocompetent cells each following the manufacturer’s instructions (Biosearch Technologies, 60242-1). Recovered cells were serially diluted and plated onto 100 mm diameter plates for the estimation of colony numbers, while the remaining cells were plated onto several 245 mm x 245 mm plates. After overnight incubation at 30°C, colonies on dilution plates were counted, and enough large plates were scraped and maxiprepped (Machery-Nagel, 740424.50) such that the average coverage was between 100-150X colonies per member. Strict coverage constraints were necessary both to adequately represent shorter effector pairs and to ensure that there were few enough barcodes to map via long-read sequencing.

Long-read nanopore sequencing was used to map barcodes to bivalent effector pairs. The plasmid pool was prepared for sequencing using two Cas9 digestion steps followed by nanopore adapter ligation in an attempt to bias reads towards the barcoded bivalent effector region (**Supplementary Fig. 3i**). First, two Alt-R *S.p.* Cas9 (IDT, 1081060) RNP pools were made, one containing the outer guides M1 and P1 and the other containing the inner guides M2 and P2, by prepooling 1 μL of each 100 μM crRNA and following the Alt-R CRISPR-Cas9 System protocol (**Supplementary Table 8**, Guides). The plasmid pool was then digested with the outer guides M1 and P1 (5 μg plasmid pool, 10 μL M1+P1 Cas9 RNP, 4 μL 10X rCutSmart buffer (NEB, B6004S), and H_2_O to 40 μL incubated at 37°C for 1 hour). The reaction was stopped by the addition of Proteinase K (+5 μL 20 mg/mL Proteinase K (Zymo, D3001-2-20), incubated at 56°C for 10 min). A 1X AMPure XP bead (Beckman Coulter, A63881) cleanup was performed before dephosphorylation of ends followed by second digestion with the inner guides M2 and P2 (24 μL eluate, 3 μL Quick CIP (NEB, M0525S), and 3 μL 10X rCutSmart buffer (NEB, B6004S) incubated at 37°C for 10 min and heat inactivated at 80°C for 2 min, followed by addition of 10 μL M2+P2 Cas9 RNP and incubation at 37°C for 1 hour). The reaction was stopped by the addition of Proteinase K (+5 μL 20 mg/mL Proteinase K (Zymo, D3001-2-20), incubated at 56°C for 10 min). A 0.5X AMPure XP bead (Beckman Coulter, A63881) cleanup was performed before dA tailing using Taq polymerase (34 μL eluate, 1 μL 10 mM dNTP mix (NEB, N0447S), 1 μL Taq polymerase (NEB, M0267S), and 4 μL 10X rCutSmart buffer (NEB, B6004S) incubated at 72°C for 5 min). Another 0.5X AMPure XP bead (Beckman Coulter, A63881) cleanup was performed before nanopore adapter ligation (60 μL eluate, 25 μL Ligation Buffer (LNB, Oxford Nanopore, SQK-LSK114), 10 μL NEBNext Quick T4 DNA Ligase (NEB, E6056S), and 5 μL Ligation Adapter (LA, Oxford Nanopore, SQK-LSK114) incubated at room temperature for 20 min). A final AMPure XP bead (Beckman Coulter, A63881) cleanup was performed following the SQK-LSK114 protocol using the Short Fragment Buffer (SFB, Oxford Nanopore, SQK-LSK114). The resulting library was quantified using Qubit 1X dsDNA High Sensitivity Assay (Invitrogen, Q33231) and diluted to 20 fmol in 12 μL elution buffer (EB, Oxford Nanopore, SQK-LSK114) for sequencing on a MinION flow cell (Oxford Nanopore, FLO-MIN114) followed by a PromethION flow cell (Oxford Nanopore, FLO-PRO114M).

Nanopore reads were basecalled and duplexed using Dorado’s super-accuracy model v4.1.0 before conversion to .fastq file format by executing supduplex2fastq.sh on an appropriately configured GPU virtual machine. Barcode to effector pairings were then mapped by executing bc_extraction.py. Briefly, Effector 1, Effector 2, and barcode sequences were extracted from each read using cutadapt and appropriate flanking sequences before effectors were mapped to individual effectors using minimap2^89,90^. Effector mappings with <75% identity were discarded. The mapped barcodes from all sequencing runs were then combined into a single table of unique pairings and filtered to remove any duplicate barcodes (mappedBCs_filt.csv). In total, 7.8 million barcodes were mapped and represented every possible bivalent combination with the number of barcodes mapped scaling exponentially by bivalent effector length (**Supplementary Fig. 3j,k**).

### High-throughput assay to measure transcriptional effects of Library 2

A pooled lentiviral library was generated from Library 2 using the LV-MAX kit (Gibco, A35348) and protocol in a 1L flask. Lentivirus was concentrated 10X using Lenti-X concentrator following the manufacturer’s instructions (Takara, 631232). Functional titer was measured by transducing 40,000 K562 cells per well of a 96 well plate with serial dilutions of lentivirus and 8 μg/mL polybrene (Millipore, TR-1003-G) via 90 minute spinfection at 1000g and 32°C. After overnight incubation, media containing lentivirus was replaced with fresh media and cells were allowed to recover for another 24 hours before being split equally into media with and without puromycin (2 μg/mL final, Gibco, A1113803). After five days of selection, cell survival in each well was quantified on a Tecan Spark plate reader using CellTiter-Glo 2.0 Cell Viability Assay (Promega, G9242). Percent survival was calculated as the ratio of luminescence in the presence versus absence of puromycin for each lentiviral dilution, and functional lentiviral titer was calculated and averaged for all dilutions with 5-30% survival. Two independent full scale transductions were then performed by transducing the PB Tet-On dCas9 3X CD81 K562 cell line at 100X library coverage and an MOI <0.1 with 8 μg/mL polybrene (Millipore, TR-1003-G) via 90 minute spinfection at 1000g and 32°C. After overnight incubation, media containing lentivirus was replaced with fresh media and cells were allowed to recover for another 24 hours before starting selection with 2 μg/mL puromycin for 5 days (Gibco, A1113803).

To assess bivalent chromatin effector activity in high-throughput, engineered cells were induced with 1 μg/mL doxycycline in 200 mL media in a 1L Erlenmeyer flask, maintaining >1000X library coverage and replacing doxycycline media each day. After 5 days of induction (Day 0 timepoint), roughly 150 million cells (∼3750X coverage) were frozen in 10 mL media supplemented with 10% DMSO at -80°C in a 15 mL falcon tube. Roughly 50 million remaining cells (∼1250X coverage) were passaged to assess memory by washing twice with DPBS (Gibco 14190144) and resuspending in 400 mL fresh media without doxycycline in a 2L Erlenmeyer flask. Every 3 days for 24 total days, two frozen stocks of ∼3000X cells were made as described above, and cells were diluted to a minimum density of 1e5/mL in 400 mL media in a 2L Erlenmeyer flask. Two frozen timepoints, Day 0 and Day 12, were chosen to assess transient effector activity and epigenetic memory on CD81 (**Supplementary Fig. 6a**). For a given timepoint and replicate, the frozen aliquot was thawed in a 37°C water bath for 4 min before being diluted in 35 mL stain buffer (BD, 554656) and pelleted at 300g for 10 min. The cell pellet was resuspended in 14 mL stain buffer containing 750 μL PE-conjugated CD81 monoclonal antibody (Invitrogen, MA1-10292), and the tube was rotated for 30 min at 4°C. After staining, cells were spun down at 300g for 8 min and washed twice with 15 mL stain buffer before resuspending in 15 mL and straining through a 40 μM filter. Using a BD FACSAria Fusion Special Order Research Product, between 7-13 million cells were sorted from the top and bottom 25% of the PE fluorescence distribution (**Supplementary Fig. 8b**-d).

To measure bivalent effector representation in each sorted population, genomic DNA was harvested using NucleoSpin Blood L kit and protocol (Machery-Nagel, 740954.20), adding 200 μL Monarch RNAse A (NEB, T3018L) before lysis and using high-yield elution recommendations. Then NGS libraries of bivalent effector barcodes were prepared using two sequential PCR steps. In PCR1, barcodes were amplified from gDNA using primer pairs containing staggers to diversity NGS reads (**Supplementary Table 8**, Primers, MH160-MH175) (up to 100 μg gDNA, 24 μL PCR1 primer set at 50 μM each, 600 μL NEBNext Ultra II Q5 MM (NEB, M0544L), and H_2_O to 1200 μL, divided into 12× 100 μL reactions in a PCR plate and incubated at 98°C for 30 sec, 20 cycles of 98°C for 10 sec and 65°C for 75 sec, final extension at 65°C for 5 min). Split reactions were pooled and mixed thoroughly before 25 μL were cleaned using a two sided AMPure XP bead (Beckman Coulter, A63881) cleanup from 0.7X to 1.8X. In PCR2, sample indices were added along with flow cell binding sequences p5 and p7 (50-100 ng purified PCR1, 5 μL PCR2 primer set at 10 μM each, 25 μL NEBNext Ultra II Q5 MM (NEB, M0544L), and H_2_O to 50 μL, incubated at 98°C for 30 sec, 7 cycles of 98°C for 10 sec and 65°C for 75 sec, final extension at 65°C for 5 min). The reactions were cleaned using a two sided AMPure XP bead (Beckman Coulter, A63881) cleanup from 0.7X to 1.5X. Each sample was quantified using Qubit 1X dsDNA High Sensitivity Assay (Invitrogen, Q33231) and 60 ng of each were pooled into a single NGS library that was sequenced on a NextSeq 2000 using a P3 2×51 cycle paired-end run (Illumina, 20040559).

### Targeted nanopore sequencing to assess barcode swapping in Library 2

To address the issue of barcode swapping, which is commonly associated with the use of barcodes in lentiviral libraries^91^, Library 2 contained only a short 39 bp homologous sequence between the end of the effector fusion and the barcode. Still, to quantitatively measure lentiviral recombination between bivalent effector pairings and barcodes, barcode swapping was assessed post-integration via long-read nanopore sequencing using Cas9-guided adapter ligation (**Supplementary Fig. 7a**)^92^. Genomic DNA from one full-scale lentiviral infection replicate was extracted using Quick-DNA Midiprep Plus kit and protocol (Zymo, D4075). An Alt-R *S.p.* Cas9 (IDT, 1081060) RNP pool was made by prepooling 1 μL of crRNAs M1, M2, P1, and P2 at 100 μM and following the Alt-R CRISPR-Cas9 System protocol (**Supplementary Table 8**, Guides). Genomic DNA was dephosphorylated before Cas9 cleavage and dA-tailing (5 μg gDNA, 3 μL Quick CIP (NEB, M0525S), 3 μL 10X rCutSmart buffer (NEB, B6004S), H_2_O to 30 μL incubated at 37°C for 10 min and heat inactivated at 80°C for 2 min, followed by addition of 10 μL M1+M2+P1+P2 Cas9 RNP, 1 μL dATP at 10 mM (NEB, N0440S), and 1 μL Taq polymerase (NEB, M0267S) and incubation at 37°C for 20 min and 72°C for 5 min). The reaction was cleaned using AMPure XP beads (Beckman Coulter, A63881) at 0.5X and DNA eluted in 60 μL H_2_O. Nanopore adapters were ligated following the SQK-LSK114 protocol, and the resulting library was sequenced on a MinION flow cell (Oxford Nanopore, FLO-MIN114).

Nanopore reads were basecalled and converted to .fastq as previously described. Sequences containing complete reads through both effector positions were extracted using cutadapt and appropriate flanking sequences^90^. Barcode to effector pairings were mapped using bc_extraction.py, modified to retain all found barcodes, which were then referenced against mappedBCs_filt.csv. Reads from any mismatches between barcode to effector pairings were manually examined to determine the mismatch position (**Supplementary Fig. 7b**).

### NGS analysis of barcodes and bivalent domains from Library 2

NGS reads were converted into barcode counts using the script count_barcodes.py. In summary, reads were merged with fastp before barcodes were excised with cutadapt using forward and reverse NGS primer sequences^89,90^. Then, barcodes were dereplicated and tabulated using SeqFu, and the results were combined into a single table of barcode occurrences by experimental condition^93^. After counting, any barcodes that were not mapped to effector pairs via nanopore were discarded, leaving an average of 76.6% of NGS reads (**Supplementary Fig. 6e**), and any barcodes containing homopolymer repeats of 8 or more were discarded due to limitations of nanopore sequencing. Next, barcode counts mapping to the same effector pair were combined to produce a total read count for each bivalent effector in each condition. Effector pairs were considered dropouts if there were less than 100 cumulative reads in both bins in either replicate and/or less than 5 reads in any bin of either replicate to prevent artifactual inflation or deflation of enrichment scores. Then read counts were normalized by the sum of each bin, and the enrichment scores were calculated as the ratio of the normalized counts in the High bin versus the Low bin. The average enrichment score from both replicates was computed using a geometric mean of the High/Low enrichment ratio.

### Secondary validation of individual and bivalent effectors from Library 2

Selected individual and bivalent effectors from Library 2 were individually assessed via transient transfection into the PB Tet-On dCas9 K562 cell line with the 3X CD81 guide array (for assessing activity on CD81) or without the 3X CD81 guide array (for assessing activity on additional genes). To clone the constructs for transient transfection, a CcdB dropout landing pad was first made to assemble the pTRE3G-MCP-XTEN80-effector fusion while also expressing pEF1a-EGFP-P2A-PuroR in a divergent orientation (**Supplementary Table 8**, Plasmids, pMH308). The individual clonal effectors ordered from Twist Biosciences were directly used for insertion into the Effector 2 position via the BsmBI sites in the flanking sequences. To generate the proper overhangs for insertion into the Effector 1 position, clonal effectors were predigested with BsaI (600 ng clonal fragment, 1 μL BsaI-HFv2 (NEB, R3733L), 1 μL Quick CIP (NEB, M0525S), 1 μL 10X rCutSmart buffer (NEB, B6004S), and H_2_O to 10 μL incubated at 37°C for 2 hours and 80°C for 1 hour). The same XTEN16 linker was used for bivalent assembly, while an Effector 2 stuffer was used in place of both Effector 2 and the XTEN16 linker for monovalent assembly (**Supplementary Table 8**, Misc.). To prepare the XTEN16 linker and Effector 2 stuffer, individual ssDNA oligos were ordered, annealed, and phosphorylated (1 μL 100 μM F oligo, 1 μL 100 μM R oligo, 1 μL T4 ligase buffer (NEB, B0202S), 1 μL T4 PNK (NEB, M0201S), and H_2_O to 6 μL incubated at 37°C for 30 min, 65°C for 20 min, and 95°C for 5 min before cooling down to 4°C over 15 min). The final assembly of bivalent effectors was performed using Golden Gate assembly (50 ng pMH308, 0.5 μL EF1 predigest reaction, 0.2 μL XTEN16 oligo anneal, 0.5 μL EF2 clonal fragment at ∼60 ng/μL, 1 μL T4 ligase buffer (NEB, B0202S), 0.5 μL Esp3I (Thermo Fisher Scientific, FD0454), 0.5 μL T4 ligase (NEB, R0202L), and H_2_O to 10 μL incubated at 37°C for 5 min, followed by 30 cycles of 37°C for 5 min and 16°C for 10 min, followed by final ligation at 16°C for 20 min, final digestion at 37°C for 30 min, and heat inactivation at 80°C for 20 min). The final assembly of monovalent effectors was also performed using golden gate assembly (50 ng pMH308, 0.5 μL EF1 predigest reaction, 0.2 μL EF2 stuffer oligo anneal, 1 μL T4 ligase buffer (NEB, B0202S), 0.5 μL Esp3I (Thermo Scientific, FD0454), 0.5 μL T4 ligase (NEB, R0202L), and H_2_O to 10 μL incubated at 37°C for 5 min, followed by 30 cycles of 37°C for 5 min and 16°C for 10 min, followed by final ligation at 16°C for 20 min, final digestion at 37°C for 30 min, and heat inactivation at 80°C for 20 min). 2 μL of each reaction were transformed into MACH1 cells (Thermo Fisher Scientific) before individual colonies were picked and grown overnight in 3 mL TB for plasmid purification with NucleoSpin Plasmid Transfection-grade miniprep kit (Machery-Nagel, 740490). All plasmids were sequence confirmed via full plasmid nanopore sequencing.

Catalytic mutations for specific effectors were sourced through literature review (**Supplementary Table 5**). To clone mutations into sequence confirmed individual validation constructs generated above, 2-3 PCR fragments were made to overlap over the mutation site or sites and the GFP gene for Gibson assembly of the mutant plasmids (**Supplementary Table 8**, Primers, MH302-MH345). PCR fragments were generated from corresponding unmutated plasmids (10 ng parent plasmid, 0.75 μL 10 μM F primer, 0.75 μL 10 μM R primer, 12.5 μL KAPA HiFi HotStart ReadyMix (Roche, KK2601), H_2_O to 10 μL incubated at 95°C for 3 min, 18 cycles of 98°C for 20 sec and 72°C for 3 min, and 72°C for 6 min, followed by addition of 1 μL DpnI (NEB, R0176S) and incubation at 37°C for 1 hour and 80°C for 20 min). 2 μL of the resulting reactions were analyzed for product length and purity via gel electrophoresis before mutant plasmids were assembled via Gibson assembly (1 μL each corresponding PCR reaction, 10 μL 2X Gibson MM, and H_2_O to 10 μL incubated at 50°C for 1 hour). 2 μL of each reaction were transformed into MACH1 cells (Thermo Fisher Scientific) before individual colonies were picked and grown overnight in 3 mL TB for plasmid purification with NucleoSpin Plasmid Transfection-grade miniprep kit (Machery-Nagel, 740490). All mutant plasmids were sequence confirmed via full plasmid nanopore sequencing.

Sequence confirmed validation plasmids were nucleofected into the PB Tet-On dCas9 K562 cell lines using SF Cell Line 96-well Nucleofector kit (Lonza, V4SC-2096). The manufacturer’s instructions were followed using 800 ng plasmid DNA per nucleofection. For testing effectors on CD81, only the effector plasmid was nucleofected into the Tet-On dCas9 cell line with the 3X CD81 guide array. For testing effectors on additional genes, the effector plasmid and the corresponding 3X guide array plasmid were nucleofected at a 1:1 mass ratio into the Tet-On dCas9 cell line without the 3X CD81 guide array. After nucleofection, cells were plated directly into 1 μg/mL final concentration of doxycycline to induce dCas9 and MCP-effector expression. One day after nucleofection and every subsequent day until 5 days post-nucleofection, media was replaced with fresh media containing 1 μg/mL doxycycline and 2 μg/mL puromycin (Gibco, A1113803). Five days post-nucleofection, cells were analyzed for transient effector activity on the gene target by transferring to 96 well U-bottom plates and spinning down at 300g for 3 min. Media was aspirated, and cells were resuspended in 100 μL stain buffer (BD, 554656) per well. Then 100 μL stain buffer containing 1.25 μL fluorescently conjugated antibody was added to each well and mixed by pipetting before incubation at 4°C for 30 min (APC CD55 antibody (Biolegends, 311311), Alexa Fluor 647 CD58 antibody (BD Pharmingen, 563567), PE CD81 antibody (Invitrogen, MA1-10292), APC CD151 antibody (Biolegends, 350405), and APC CD155 antibody (eBioscience, 2H7CD155). After incubation, cells were washed twice with 200 μL stain buffer before final resuspension in 200 μL stain buffer per well. Stained cells were analyzed by flow cytometry on an Attune NxT Flow Cytometer (Thermo Fisher Scientific). To assess for long-term effector activity, cells at five days post-nucleofection were transferred to 96 well V-bottom plates and washed twice with 200 μL DPBS (Gibco, 14190144) per well before resuspending in fresh media without doxycycline or puromycin. Three days later and at subsequent timepoints, cells were split to maintain growth, and the remaining cells were stained and analyzed as above.

To systematically analyze large batches of flow cytometry data, Cytoflow’s Jupyter notebook integration was used^94^. Live and singlet gates were first defined on WT K562 cells measured using the same cytometer settings, and the gates were applied to each sample file. Then, for transient timepoint analysis, cells were further gated for BFP and GFP, indicating dCas9 expression and the presence of the MCP-effector plasmid. Transient timepoints on genes other than CD81 were also gated on mCherry, indicating the presence of the 3X guide array plasmid. Threshold gates were defined at the 99.9th percentile of the channel measurements in WT K562 cells. For memory timepoints, no gating beyond live and singlet gates was performed. After gating, mean fluorescence intensity (MFI) was calculated for each sample using the flow.geom_mean function in Cytoflow on the channel corresponding to the fluorescent antibody. Percent change in MFI was calculated using the MFI from the neutral DMD control as a reference. Percent cells activated and percent cells repressed were calculated using threshold gating at the 1st and 99th percentile of the neutral DMD control effector, and the percent cells silenced was calculated using threshold gating at the 99th percentile of unstained WT K562 cells.

### Analysis of individual and bivalent effector strength

To calculate the total number of significant effector pairs for each library, the raw count matrices were analyzed with pyDESeq2^95^ which corresponds to DESeq2 (v1.34.0) with single-factor analysis, Wald tests, and LFC shrinkage mode (**Supplementary Tables 3** and **4**). For screen 1, candidates with total NGS reads of at least 80 were analyzed. For screen 2, the total NGS reads threshold was set as 100. Threshold for statistical significance was set as l log2FC l > 1 and p_adjusted_ < 0.05.

A marginal effect test was defined to assess the marginal, or average, effect of a given individual effector. To conduct the test for a given effector, the bivalent pairs and corresponding average fold enrichment scores were split into two groups, those containing the effector and those not containing the effector. The marginal score was defined as the difference in mean fold enrichment between the two groups, and if ≥ 30 pairs were present in both groups, a Welsh’s t-test was conducted. It was determined that a difference of means t-test was necessary, rather than a one sample t-test, to correct for library composition bias. In other words, the mean fold enrichment of any given individual effector would naturally be positive or negative if the overall library had a bias in either direction simply because the given effector is paired with every other library member. A Welsh’s t-test was chosen as opposed to a Student’s t-test because it was expected that individual effectors may vary in their interactions with other library members, thus not only altering the mean fold enrichment but also the standard deviation of fold enrichment scores. Once all the effectors were analyzed, p-values were corrected for a false discovery rate of 5%.

### Synergy scoring to identify synergistic and antagonistic combinations

To identify bivalent combinations with unexpected activities, synergy scores were calculated for each bivalent combination based on the expected effects of each individual effector (**Fig. 4a**). First, the individual effects of each effector were calculated using our marginal score metric, with the difference that each effector was scored for both the N-terminal and C-terminal position. Then, the sum of the corresponding marginal scores for each bivalent combination was computed. Because effectors could still be additive in nature even if the measured score did not exactly match the sum, a range for additive activity was defined that assumes that effectors may not contribute their full marginal strength to the combination but that the contribution of each effector should still be in the expected direction. For expected repressors, the lower bound of the additive range was defined as the minimum of the N-terminal marginal score, the C-terminal marginal score, and the sum of the marginal scores (min(N-term, C-term, Sum)), and the upper bound of the range was defined as the maximum of the sum of the marginal scores and the minimum of the two marginal scores (max(Sum, min(N-term, C-term))). For expected activators, the expressions are reversed and the additive range becomes (min(Sum, max(N-term, C-term))) to (max(N-term, C-term, Sum)). Log_2_ fold enrichment scores from HTS data were first mean centered, a logical step considering that marginal scores are relative contributions and are unaffected by central tendency, and subsequently compared to the additive range. In all cases, if the observed data fell within the additive range, the pair was considered additive and assigned a synergy score of 0. For expected repressors, bivalent combinations were considered “synergistic” if the measured score fell below the additive range, i.e. it was more repressive than expected, and the combination was considered “antagonistic” if the measured score fell above the additive range, i.e. it was less repressive than expected. Scores for expected activators were determined in the same way but in the opposite direction, with synergistic combinations exhibiting unexpectedly high observed scores and antagonistic combinations exhibiting unexpectedly low observed scores. In both cases, the synergy score was defined as how far outside the additive range the observed score fell, being positive for synergy and negative for antagonism. Because Library 1 was entirely composed of expected repressors, every combination was scored as a repressor. For Library 2, the sum of the marginal scores determined whether the combination was scored as an activator or a repressor with positive sums being scored as expected activators and negative sums being scored as expected repressors.

To test if individual effectors broadly synergize or antagonize with KRAB members from Library 1 or HDAC members from Library 2, the synergy scores from all combinations containing the given effector with a KRAB or HDAC member were tested against the null hypothesis that the average synergy score was zero using a one-sample Wilcoxon test. Only KRAB or HDAC members with marginal scores less than -0.5 were considered, and p-values were subsequently FDR corrected for multiple hypothesis testing. In addition to performing significance testing, the average synergy score and the average log_2_ fold enrichment score for the combinations were calculated.

### Comparison between stdMCP-KRAB and stdMCP-KRAB-L3MBTL3

pTRE3G-PuroR-T2A-stdMCP-XTEN80-KRAB(ZNF10)-L3MBTL3-WPRE and pTRE3G-PuroR-T2A-stdMCP-XTEN80-KRAB(ZNF10)-WPRE were infected by lentivirus to Tet-On dCas9 K562 (without CD81 guide array) in similar MOI (0.35 ± 0.04 for KRAB-L3MBTL3 and 0.31 ± 0.01 for KRAB). For comparing across different panels of genes and guides, an equal amount of lentivirus encoding both sgRNA and mCherry marker were infected to both cell lines. Doxycycline concentration was maintained at 1-2 µg/mL since the LV-sgRNA infection. 5 days after lentiviral infection of sgRNAs, cells were stained with corresponding antibodies (APC anti-human CD55 antibody (Cat# 311311, Biolegends), Alexa Fluor 647 Mouse Anti-Human CD58 antibody (Cat# 563567, BD Pharmingen), APC Mouse Anti-Human CD81 antibody (Cat# 561958, BD Bioscience), and APC anti-human CD151 antibody (Cat# 350405, Biolegends)) and expression level was measured by Attune NxT Flow Cytometer (Thermo Fisher Scientific). For the bidirectional perturbations using the MS2/PP7 system, pTREtight-stdPCP-p65-HSF1-T2A-BFP were lentivirally infected to the previous two cell lines described above. pTREtight-stdPCP-p65-HSF1-T2A-BFP was cloned from the construct pHR-CMV-stdPCP-p65-HSF1-T2A-GFP (Addgene #199456)^63^. The same amounts of lentivirus encoding both mouse U6 promoter-driven 2x MS2 sgRNA and human U6 promoter-driven 2x PP7 sgRNA were infected to both cell lines. Doxycycline concentration was maintained at 1-2 µg/mL since the day of LV-dual MS2/PP7 sgRNAs infection. CD81/CD274 expression profiles were measured 5 days post-infection of dual MS2/PP7 sgRNAs following staining with antibodies against both targets (APC Mouse Anti-Human CD81 antibody (Cat# 561958, BD Bioscience) and Alexa Fluor 488 Mouse Anti-Human CD274 antibody (Cat# 53-5983-42, Invitrogen)) by Attune NxT Flow Cytometer (Thermo Fisher Scientific).

## Data availability

All data to support the results are in the main text, figures and supplementary tables. Illumina and Nanopore sequencing datasets generated in this study will be available on the NCBI Sequence Read Archive, Bioproject PRJNA1143488.

## Code availability

The code for analyses performed in this paper will be available on GitHub.

## Acknowledgments

We thank the entire Hsu lab for helpful discussions and feedback, as well as B. Plosky and the Arc Institute Scientific Publications Team for assistance with the manuscript. P.D.H. is supported by funding from the Arc Institute, Yosemite, the Biswas Foundation, the Rainwater Foundation, the Curci Foundation, the Rose Hill Innovators Program, S. Altman, V. and N. Khosla, and by anonymous gifts to the Hsu Laboratory.

## Author Contributions

H.C.M., M.H.H., and P.D.H. conceived the study. H.C.M., M.H.H., S.K., and P.D.H. designed experiments. H.C.M. and M.H.H. performed experiments and computational analyses. P.D.H. and S.K. supervised the research. H.C.M., M.H.H., A.P., S.K., and P.D.H. wrote the manuscript.

