## Supplementary Information for "A combinatorial domain screening platform reveals epigenetic effector interactions for transcriptional perturbation"

### Table of contents

1. **Supplementary Figures**
    1. Classifications and size distributions of individual domains from Library 1 and Library 2
    2. Library 1 combinatorial cloning strategy
    3. Library 2 combinatorial cloning and barcode mapping strategy
    4. Doxycycline-inducible dCas9/MS2 system enables temporal control of transcriptional perturbation
    5. Detailed procedures for Library 1 HTS
    6. Detailed procedures for Library 2 HTS and barcode mapping
    7. Investigation of lentiviral barcode swapping by targeted nanopore sequencing
    8. Number of acquired NGS reads versus bivalent effector length
    9. Heatmaps of enrichment scores from later HTS timepoints
    10. N- versus C-terminal position effects
    11. Synergy score heatmaps highlight synergistic and antagonistic interactions in Library 1
    12. Synergy score heatmaps highlight synergistic and antagonistic interactions in Library 2
    13. Synergistic interaction between the H2AK119ub1 reader and a subset of PRC1 recruiters
    14. HDAC and KRAB members included in the average synergy score calculation
    15. Epigenetic modifications at CD81 locus
    16. Key effectors from Library 2 transiently perturb CD55, CD58, CD151, and CD155
    17. Gene-by-gene scatter plots of percent change in MFI induced by key effectors from Library 2
    18. Synergistic partners of KRAB
    19. Quantification of bidirectional perturbations by MS2/PP7 system
    20. Marginal effector and synergy score analysis on Day 12
    21. DNMT3A-3L combinations induce long-term partial silencing of CD81
    22. UBE2E1 + DNMT3A induces long-term repression of CD81
    23. Violin plot timecourses of TET1 combinations
    24. TET1 combinations induce long-term activation of CD81
2. **Supplementary Note**

1. Synergistic interaction between the H2AK119ub1 reader and a subset of PRC1 recruiters

#### 3. Supplementary Tables

1. Library 1 members and sequences
2. Library 2 members and sequences
3. Library 1 HTS results
4. Library 2 HTS results
5. Catalytic mutations
6. Library 1 synergy scores
7. Library 2 synergy scores
8. Plasmid, primer, and guide sequences
9. Library 1 individual validation details

### Supplementary Figures

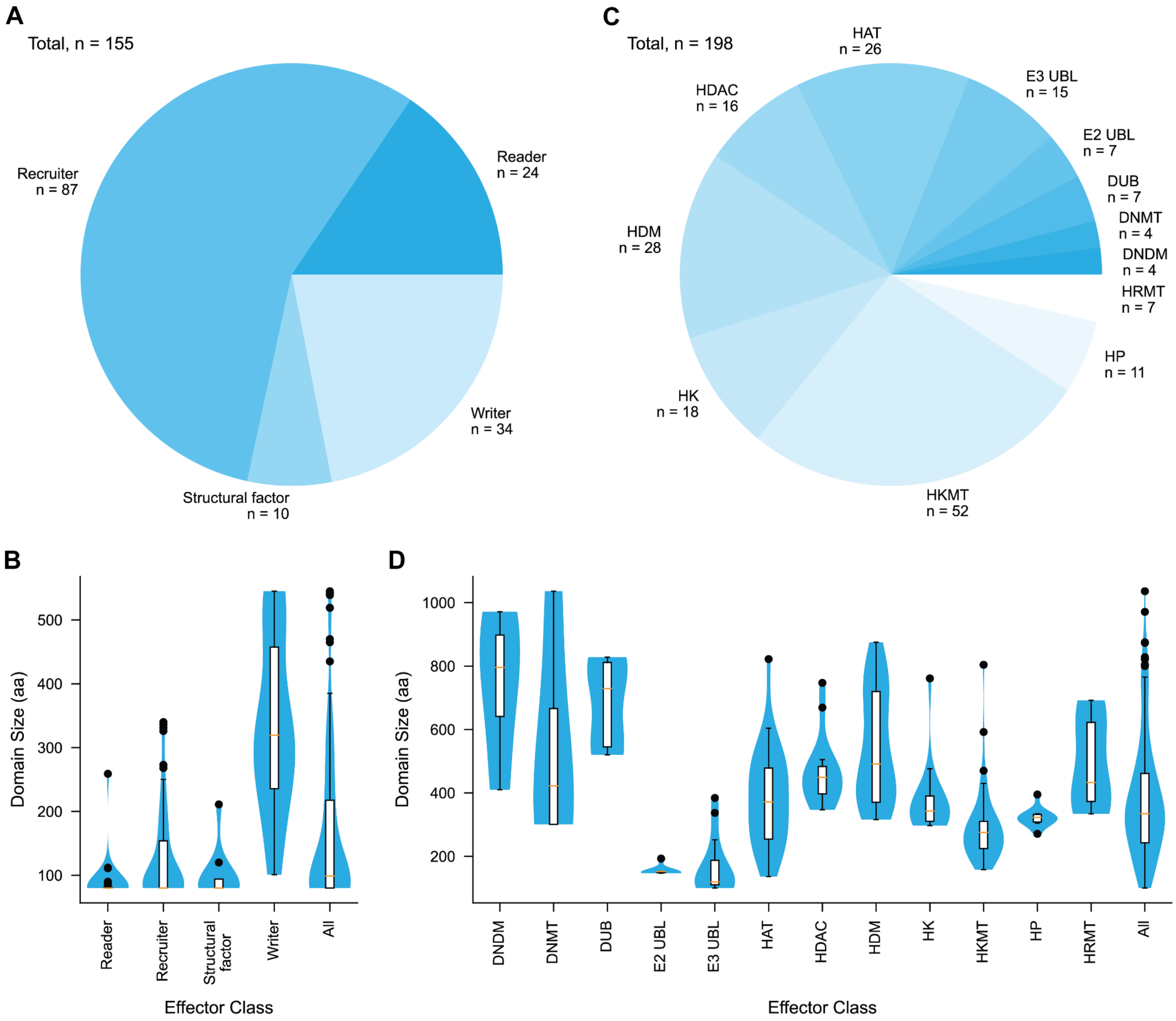


**Supplementary Figure 1. Classifications and size distributions of individual domains from Library 1 and Library 2**

(**A**) Classification of Library 1 members.

(**B**) Size distributions of specific classes from Library 1.

(**C**) Classification of Library 2 members. DNA demethylation machinery (DNDM), DNA methyltransferase (DNMT), E2 and E3 ubiquitin ligases (UBL), histone acetyltransferase (HAT), histone arginine methyltransferase (HRMT), histone deacetylase (HDAC), histone demethylase (HDM), histone deubiquitinase (DUB), histone kinase (HK), histone lysine methyltransferase (HKMT), and histone phosphatase (HP).

(**D**) Size distributions of specific classes from Library 2.

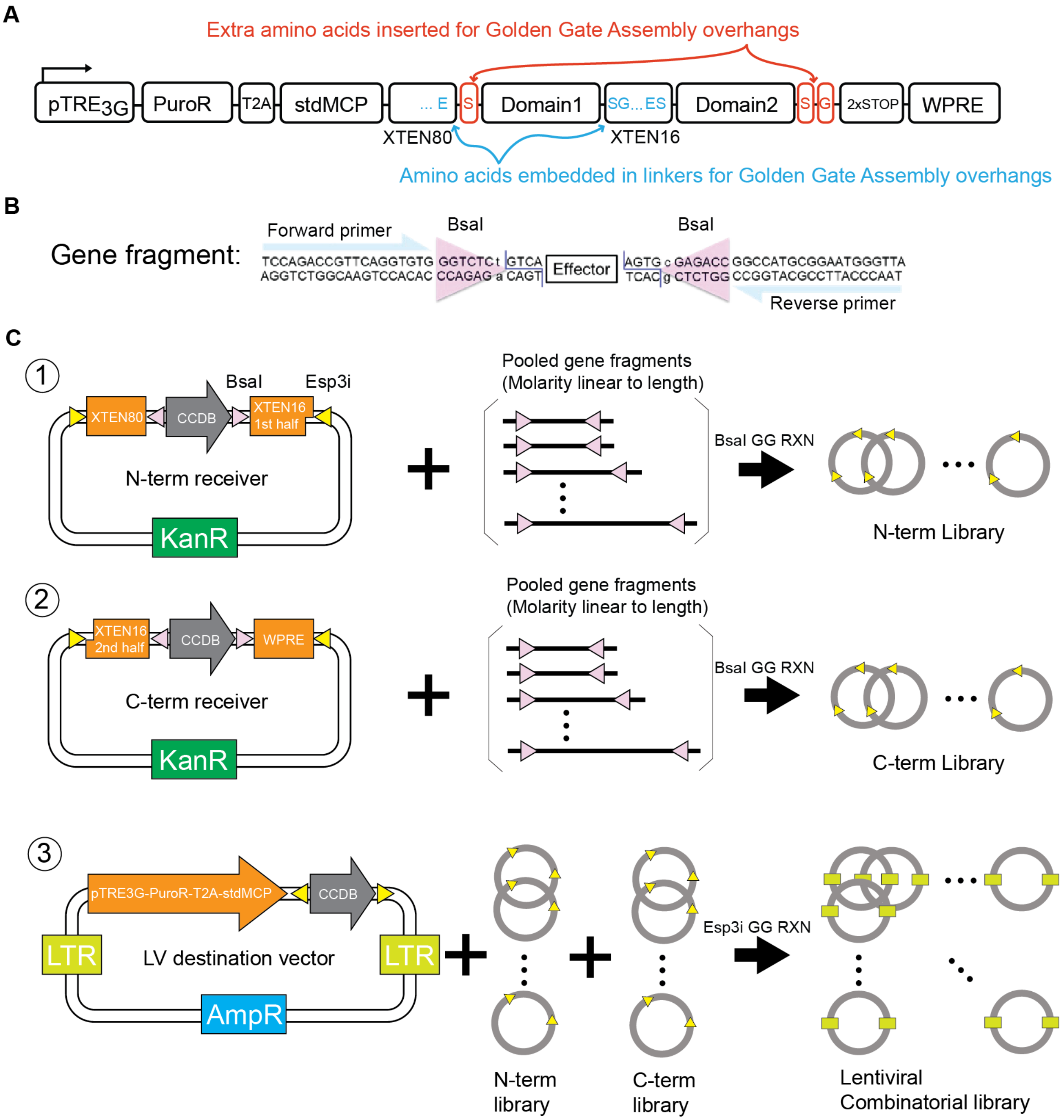


**Supplementary Figure 2. Library 1 combinatorial cloning strategy**

(**A**) Schematic of combinatorial effector candidate cassette in lentiviral vector for HTS of Library 1. DNA sequence encoding for Glutamic acid (E) – Serine (S) serves as a 5` overhang for the golden gate cloning. DNA sequence encoding for Serine (S) – Glycine (G) serves as a 3` overhang for the golden gate cloning. Cyan amino acid symbols indicate amino acids already encoded in the XTEN linker CDS. Orange amino acid symbols indicate amino acids inserted for the golden gate cloning overhang.

(**B**) Schematic of gene fragments ordered for Library 1 (**Supplementary Table 1**). Although we included PCR amplification handles in the design, it was not necessary to PCR amplify the gene fragments for cloning.

(**C**) Library cloning procedure for the generation of Library 1 bivalent domain candidates. The procedure consists of three sequential pooled Golden Gate assemblies. The first Golden Gate reaction involves the library cloning of pooled gene fragments into the N-term/KanR1 receiver (pCM001). Each gene fragment was pooled with a molar ratio linear to its size including the PCR amplification handles. For example, we added 1.5-fold of the 450 bp gene fragment compared to the 300 bp gene fragment. The second Golden Gate reaction involves the library cloning of pooled gene fragments into the C-term/KanR2 receiver (pCM002). The final Golden Gate reaction involves the library cloning of the N-terminus library and C-terminus library into the LV_AmpR_backbone (pCM003).

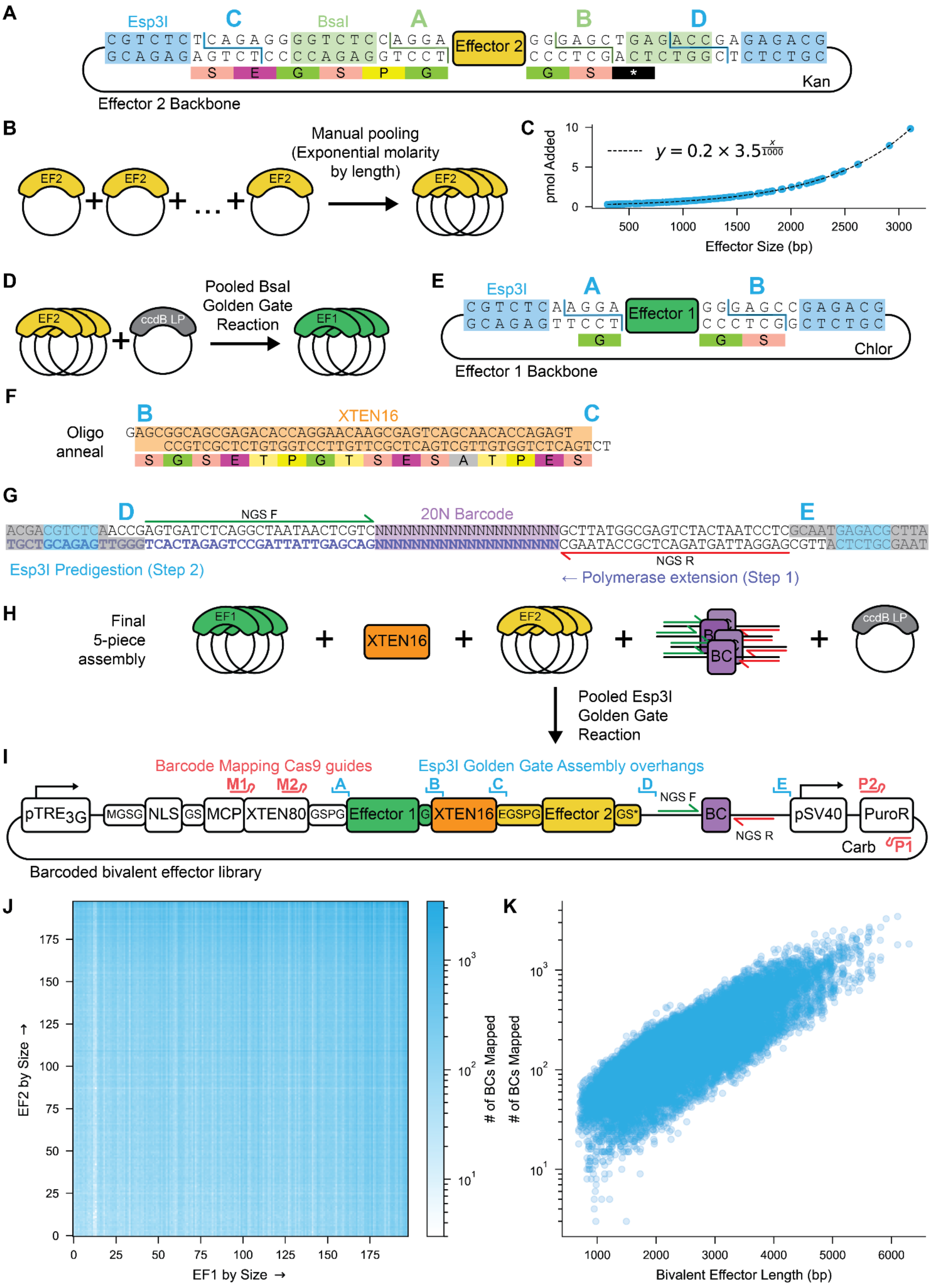

A
Esp3I
C
BsaI
A
B
D
CGTCTCTCAGAGGGGTCTCCAGGA
GCAGAGAGTCTCCCCAGAGGTCCT
Effector 2
GGGAGCTGAGACCGAGAGACG
CCCTCGACTCTGGCTCTCTGC
S E G S P G
G S *
Kan
Effector 2 Backbone
B
EF2
Manual pooling
(Exponential molarity by length)
C
pmol Added
$y = 0.2 \times 3.5^{\frac{x}{1000}}$
Effector Size (bp)
D
EF2
ccdB LP
Pooled BsaI Golden Gate Reaction
EF1
E
Esp3I
A
B
CGTCTCAAGGA
GCAGAGTTCCT
Effector 1
GGGAGCCGAGACG
CCCTCGGCTCTGC
G
G S
Chlor
Effector 1 Backbone
F
B
XTEN16
C
Oligo anneal
GAGCGGCAGCGAGACACCAGGAACAAGCGAGTCAGCAACACCAGAGT
CCGTCGCTCTGTGGTCCTTGTTCGCTCAGTCGTTGTGGTCTCAGTCT
S G S E T P G T S E S A T P E S
G
D
NGS F
20N Barcode
E
ACGACGTCTCAACCGAGTGATCTCAGGCTAATAACTCGTCNNNNNNNNNNNNNNNNNNNNGCTTATGGCGAGTCTACTAATCCTCGCAATGAGACGCTTA
TGCTGCAGAGTTGGGTCACTAGAGTCCGATTATTGAGCAGNNNNNNNNNNNNNNNNNNNNCGAATACCGCTCAGATGATTAGGAGCGTTACTCTGCGAAT
NGS R
Esp3I Predigestion (Step 2)
← Polymerase extension (Step 1)
H
Final 5-piece assembly
EF1
XTEN16
EF2
BC
ccdB LP
Pooled Esp3I Golden Gate Reaction
I
Barcode Mapping Cas9 guides
Esp3I Golden Gate Assembly overhangs
M1
M2
A
B
C
D
E
P2
NGS F
pTRE3G
MGSG
NLS
GS
MCP
XTEN80
GSPG
Effector 1
G
XTEN16
EGSPG
Effector 2
GS*
BC
NGS R
pSV40
PuroR
Carb
P1
Barcoded bivalent effector library
J
EF2 by Size →
EF1 by Size →
### of BCs Mapped
K
### of BCs Mapped
Bivalent Effector Length (bp)

**Supplementary Figure 3. Library 2 combinatorial cloning and barcode mapping strategy**

(**A**) Schematic of clonal gene fragments ordered for Library 2 with dual golden gate overhangs.
(**B**) Clonal fragments of individual effectors were pooled manually to form the Effector 2 pool.
(**C**) The amount of plasmid DNA added per effector increased 3.5-fold per kb.
(**D**) The first golden gate assembly reaction for cloning effectors from Effector 2 pool into Effector 1 landing pad plasmid.
(**E**) Schematic of cloned Effector 1 plasmid with golden gate overhangs.
(**F**) XTEN16 linker was ordered as ssDNA and annealed with proper golden gate overhangs.
(**G**) Barcodes were ordered as a ssDNA fragment containing a 20N sequence. Polymerase extension was used to fill in barcodes, forming a dsDNA fragment, before the fragment was predigested with Esp3I to form proper golden gate overhangs.
(**H**) Final 5-piece golden gate reaction for generating Library 2 bivalent effectors for HTS.
(**I**) Schematic of Library 2 bivalent effectors cloned into a lentiviral vector for HTS. Golden gate assembly overhang positions are indicated in blue. Barcode mapping Cas9 guides are indicated in red.
(**J**) Heatmap of barcodes mapped per bivalent effector, with effectors organized by increasing size.
(**K**) Scatterplot of bivalent effector length versus number of mapped barcodes.

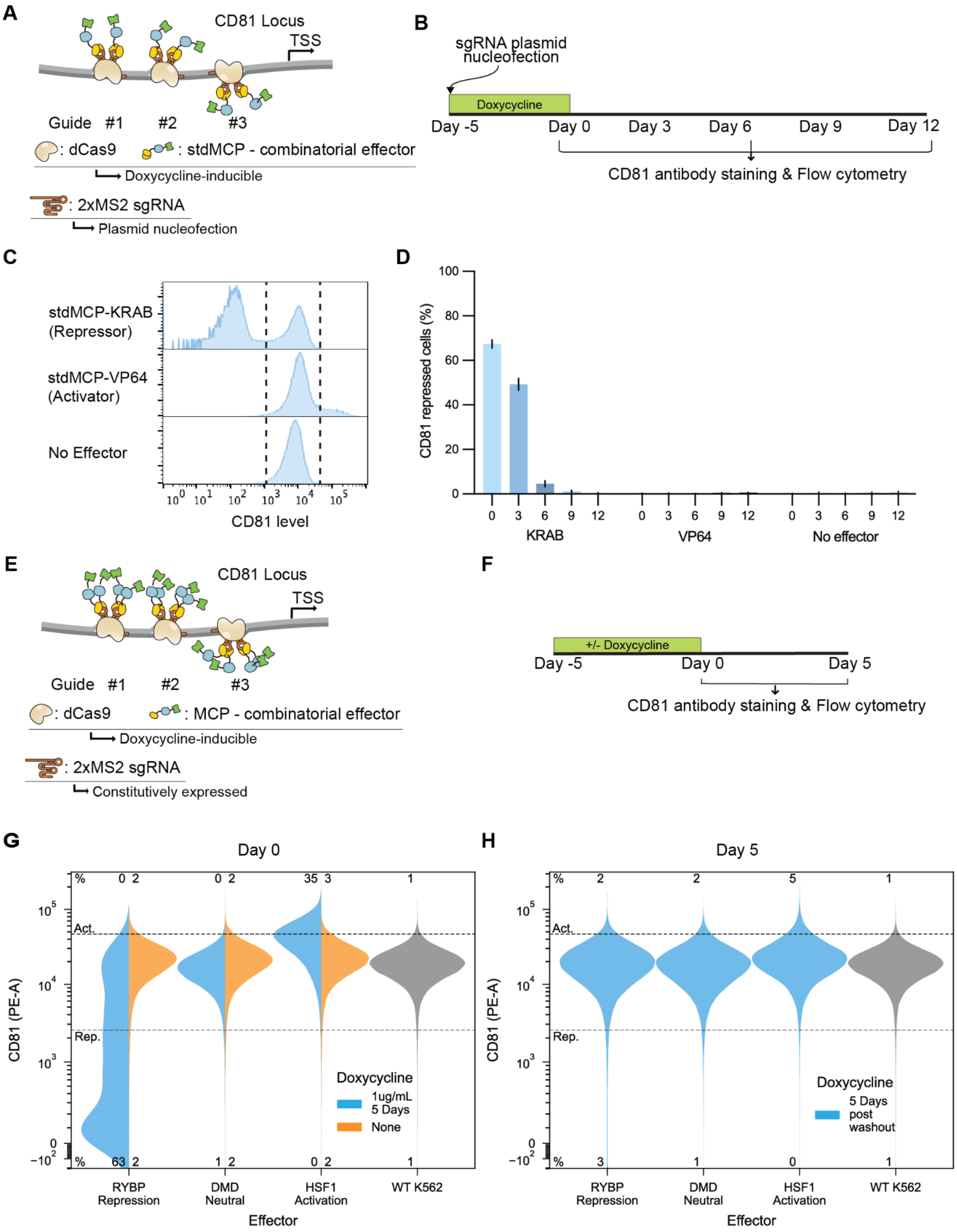
A
CD81 Locus
TSS
Guide #1 #2 #3
: dCas9
: stdMCP - combinatorial effector
Doxycycline-inducible
: 2xMS2 sgRNA
Plasmid nucleofection
B
sgRNA plasmid nucleofection
Doxycycline
Day -5
Day 0
Day 3
Day 6
Day 9
Day 12
CD81 antibody staining & Flow cytometry
C
stdMCP-KRAB (Repressor)
stdMCP-VP64 (Activator)
No Effector
CD81 level
D
CD81 repressed cells (%)
KRAB
VP64
No effector
E
CD81 Locus
TSS
Guide #1 #2 #3
: dCas9
: MCP - combinatorial effector
Doxycycline-inducible
: 2xMS2 sgRNA
Constitutively expressed
F
+/- Doxycycline
Day -5
Day 0
Day 5
CD81 antibody staining & Flow cytometry
G
Day 0
CD81 (PE-A)
Act.
Rep.
Doxycycline
1ug/mL 5 Days
None
RYBP Repression
DMD Neutral
HSF1 Activation
WT K562
Effector
H
Day 5
CD81 (PE-A)
Act.
Rep.
Doxycycline
5 Days post washout
RYBP Repression
DMD Neutral
HSF1 Activation
WT K562
Effector

**Supplementary Figure 4. Doxycycline-inducible dCas9/MS2 system enables temporal control of transcriptional perturbation**

(**A**) Illustration of dCas9/MS2 system for Library 1 HTS. Tet-On dCas9 K562 (without CD81 guide array) cell line was used for the experiment. Combinatorial effectors are fused to stdMCP and delivered by lentiviral infection. Both dCas9 and combinatorial effectors are inducible by doxycycline. 3X CD81 guide array is introduced by plasmid nucleofection.

(**B**) Timeline for testing dCas9/MS2 system used for Library 1 with established epigenetic repressor KRAB from ZNF10 protein, activator VP64, and no effector control.

(**C**) Representative histograms of CD81 expression at 5 days post-nucleofection of CD81 targeting 3X guide array plasmids.

(**D**) Timecourse of CD81 repressed cells after nucleofection of CD81 targeting 3X guide array plasmids. Error bars indicate the standard deviation of 3 biological replicates.

(**E**) Illustration of dCas9/MS2 system for Library 2 HTS. Tet-On dCas9 K562 (with CD81 guide array) cell line was used for the experiment. Combinatorial effectors were fused to MCP and introduced by lentivirus. CD81 targeting sgRNAs were constitutively expressed. Both dCas9 and combinatorial effectors are inducible by doxycycline.

(**F**) Timeline for testing dCas9/MS2 system used for Library 2 with established epigenetic repressor RYBP, activator HSF1, and neutral control 400 aa fragment of DMD.

(**G**) Violin plots of CD81 expression at experimental Day 0 with or without the addition of doxycycline.

(**H**) Violin plots of CD81 expression 5 days after washout of doxycycline after initial 5 days of doxycycline treatment.

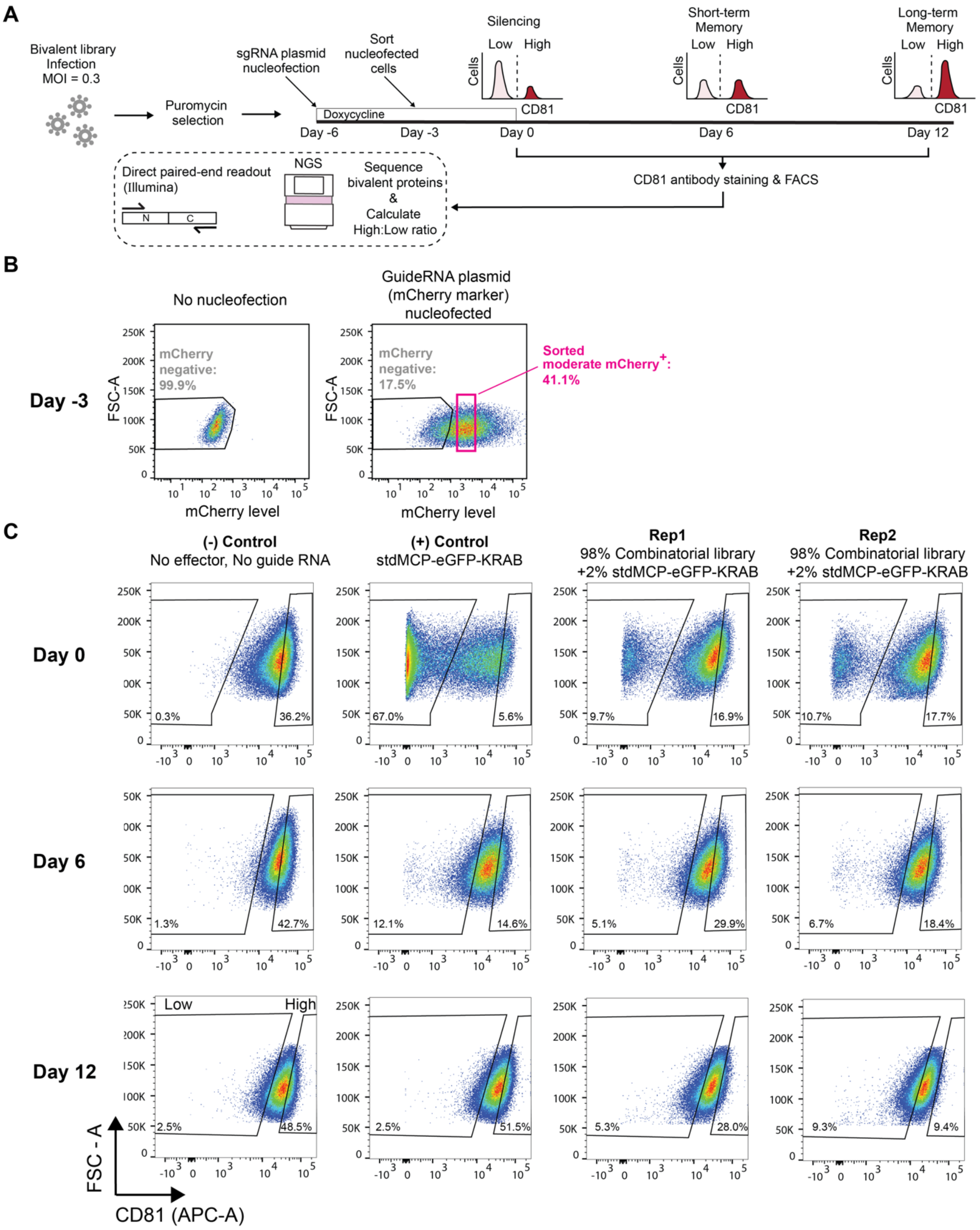


**D**

| | Sorting criteria | Rep 1 | Rep 2 |
|---|---|---|---|
| Day 0 | CD81 level High ~10 % | ~ 0.75M | ~ 1M |
| | CD81 level Low ~10 % | ~ 0.75M | ~ 1M |
| Day 6 | CD81 level High ~8 % | ~ 2.6M | ~ 2.43M |
| | CD81 level High ~9 % | ~ 2.84M | ~ 2.63M |
| Day 12 | CD81 level High ~10 % | ~ 3.33M | ~ 3.1M |
| | CD81 level Low ~10 % | ~ 3.39M | ~ 3.4M |

**Supplementary Figure 5. Detailed procedures for Library 1 HTS**

(**A**) Timeline and NGS strategy for Library 1 HTS. CD81 targeting guide array plasmid was nucleofected, and nucleofected cells were sorted based on the mCherry marker present in the plasmid (**Supplementary Table 8**, Plasmids, pMH222).

(**B**) Representative FACS plots of mCherry level at 3 days post-nucleofection of 3X guide array plasmids which express mCherry as a marker. Cells that were successfully nucleofected with 3X guide array plasmids were sorted based on mCherry level. To reduce heterogeneity in the sgRNA level, moderate mCherry-expressing cells were sorted.

(**C**) FACS plots of multiple conditions from Library 1 HTS. Approximate sorting gates are drawn. The actual gating was continuously amended during sorting to match the aimed High/Low percentage of the sorted cells. This procedure was necessary as the staining (CD81, APC) histogram shifted as the sorting procedure prolonged.

(**D**) Number of cells sorted per screening condition.

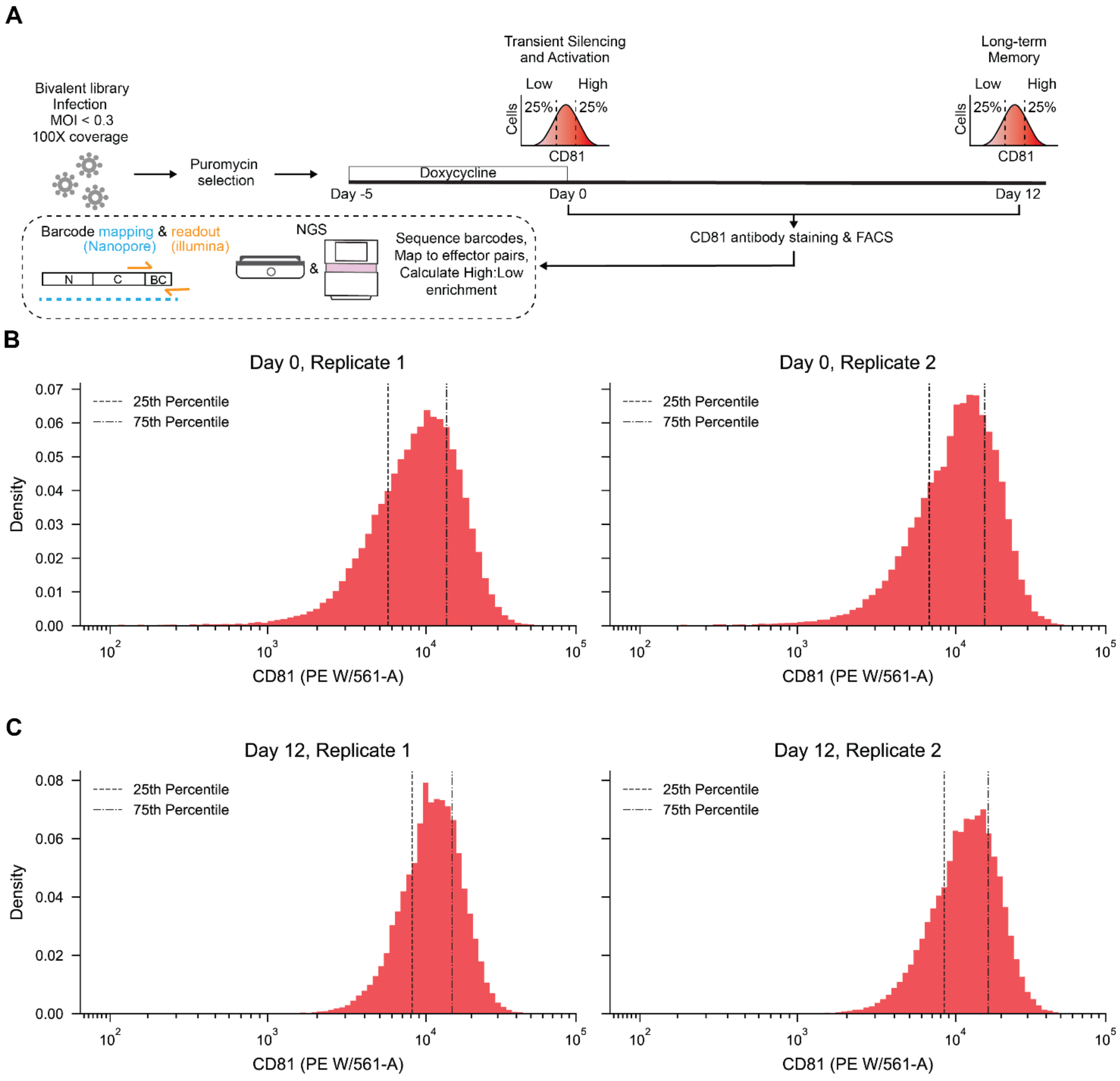


D

| Day | Replicate | Low | High |
|---|---|---|---|
| 0 | 1 | 12250852 | 10189885 |
| 0 | 2 | 8987437 | 7577374 |
| 12 | 1 | 8857130 | 7440798 |
| 12 | 2 | 11226879 | 9447047 |

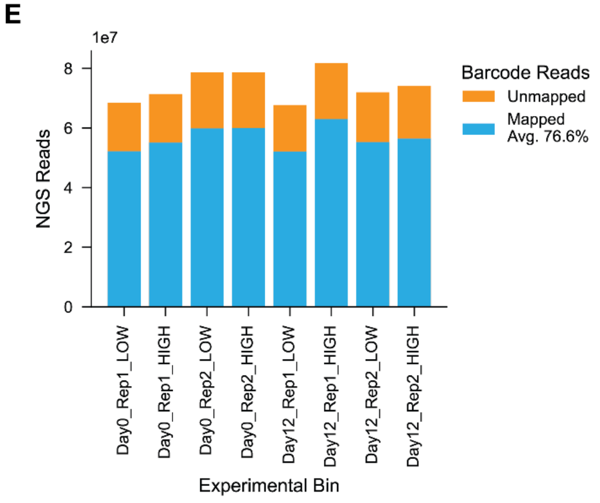

**Supplementary Figure 6. Detailed procedures for Library 2 HTS and barcode mapping**

(**A**) Timeline and NGS strategy for Library 2 HTS. CD81 targeting sgRNAs were constitutively expressed from Tet-On dCas9 K562 cell lines (with CD81 guide array).
(**B**) FACS plots of Library 2 HTS on Day 0 illustrating 25th and 75th percentile gates, which were continuously adjusted throughout the sort.
(**C**) FACS plots of Library 2 HTS on Day 12 illustrating 25th and 75th percentile gates, which were continuously adjusted throughout the sort.
(**D**) Number of cells sorted per screening condition.
(**E**) Number of NGS reads per screening condition with the fraction of reads mapped to bivalent effectors indicated in blue.

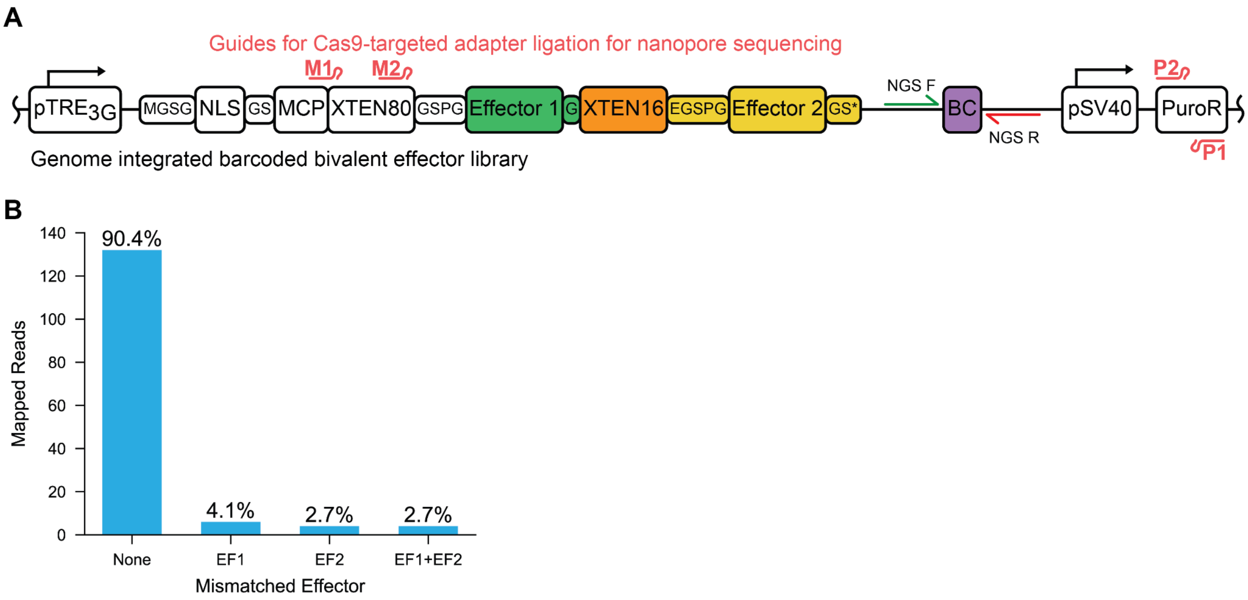


**Supplementary Figure 7. Investigation of lentiviral barcode swapping by targeted nanopore sequencing**

(**A**) Schematic of genome integrated bivalent effectors from Library 2. Cas9 guide positions for targeted nanopore sequencing are shown in red.

(**B**) Number and percentage of genome-integrated nanopore reads with barcodes matching or mismatching effector pairs mapped by nanopore plasmid sequencing.

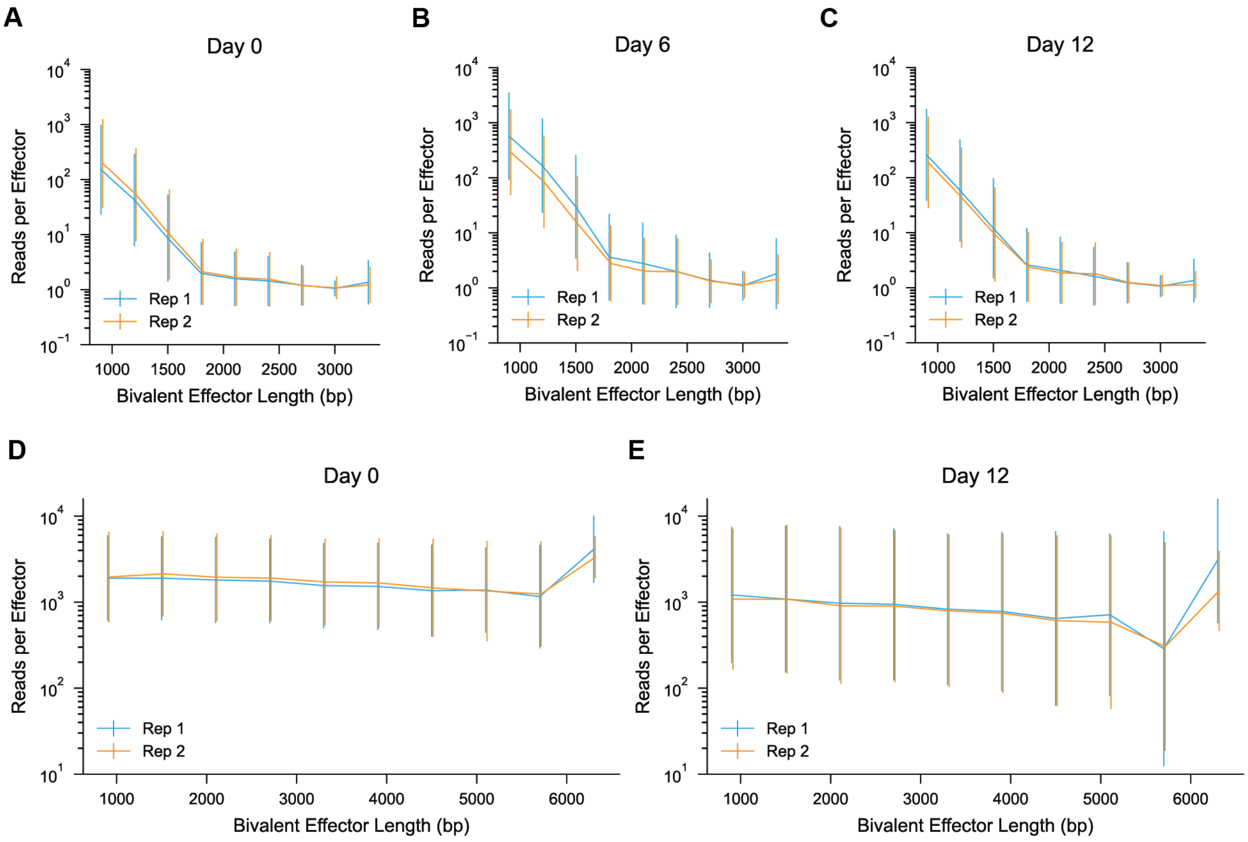


**Supplementary Figure 8. Number of acquired NGS reads versus bivalent effector length**

(**A**) Number of NGS reads versus length of bivalent effector from Library 1 HTS Day 0 data.
(**B**) Number of NGS reads versus length of bivalent effector from Library 1 HTS Day 6 data.
(**C**) Number of NGS reads versus length of bivalent effector from Library 1 HTS Day 12 data.
(**D**) Number of NGS barcode reads versus length of the corresponding bivalent effector from Library 2 HTS Day 0 data.
(**E**) Number of NGS barcode reads versus length of the corresponding bivalent effector from Library 2 HTS Day 12 data.

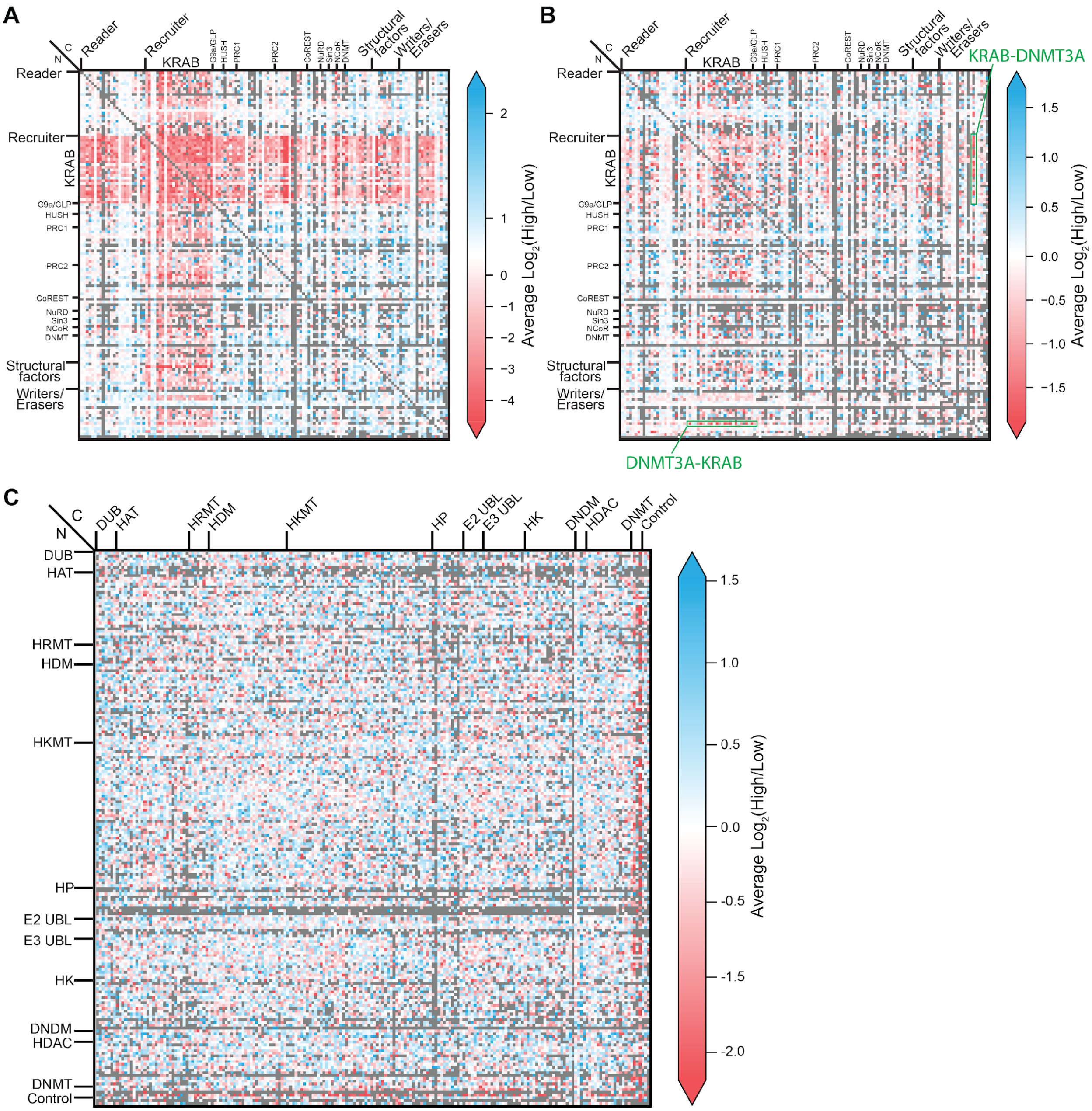


**Supplementary Figure 9. Heatmaps of enrichment scores from later HTS timepoints**

(**A**) Enrichment score heatmap from Library 1 HTS on Day 6.

(**B**) Enrichment score heatmap from Library 1 HTS on Day 12. KRAB-DNMT3A and DNMT3A-KRAB combinations are marked with green boxes (**Supplementary Table 3**).

(**C**) Enrichment score heatmap from Library 2 HTS on Day 12.

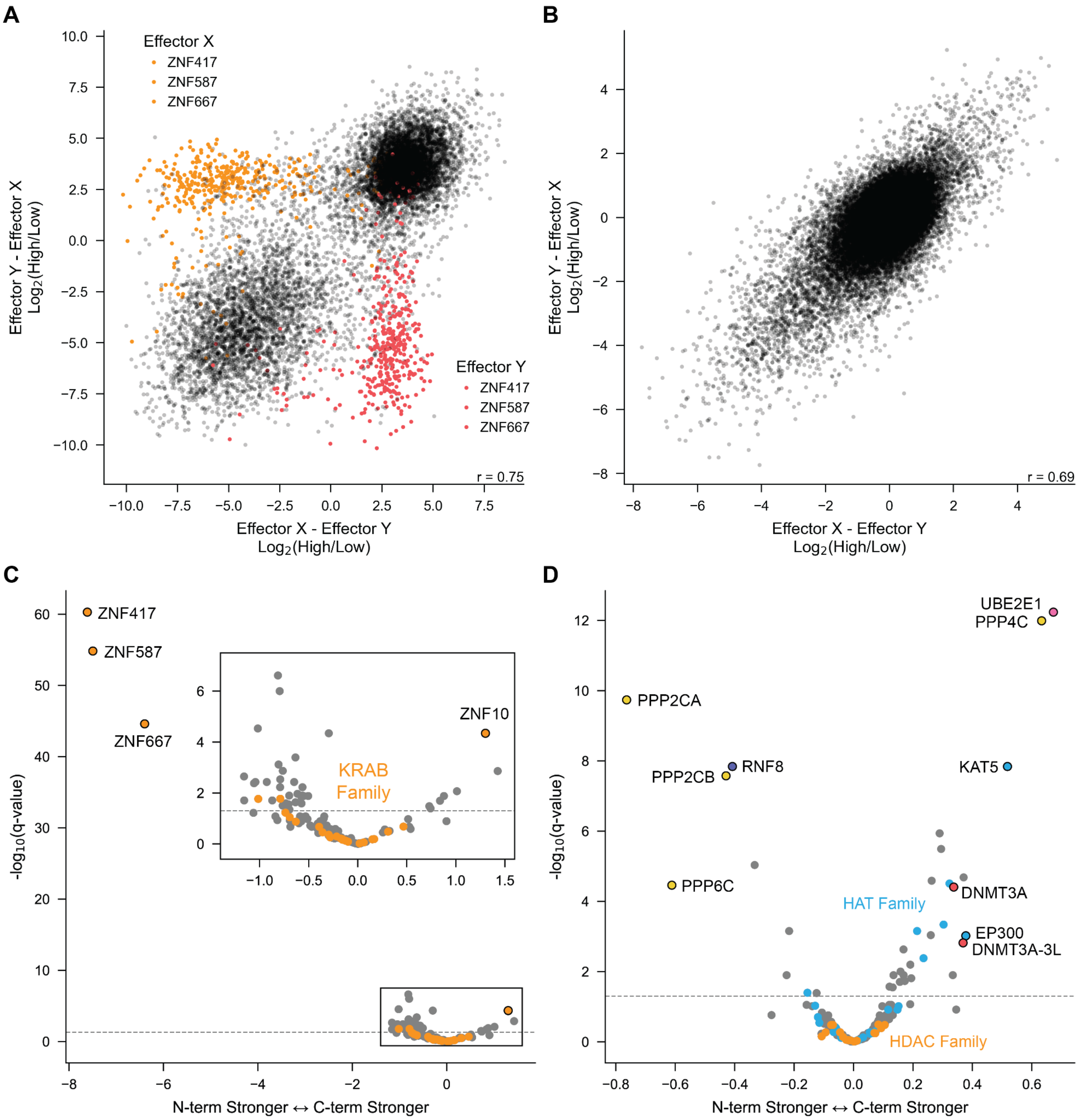


**Supplementary Figure 10. N- versus C-terminal position effects**

(**A**) Scatter plot of enrichment scores from X-Y versus Y-X bivalent effector pairs from Library 1 HTS on Day 0. Effectors with strong positional preference for the N-terminus are indicated in the X position in orange and the Y position in red.

(**B**) Scatter plot of enrichment scores from X-Y versus Y-X bivalent effector pairs from Library 2 HTS on Day 0.

(**C**) Volcano plot illustrating positional preference of individual effectors from Library 1. X-axis score calculated as the difference in mean $\log_2$(High/Low) between matched effector pairs with given effector at N- versus C-terminus (C-terminus minus N-terminus), multiplied by -1 for effectors with repressive marginal scores and +1 for effectors with activating marginal scores. Q-values were calculated as FDR-corrected p-values from a two-sample t-test. Only effectors with ≥30 matched pairs are shown.

(**D**) Volcano plot illustrating positional preference of individual effectors from Library 2. Calculated as (**C**).

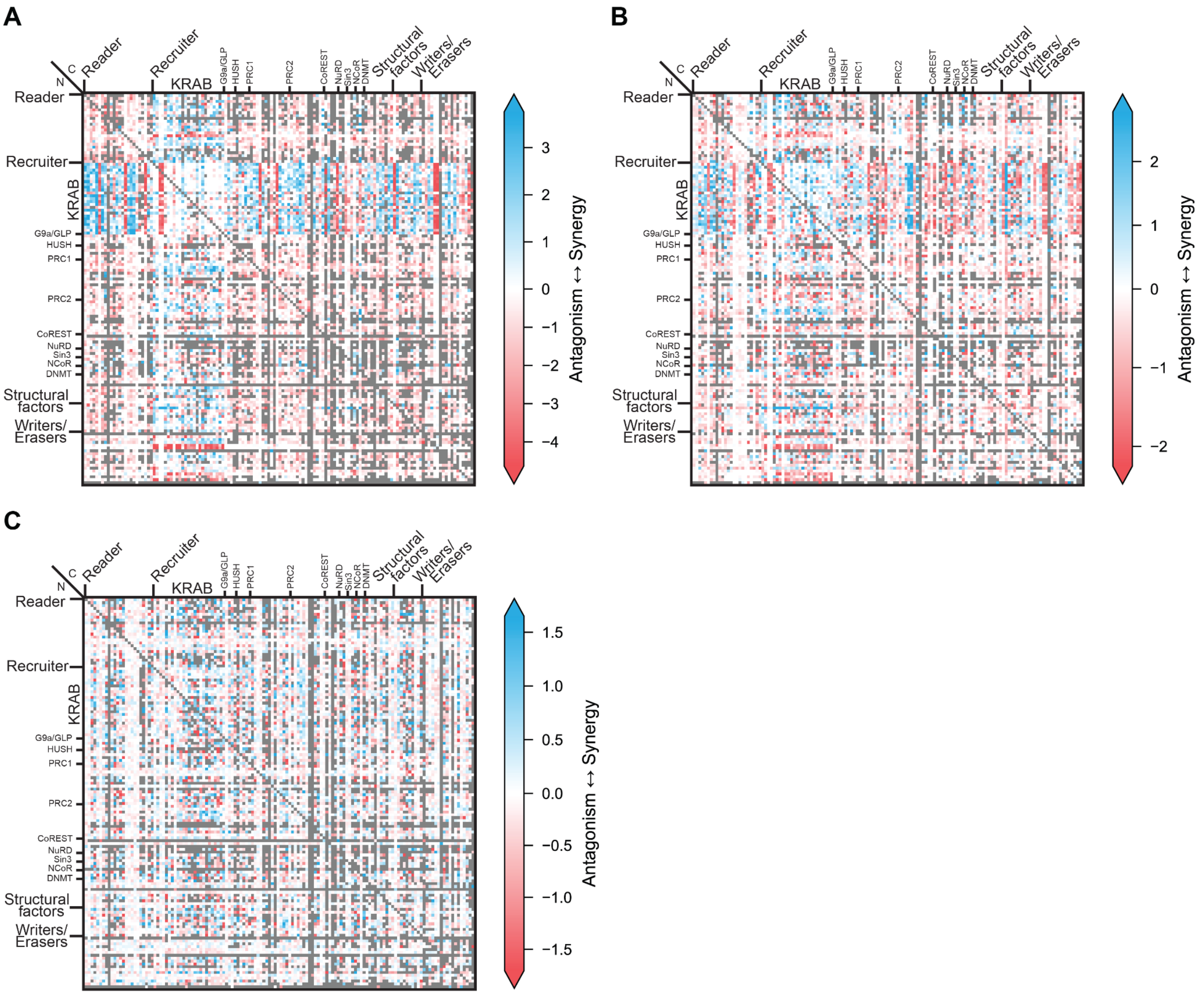


**Supplementary Figure 11. Synergy score heatmaps highlight synergistic and antagonistic interactions in Library 1**

Heatmaps illustrating synergy scores for bivalent combinations in Library 1 on Day 0 (**A**), Day 6 (**B**), and Day 12 (**C**). Red indicates antagonism where measured scores were less repressive than expected, and blue indicates synergy where scores were more repressive than expected. The magnitude of the score represents how far measured scores were outside of the expected (additive) range.

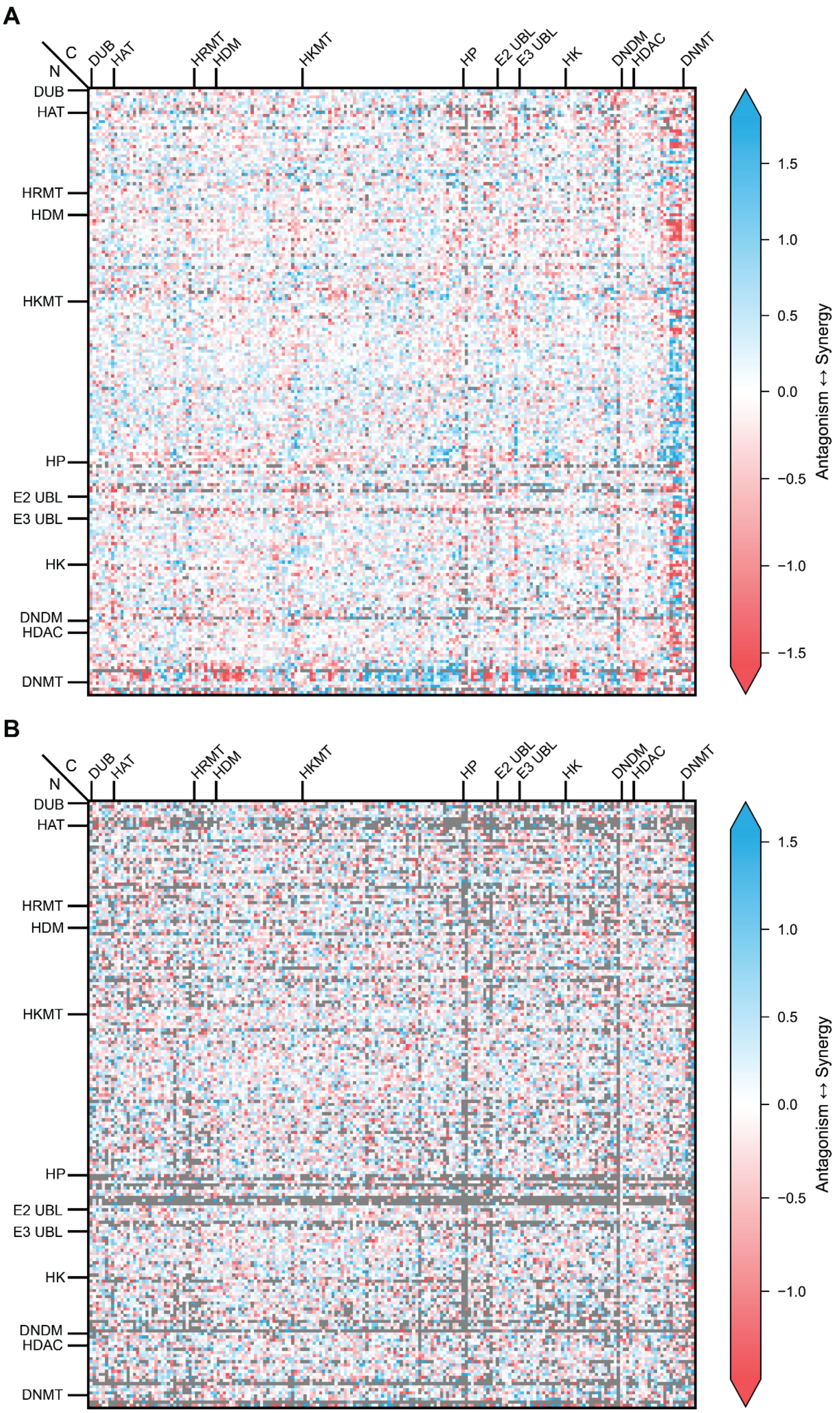


**Supplementary Figure 12. Synergy score heatmaps highlight synergistic and antagonistic interactions in Library 2**

Heatmaps illustrating synergy scores for bivalent combinations in Library 2 on Day 0 (**A**) and Day 12 (**B**). Red indicates antagonism where measured scores were less repressive or activating than expected, and blue indicates synergy where scores were more repressive or activating than expected. Scores were computed in the direction of the stronger effector. The magnitude of the score represents how far measured scores were outside of the expected (additive) range.

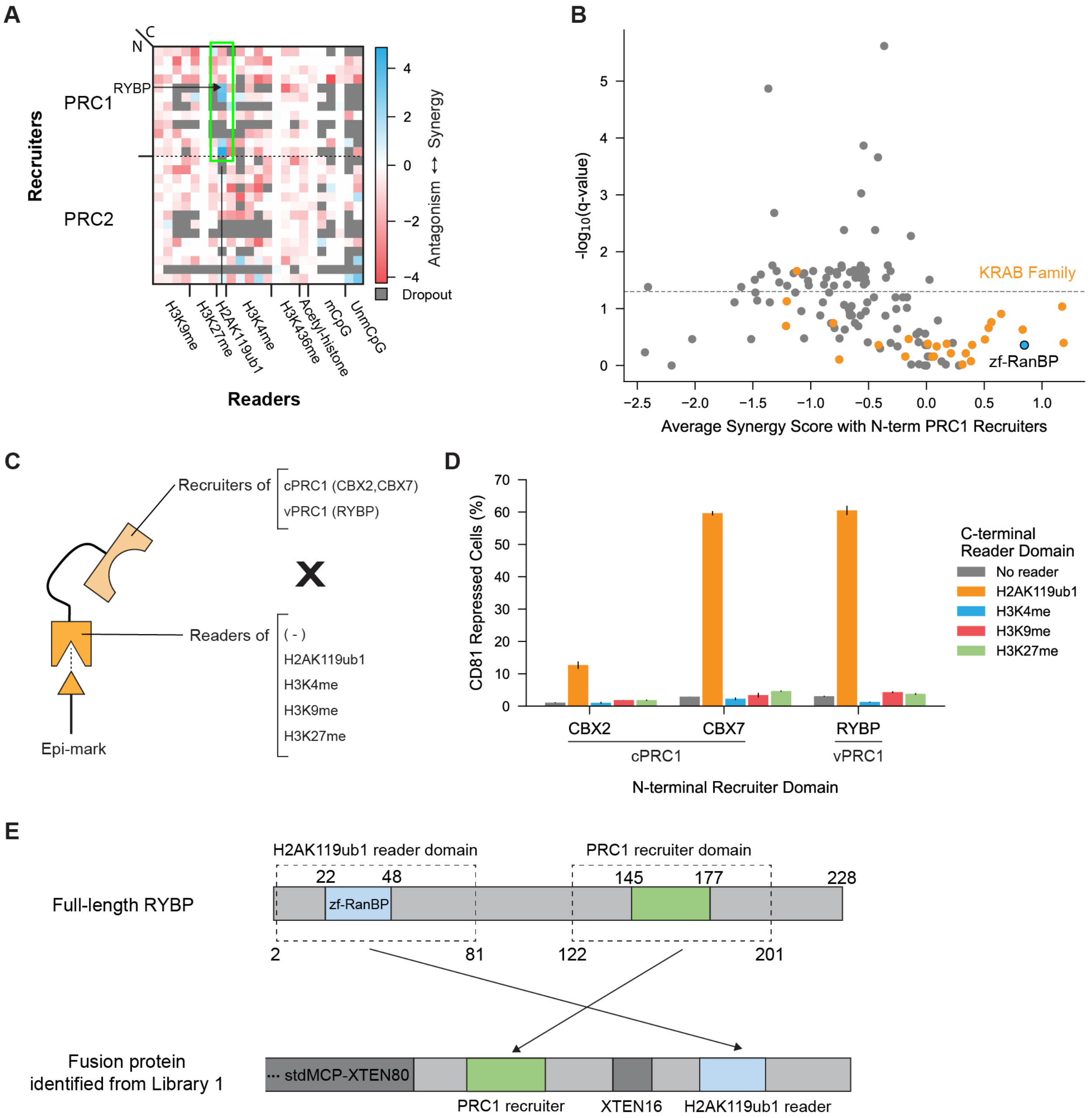


#### Supplementary Figure 13. Synergistic interaction between the H2AK119ub1 reader and a subset of PRC1 recruiters

(**A**) Sub-heatmap of synergy scores from Library 1 HTS Day 0 data with recruiters of PRC1&2 on the N-terminus and readers of epigenetic modifications on the C-terminus. The green box indicates combinations between PRC1 recruiters and the H2AK119ub1 reader domain from the RYBP protein. The PRC1 recruiting domain from the RYBP protein is indicated by a horizontal arrow.

(**B**) Volcano plot of average synergy scores with PRC1 recruiters on the N-terminus from Library 1 HTS Day 0 data. Combinations with the KRAB family on the C-terminus are colored orange. Combination with the H2AK119ub1 mark reader, the zf-RanBP domain from RYBP protein, on

the C-terminus is colored blue. Q-values were calculated as FDR-corrected p-values from a one-sample Wilcoxon test. Horizontal dashed line indicates q-value of 0.05.
(**C**) Illustration of PRC1 recruiter + H2AK119ub1 reader fusion proteins tested for validation.
(**D**) Percentage of CD81 repressed cells at 4 days post-nucleofection of fusion protein plasmids. CBX2 and CBX7 are components of canonical PRC1 (cPRC1) whereas RYBP is a component of variant PRC1 (vPRC1). Error bars indicate the standard deviation of 2 biological replicates.
(**E**) Illustration of full-length RYBP and potent fusion protein identified from Library1 HTS on Day 0.

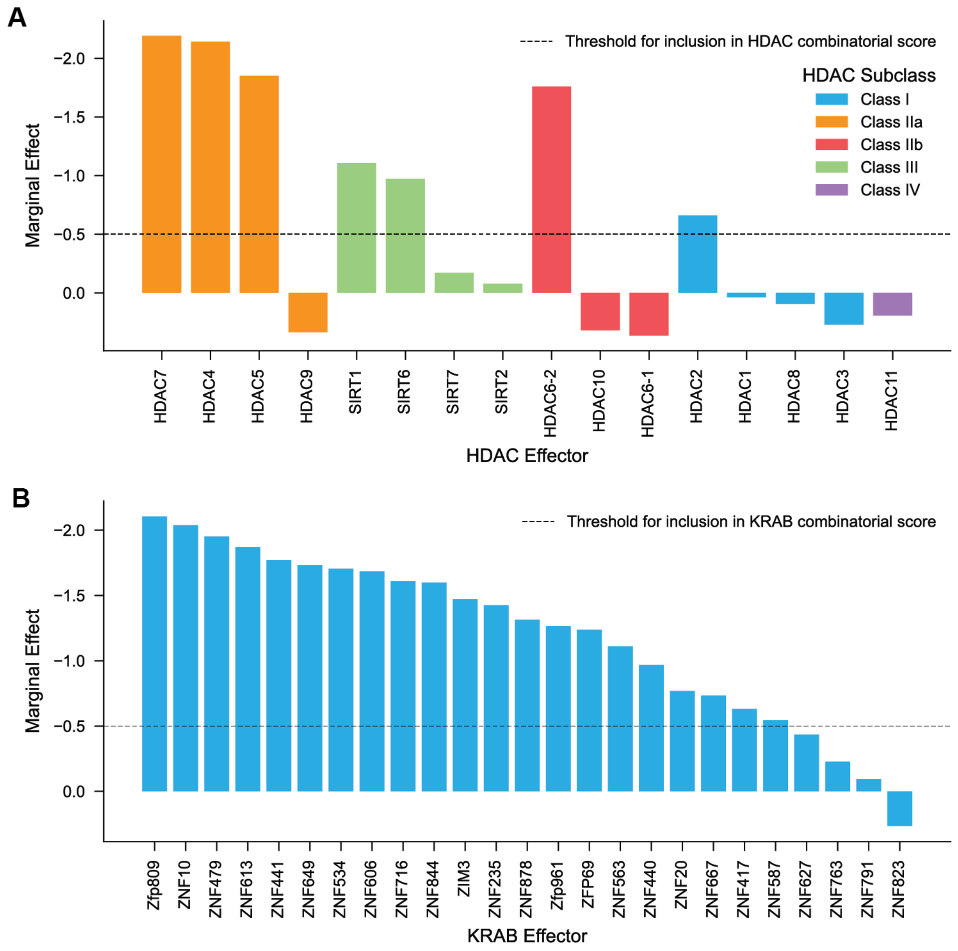


**Supplementary Figure 14. HDAC and KRAB members included in the average synergy score calculation**

(**A**) Marginal scores of HDAC members from Library 2 stratified by HDAC subclass. The dashed line indicates the threshold for inclusion in the average synergy score calculation. Only members with marginal scores less than -0.5 were included.

(**B**) Marginal scores of KRAB members from Library 1. The dashed line indicates the threshold for inclusion in the average synergy score calculation. Only members with marginal scores less than -0.5 were included.

**A**

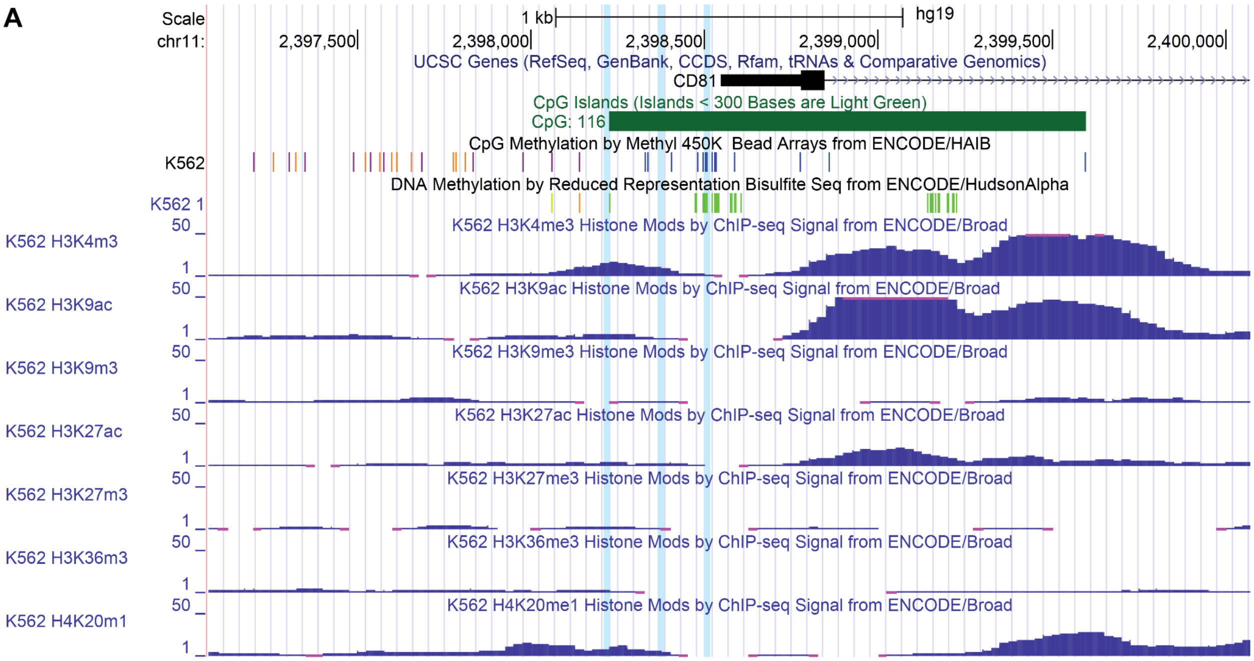


**Supplementary Figure 15. Epigenetic modifications at CD81 locus**

(**A**) Visualization of endogenous CD81 locus with mapped epigenetic modifications. Vertical sky blue bars indicate positions of sgRNA binding. Plot generated with UCSC Genome Browser at http://genome.ucsc.edu.

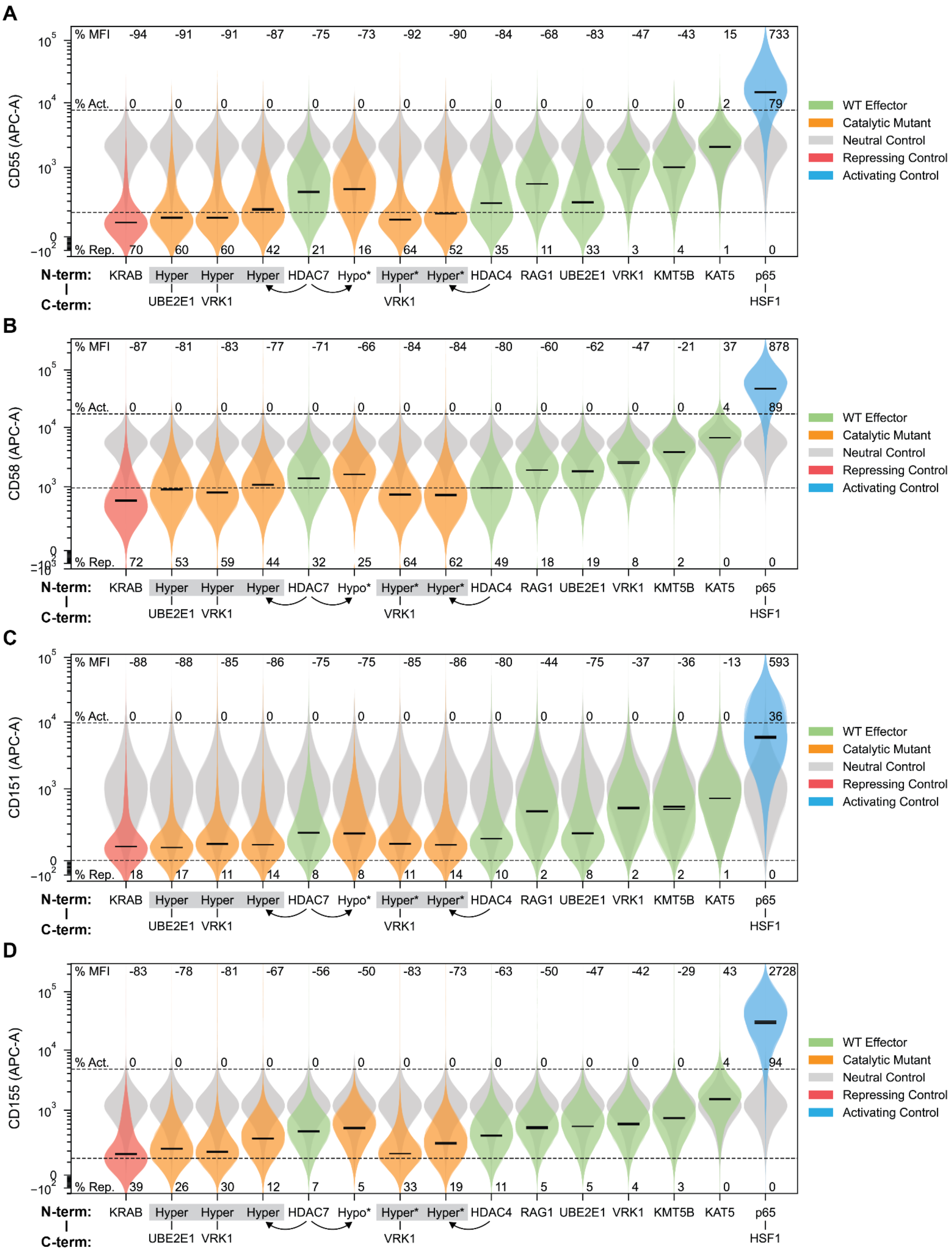
A
B
C
D
CD55 (APC-A)
CD58 (APC-A)
CD151 (APC-A)
CD155 (APC-A)
% MFI
% Act.
% Rep.
N-term:
C-term:
KRAB
Hyper
UBE2E1
VRK1
HDAC7
Hypo*
Hyper*
HDAC4
RAG1
KMT5B
KAT5
p65
HSF1
WT Effector
Catalytic Mutant
Neutral Control
Repressing Control
Activating Control

**Supplementary Figure 16. Key effectors from Library 2 transiently perturb CD55, CD58, CD151, and CD155**

(**A**) Violin plots of CD55 expression at 5 days after nucleofection and induction of key effectors identified from Library 2. WT domains are colored green. Mutants or combinations containing mutants are colored orange. Repressing and activating controls are colored red and blue respectively. MCP-DMD neutral condition is colored in gray. 3 independent replicates are illustrated as translucent overlays. The geometric means of each replicate are shown as solid black lines. Dashed lines indicate repression and activation gate at the 1st and 99th percentile of neutral condition. The average percentages of repressed and activated cells in each condition are indicated. The average percent changes in MFI versus the neutral DMD domain for each condition are indicated. *proposed mutant inferred from homology. Lack of C-terminal effector indicates that the effector was tested monovalently. Arrows point from WT effector to mutant form of the same effector. Gray boxes indicate the same effector.

(**B**) Violin plots of CD58 expression at 5 days after nucleofection and induction of key effectors identified from Library 2. Described as (**A**).

(**C**) Violin plots of CD151 expression at 5 days after nucleofection and induction of key effectors identified from Library 2. Described as (**A**).

(**D**) Violin plots of CD155 expression at 5 days after nucleofection and induction of key effectors identified from Library 2. Described as (**A**).

A

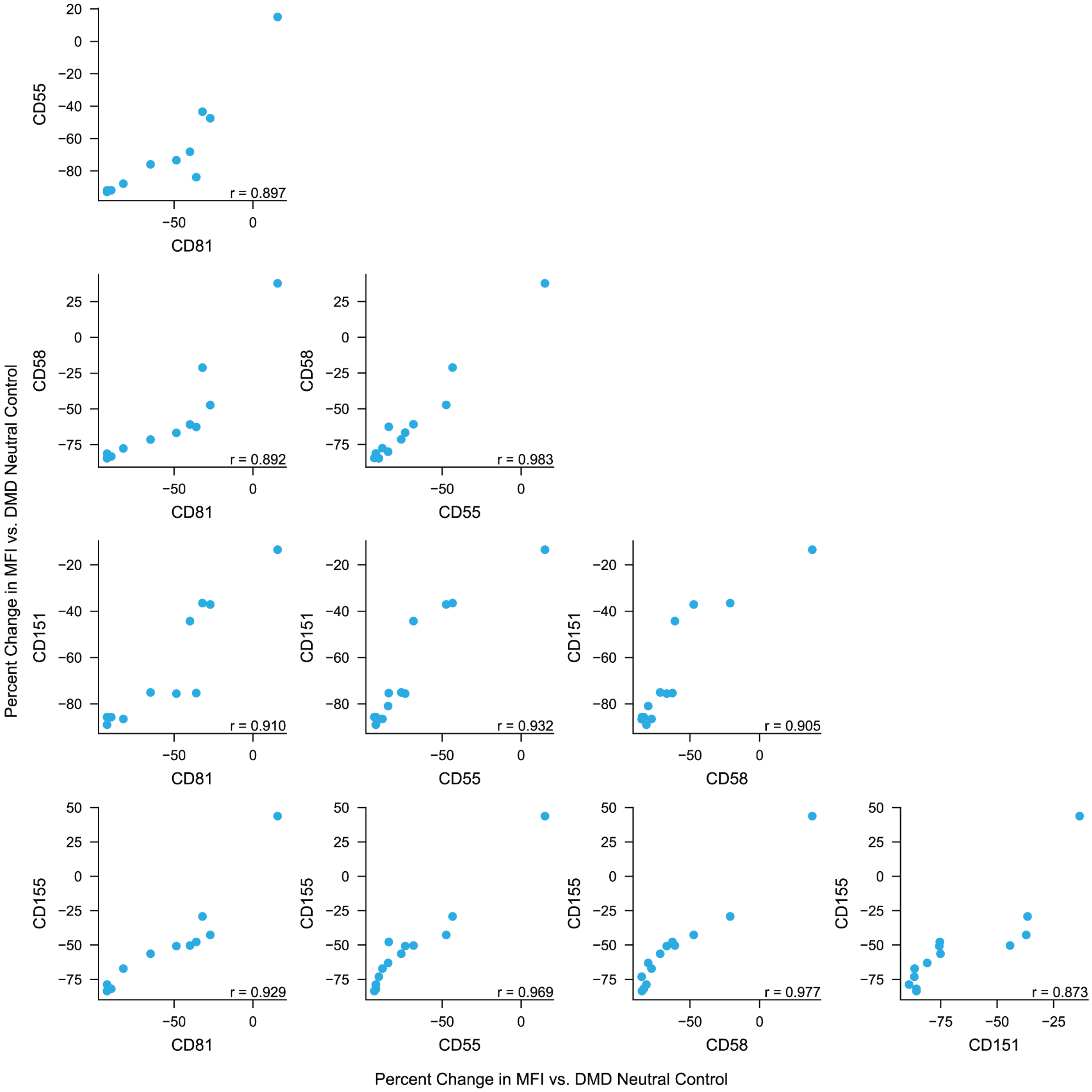


**Supplementary Figure 17. Gene-by-gene scatter plots of percent change in MFI induced by key effectors from Library 2**

(**A**) Scatter plots illustrating gene-by-gene comparisons of percent change in MFI versus the DMD neutral control for each of the 13 effectors tested in **Supplementary Fig. 16**. Pearson correlation for each comparison is indicated. KRAB and p65-HSF1 controls are not included in correlation calculation.

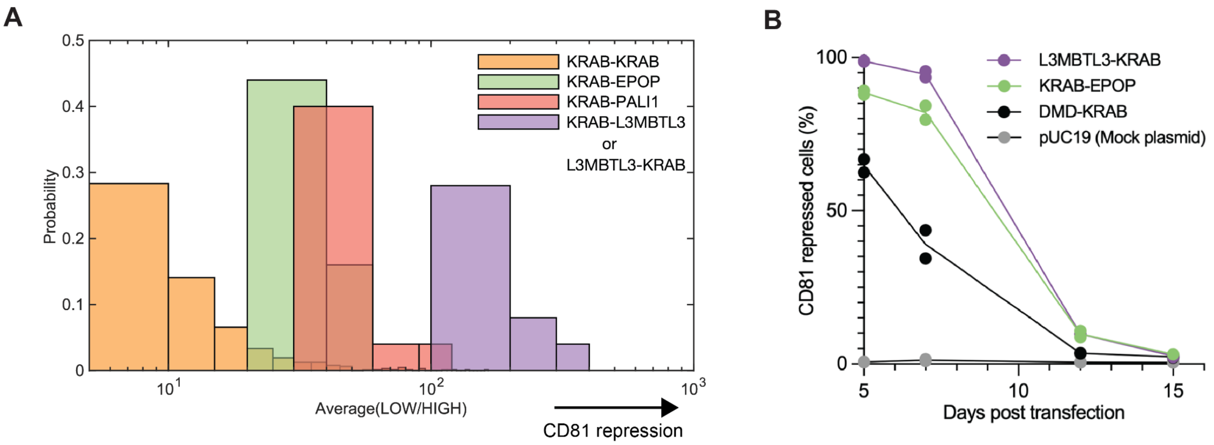


**Supplementary Figure 18. Synergistic partners of KRAB**

(**A**) Probability histograms of average HTS (Low/High) values of the selected KRAB combinations from Library 1 HTS on Day 6.

(**B**) Arrayed validation of synergistic KRAB combinations including L3MBTL3 and EPOP. KRAB domain is from the ZNF10 protein. DMD is an 80 aa fragment stuffer that has been validated to be well expressed and does not have transcriptional perturbation activity from a previous study[24].

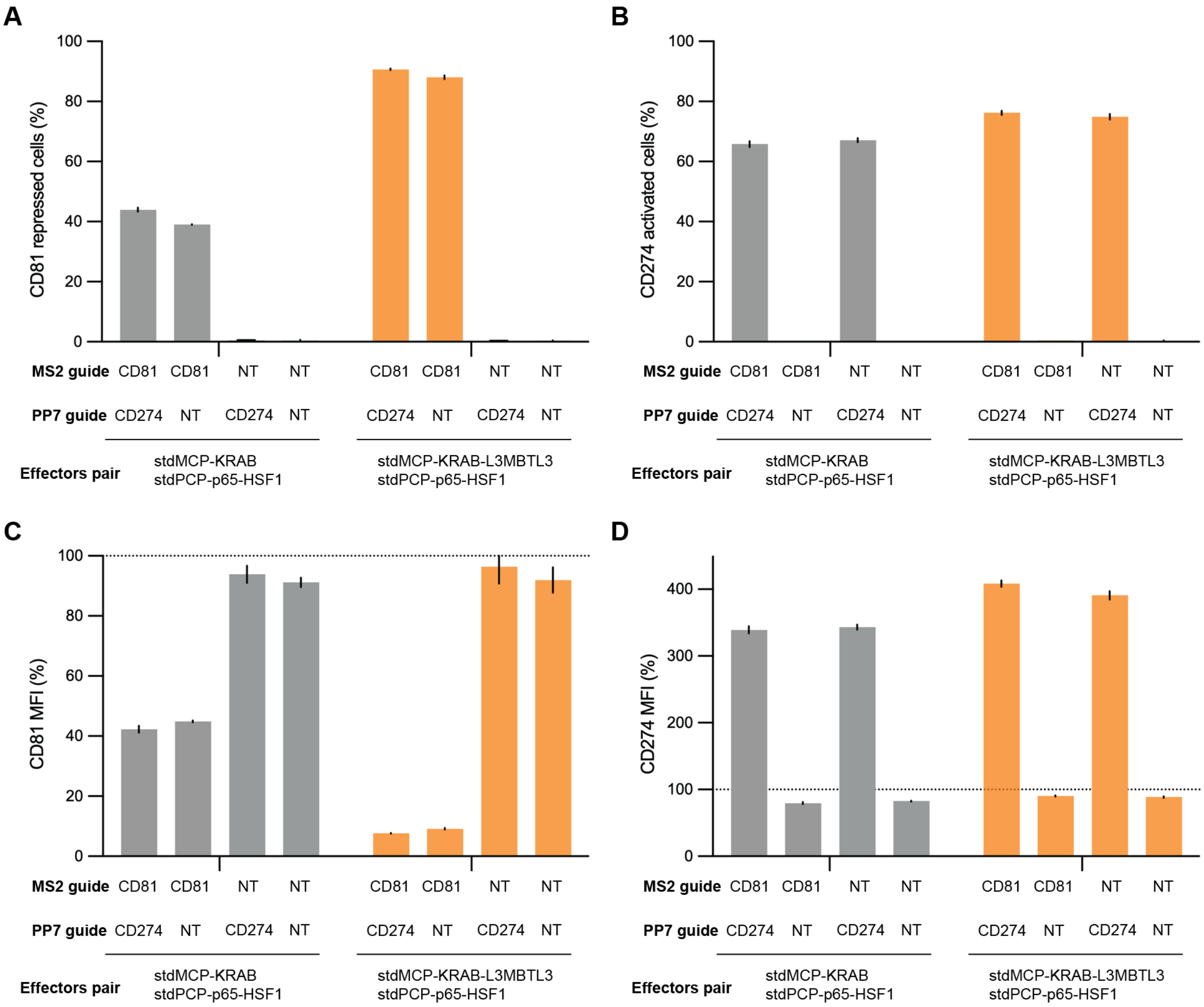


**Supplementary Figure 19. Quantification of bidirectional perturbations by MS2/PP7 system**

(**A**) Percentage of CD81 repressed cells 5 days post-infection of dual MS2/PP7 sgRNAs to the respective cell lines. Error bars indicate the standard deviation between 3 independent replicates.

(**B**) Percentage of CD274 activated cells 5 days post-infection of dual MS2/PP7 sgRNAs to the respective cell lines. Error bars indicate the standard deviation between 3 independent replicates.

(**C**) Mean fluorescence intensity (MFI) of CD81 repressed cells 5 days post-infection of dual MS2/PP7 sgRNAs to the respective cell lines. Expression level was normalized to control cells with no effectors. Error bars indicate the standard deviation between 3 independent replicates.

(**D**) Mean fluorescence intensity (MFI) of CD274 activated cells 5 days post-infection of dual MS2/PP7 sgRNAs to the respective cell lines. Expression level was normalized to control cells with no effectors. Error bars indicate the standard deviation between 3 independent replicates.

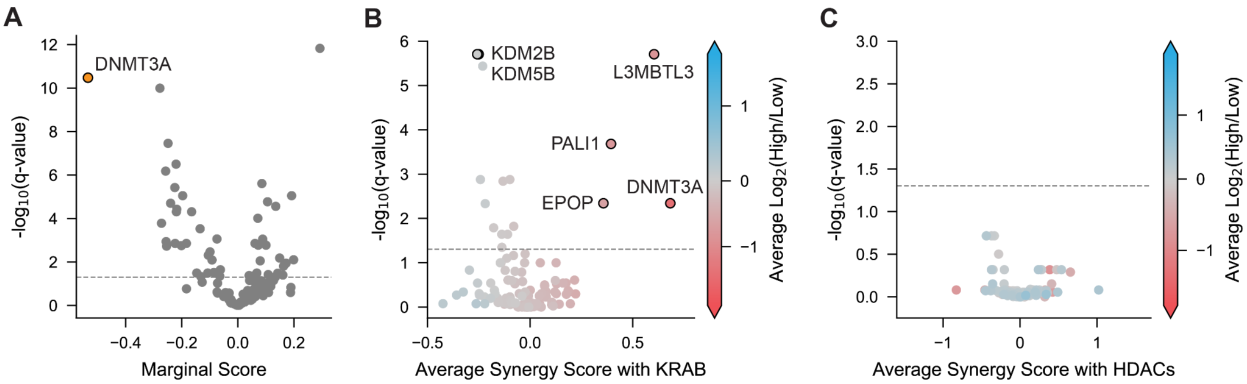


**Supplementary Figure 20. Marginal effector and synergy score analysis on Day 12**

(**A**) Volcano plot of marginal scores from Library 1 HTS on Day 12. DNMT3A with a marginal score of -0.53 is highlighted in orange. Horizontal dashed line indicates q-value of 0.05.

(**B**) Volcano plot of average synergy scores with KRAB family from Library 1 HTS on Day 12. Effectors are colored based on the average $log_2$(High/Low) enrichment scores across all combinations of the given effector with significant KRAB repressors. Q-values were calculated as FDR-corrected p-values from a one-sample Wilcoxon test. Horizontal dashed line indicates q-value of 0.05.

(**C**) Volcano plot of average synergy scores with HDAC family from Library 2 HTS on Day 12. Effectors are colored based on the average $log_2$(High/Low) enrichment scores across all combinations of the given effector with significant HDAC repressors. Q-values were calculated as FDR-corrected p-values from a one-sample Wilcoxon test. Horizontal dashed line indicates q-value of 0.05.

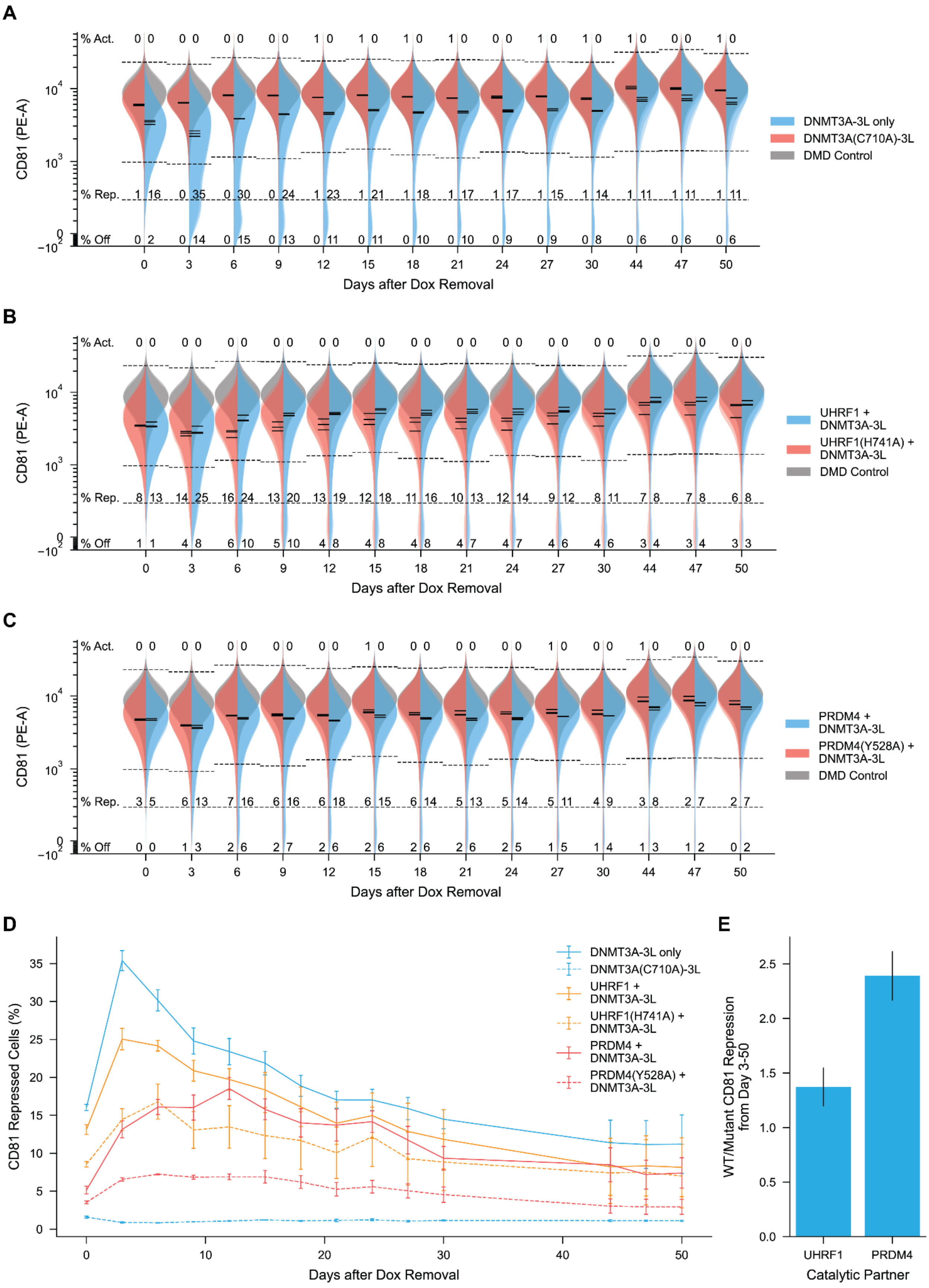

A
% Act.
% Rep.
% Off
CD81 (PE-A)
Days after Dox Removal
DNMT3A-3L only
DNMT3A(C710A)-3L
DMD Control
B
UHRF1 + DNMT3A-3L
UHRF1(H741A) + DNMT3A-3L
DMD Control
C
PRDM4 + DNMT3A-3L
PRDM4(Y528A) + DNMT3A-3L
DMD Control
D
CD81 Repressed Cells (%)
Days after Dox Removal
DNMT3A-3L only
DNMT3A(C710A)-3L
UHRF1 + DNMT3A-3L
UHRF1(H741A) + DNMT3A-3L
PRDM4 + DNMT3A-3L
PRDM4(Y528A) + DNMT3A-3L
E
WT/Mutant CD81 Repression from Day 3-50
UHRF1
PRDM4
Catalytic Partner

**Supplementary Figure 21. DNMT3A-3L combinations induce long-term partial silencing of CD81**

(**A**) Violin plot timecourse of CD81 expression following 5 days transient recruitment of DNMT3A-3L and catalytic mutant. MCP-DMD neutral condition colored in gray. 3 independent replicates are illustrated as translucent overlays. The geometric means of each replicate are shown as solid black lines. Dashed lines indicate repression, activation, and off gate. Repression and activation gates defined as 1st and 99th percentile of neutral condition. Off gate defined as 99th percentile of unstained WT K562s. The average percentage of each population in each condition is indicated.
(**B**) Violin plot timecourse of CD81 expression following 5 days transient recruitment of UHRF1 + DNMT3A-3L and catalytic mutant of UHRF1. Described as (**A**).
(**C**) Violin plot timecourse of CD81 expression following 5 days transient recruitment of PRDM4 + DNMT3A-3L and catalytic mutant of PRDM4. Described as (**A**).
(**D**) Timecourse of CD81 repressed cells following 5 days recruitment of WT combinations and mutant combinations. Error bars denote standard deviation between 3 independent replicates.
(**E**) Average fold change in CD81 repressed cells for WT effectors versus mutant effectors in DNMT3A-3L combinations. Error bars denote standard deviation between ratios calculated from average percentages across the 50 day timecourse.

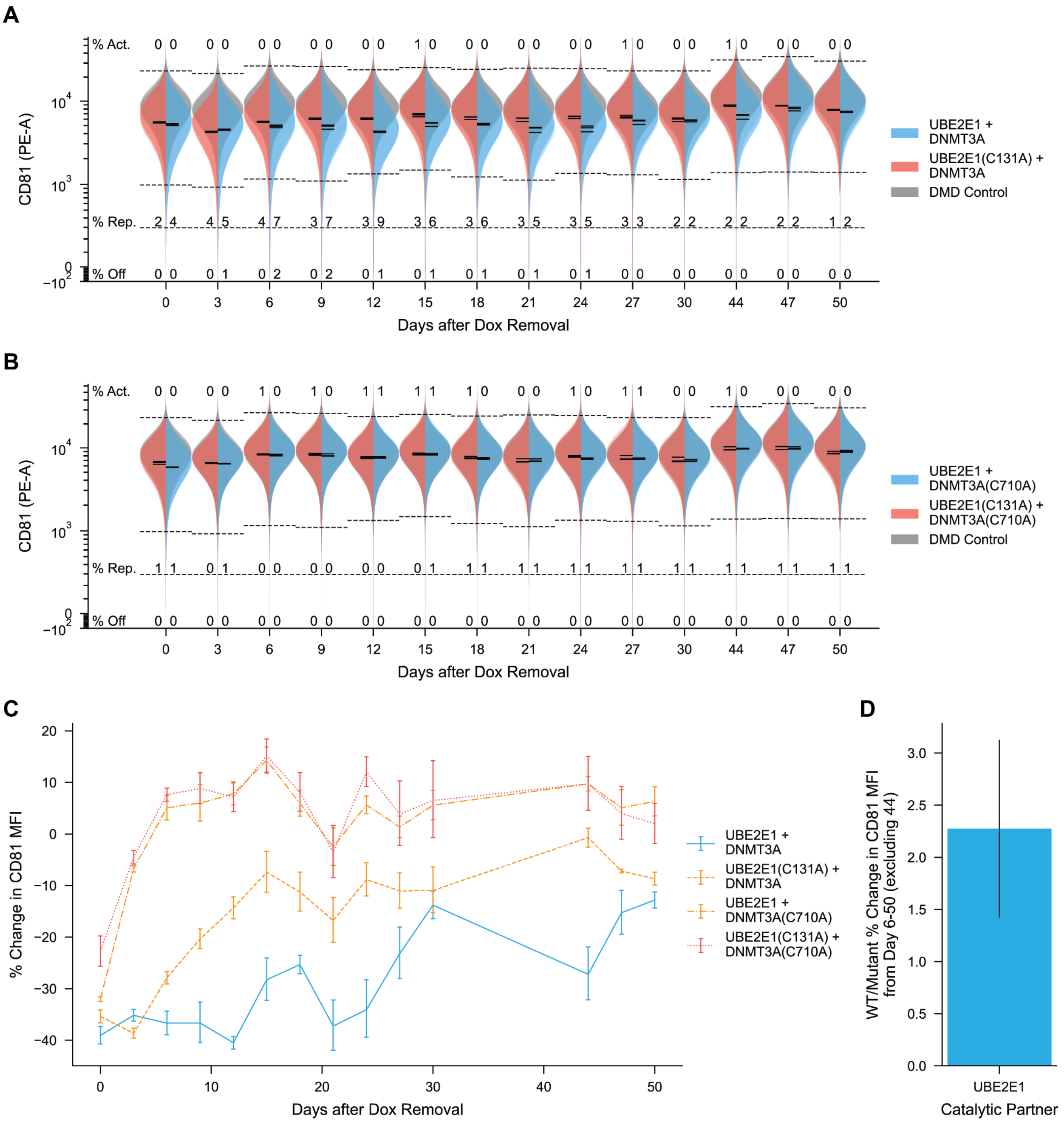


**Supplementary Figure 22. UBE2E1 + DNMT3A induces long-term repression of CD81**

(**A**) Violin plot timecourse of CD81 expression following 5 days transient recruitment of UBE2E1 + DNMT3A and UBE2E1(C131A) + DNMT3A. MCP-DMD neutral condition colored in gray. 3 independent replicates are illustrated as translucent overlays. The geometric means of each replicate are shown as solid black lines. Dashed lines indicate repression, activation, and off gate. Repression and activation gates defined as 1st and 99th percentile of neutral condition. Off gate defined as 99th percentile of unstained WT K562s. The average percentage of each population in each condition is indicated.

(**B**) Violin plot timecourse of CD81 expression following 5 days transient recruitment of UBE2E1 + DNMT3A(C710A) and UBE2E1(C131A) + DNMT3A(C710A). Described as (**A**).
(**C**) Timecourse of % change in CD81 MFI versus neutral DMD control following 5 days recruitment of UBE2E1 + DNMT3A and mutant combinations. Error bars denote standard deviation between 3 independent replicates.
(**D**) Average fold change in % change in CD81 MFI for UBE2E1 + DNMT3A versus UBE2E1(C131A) + DNMT3A. Error bars denote standard deviation between ratios calculated from average percentages across the 50 day timecourse, excluding outlier at day 44 timepoint.

A

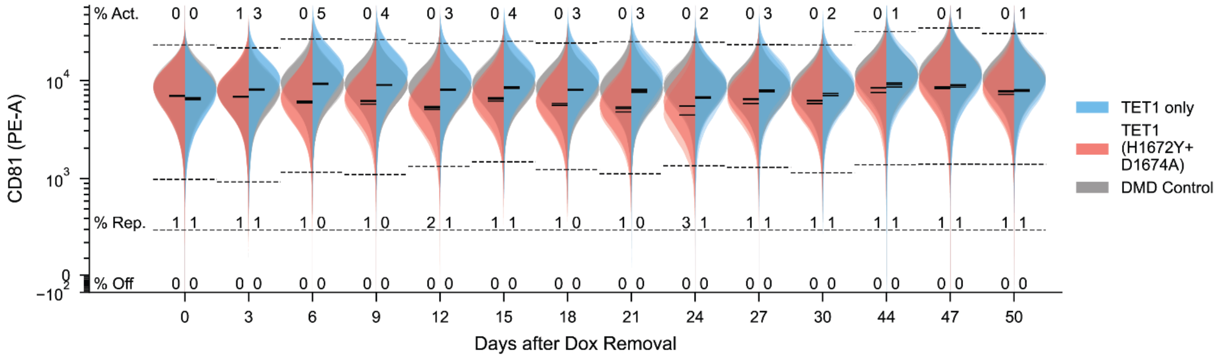
CD81 (PE-A)
% Act.
% Rep.
% Off
Days after Dox Removal
0 3 6 9 12 15 18 21 24 27 30 44 47 50
TET1 only
TET1 (H1672Y+ D1674A)
DMD Control

B

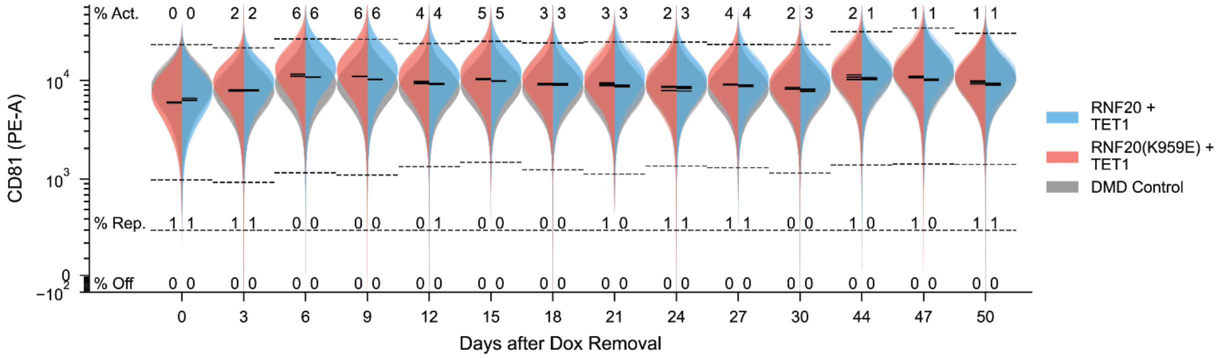
CD81 (PE-A)
% Act.
% Rep.
% Off
Days after Dox Removal
0 3 6 9 12 15 18 21 24 27 30 44 47 50
RNF20 + TET1
RNF20(K959E) + TET1
DMD Control

C

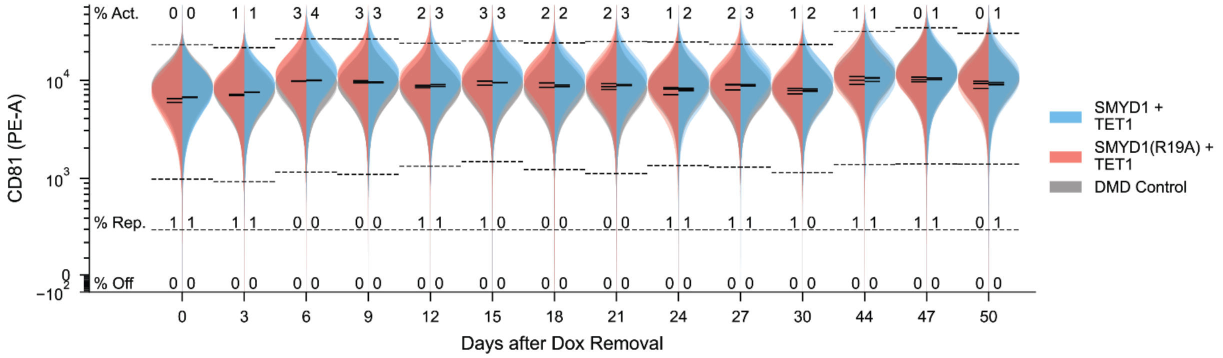
CD81 (PE-A)
% Act.
% Rep.
% Off
Days after Dox Removal
0 3 6 9 12 15 18 21 24 27 30 44 47 50
SMYD1 + TET1
SMYD1(R19A) + TET1
DMD Control

D

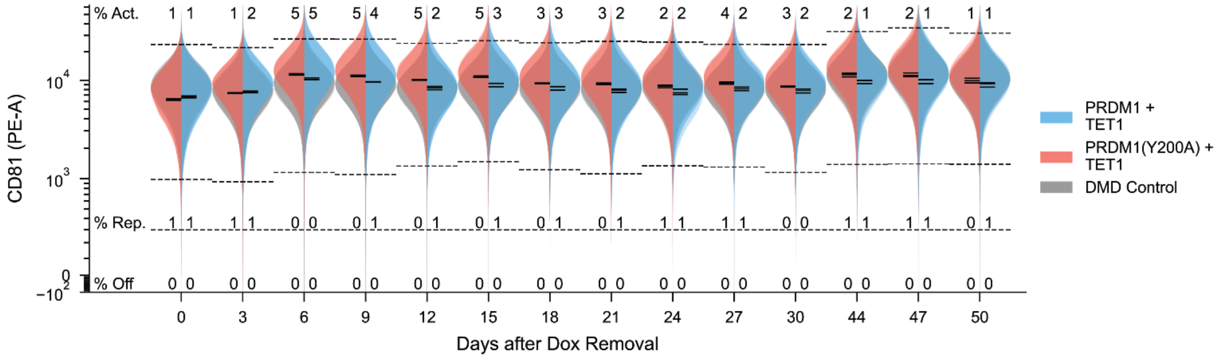
CD81 (PE-A)
% Act.
% Rep.
% Off
Days after Dox Removal
0 3 6 9 12 15 18 21 24 27 30 44 47 50
PRDM1 + TET1
PRDM1(Y200A) + TET1
DMD Control

**Supplementary Figure 23. Violin plot timecourses of TET1 combinations**

(**A**) Violin plot timecourse of CD81 expression following 5 days transient recruitment of TET1 and TET1(H1672Y+D1674A). MCP-DMD neutral condition colored in gray. 3 independent replicates are illustrated as translucent overlays. The geometric means of each replicate are shown as solid black lines. Dashed lines indicate repression, activation, and off gate. Repression and activation gates defined as 1st and 99th percentile of neutral condition. Off gate defined as 99th percentile of unstained WT K562s. The average percentage of each population in each condition is indicated.

(**B**) Violin plot timecourse of CD81 expression following 5 days transient recruitment of RNF20 + TET1 and RNF20(K959E) + TET1. Described as (**A**).

(**C**) Violin plot timecourse of CD81 expression following 5 days transient recruitment of SMYD1 + TET1 and SMYD1(R19A) + TET1. Described as (**A**).

(**D**) Violin plot timecourse of CD81 expression following 5 days transient recruitment of PRDM1 + TET1 and PRDM1(Y200A) + TET1. Described as (**A**).

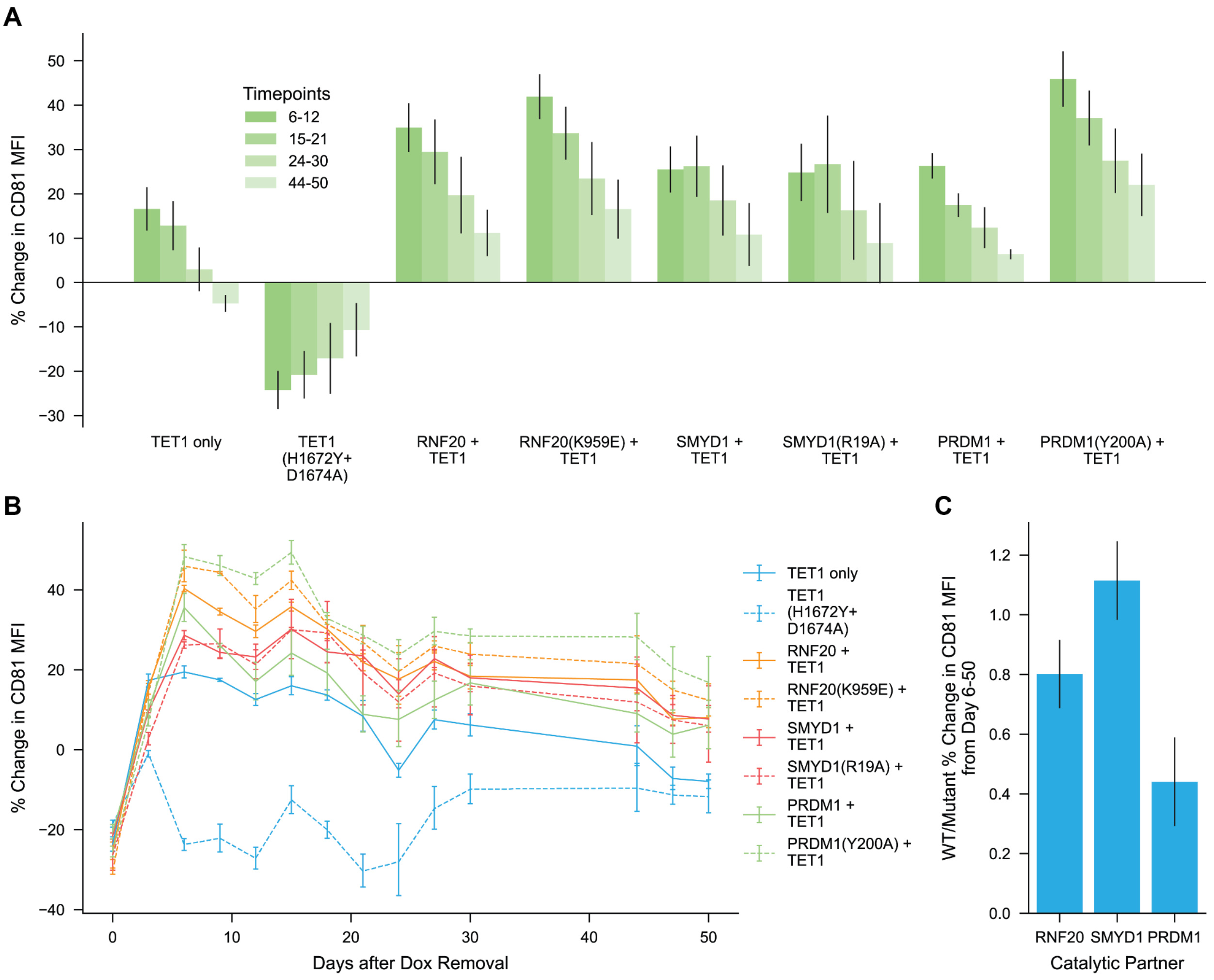


**Supplementary Figure 24. TET1 combinations induce long-term activation of CD81**

(**A**) Timecourse of percent change in CD81 MFI versus neutral DMD control following nucleofection of MCP-TET1 combination encoding plasmids and 5 days recruitment of dCas9/MS2 complex. Day 0 indicates the day of doxycycline washout. Each bar represents 3 timepoints collected within the specified days. The bar heights represent the average perturbation across the timepoints, and the error bars represent the standard deviation between 3 independent replicates.

(**B**) Timecourse of % change in CD81 MFI versus neutral DMD control following 5 days recruitment of TET1 combinations and mutants. Error bars denote standard deviation between 3 independent replicates.

(**C**) Average fold change in % change in CD81 MFI for WT effectors versus mutant effectors in TET1 combinations. Error bars denote standard deviation between ratios calculated from average percentages across the 50 day timecourse.

### Supplementary Note 1

#### Synergistic interaction between the H2AK119ub1 reader and a subset of PRC1 recruiters

Recognition of epigenetic modifications by a reader domain is a crucial step in establishing epigenetic feedback loops, representing a fundamental aspect of epigenetic interactions. Specifically, PRC1 and PRC2, responsible for installing H2AK119ub1 and H3K27me3, respectively, are well-known for their intricate feedback mechanisms. These mechanisms involve the spreading of a single epigenetic modification or cross-talk between two distinct modifications[4], all of which require the recognition of epigenetic modifications by the reader domain[6]. By focusing on the sub-heatmap defined by recruiters of PRC1 and PRC2 on the N-terminus and readers of epigenetic modifications on the C-terminus, we were able to notice a prominent signal that emerged at the intersection of a subset of PRC1 recruiters and the H2AK119ub1 reader (**Supplementary Fig. 13a**). Out of 483 combinations within the sub-heatmap, 5 combinations out of the top 8 most repressive combinations were from combinations between PRC1 recruiters and the H2AK119 monoubiquitination (H2AK119ub1) reader domain from RYBP protein. Also, the H2AK119ub1 reader domain was nominated as the strongest synergistic C-terminus partner of PRC1 recruiters excluding the KRAB family, although the q-value was higher than the threshold of 0.05 (**Supplementary Fig. 13b**).

To validate these combinations between PRC1 recruiters and the H2AK119ub1 reader, we tested several PRC1 recruiter domains encompassing domains from canonical PRC1 (CBX2/CBX7) and variant PRC1 (RYBP), in combination with the readers of distinct histone modifications (H2AK119ub1 by the RYBP Ranbp2-ZnF domain, H3K27me by the CBX6 Chromo domain, H3K9me by the CBX5 Chromo domain, and H3K4me by the ING1 PHD domain) (**Supplementary Fig. 13c**). After 4 days of recruitment, our results confirmed that only the combinations containing the H2AK119ub1 reader domain from RYBP were sufficient to induce substantial gene silencing (CBX2 + H2AK119ub1 reader: 12.7%, CBX7 + H2AK119ub1 reader: 59.7%, RYBP + H2AK119ub1 reader: 60.5% CD81 repressed cells), irrespective of whether the PRC1 component originated from the canonical or variant complex (**Supplementary Fig. 13d**). Interestingly, the validated fusion protein containing the PRC1 recruiter domain and H2AK119ub1 reader domain from RYBP essentially reconstitutes full-length RYBP with some truncations (**Supplementary Fig. 13e**). Variant PRC1 (vPRC1) comprising RYBP subunits has the unique ability to both read and write the H2AK119ub1 mark owing to the H2AK119ub1 reader domains found within RYBP protein[37]. This ability to read and write the same epigenetic modifications has been speculated as an efficient strategy for propagating silencing marks to adjacent regions[96], suggesting that interactions detected from our HTS experiment may recapitulate a natural architecture inherent within the PRC1 complex[37].
